# Co-utilization of microalgae and heterotrophic microorganisms improves wastewater treatment efficiency

**DOI:** 10.1101/2024.06.05.597533

**Authors:** Miiku Takahashi, Yukino Karitani, Ryosuke Yamada, Takuya Matsumoto, Hiroyasu Ogino

**Author notes:** Corresponding author: Ryosuke Yamada E-mail address (R. Yamada).

## Abstract

Wastewater treatment using co-culture systems of microalgae and heterotrophic microorganisms is expected to be useful under atmospheric dilute carbon dioxide conditions. In this study, we investigated the combination of microalgae and heterotrophic microorganisms to improve the efficiency of wastewater treatment. Furthermore, to elucidate the cause of the changes in wastewater treatment efficiency in the co-culture system, changes in gene expression were revealed through transcriptome analysis. Three types of microalgae and five heterotrophic microorganisms were used in combination for wastewater treatment. The combination of *Chlamydomonas reinhardtii* NIES-2238 and *Saccharomyces cerevisiae* SH-4 showed the highest wastewater treatment efficiency. Using this combination for artificial wastewater treatment, the removal rates of TOC (Total organic carbon), PO4^3-^, and NH_4_^+^ reached 80%, 93%, and 63%, respectively, after 18 h of treatment. Transcriptome analysis revealed that the combined wastewater treatment altered the expression of 1371 and 692 genes in *C. reinhardtii* and *S. cerevisiae*, respectively. The genes upregulated in *C. reinhardtii* included those related to molecular and ion transport. Genes upregulated in *S. cerevisiae* included those related to cell protection from various types of damage and stress. To the best of our knowledge, this is the first study to show that a combination of green algae and yeast improves the efficiency of wastewater treatment. As both the green alga *C. reinhardtii* and the yeast *S. cerevisiae* are highly safe microorganisms, the establishment of their effective combination for wastewater treatment is highly significant.

## 1. Introduction

Daily household and industrial activities discharge water containing carbon and nutrient compounds, such as nitrogen and phosphorus. These compounds are the main causes of eutrophication in oceans and lakes (Dodds et al., 2016; Nguyen et al., 2020). Eutrophication disrupts the balance of the water ecosystem and leads to environmental pollution; therefore, it is necessary to remove nutrient compounds to avoid eutrophication (Nguyen et al., 2020). In general wastewater treatment, the activated sludge method is used in which organic matter is decomposed by microorganisms (Gernaey et al., 2004; Mujtaba et al., 2017; Wilén et al., 2018). However, the decomposition of organic matter using the activated sludge method requires a large amount of oxygen, which means that a large amount of electricity is required for aeration. The energy for aeration accounts for 60–80% of the total energy used in wastewater treatment using the activated sludge method (Chauchuat et al., 2005). Furthermore, because the activated sludge method has low nitrogen and phosphorus removal efficiency, additional treatment is often required.

Wastewater treatment with microalgae has attracted attention as an energy-saving alternative to activated sludge treatment (Abdel-Raouf et al., 2012; Mujtaba et al., 2017; Wang et al., 2016). As microalgae perform photosynthesis, they do not require aeration and are expected to fix dilute CO_2_ in the atmosphere. In addition, they are characterized by high nitrogen and phosphorus removal capacities compared to common microorganisms. In a previous study, the green alga *Chlamydomonas reinhardtii* was used for wastewater treatment in a palm oil mill with removal rates of 100% for ammonium and nitrate, 66–89% for phosphate, and 16.7–33.2% for chemical oxygen demand (COD) (Mohd et al., 2024). In another report, more than 80% of the phosphorus and nitrogen in wastewater were removed when wastewater was treated with the green algae *Chlorella vulgaris* and *Scenedesmus sp.* (Tam et al., 1990). In addition, when wastewater was treated with the cyanobacterium *Arthrospira platensis*, more than 98% of the COD, phosphorus, and nitrogen were removed within 4–5 days of treatment (Hena et al., 2018).

One problem with photosynthetic microalgae is that their growth rate is lower than that of common microorganisms in an atmosphere of dilute CO_2_ (Chinnasamy et al., 2009). In microalgae cultivation, high-concentration carbon dioxide aeration is performed to increase the growth ability of the microalgae. Therefore, the co-cultivation of microalgae and heterotrophic microorganisms is attracting attention to improve the growth of microalgae without the aeration of high-concentration carbon dioxide (Naseema Rasheed et al., 2023; Ray et al., 2022). Previous studies reported that the growth rate of microalgae increases under dilute CO_2_ atmospheric conditions when *C. reinhardtii* and *Saccharomyces cerevisiae* are co-cultured (Karitani et al., 2024a; Karitani et al., 2024b). Co-cultivation of *C. reinhardtii* and *Escherichia coli* promotes microalgal growth (Yamada et al., 2023). These coculture systems of microalgae and heterotrophic microorganisms are expected to increase the growth rate of microalgae, even in wastewater treatment, and improve the efficiency of wastewater treatment using microalgae.

In this study, we investigated different combinations of microalgae and heterotrophic microorganisms to improve the efficiency of wastewater treatment. Specifically, three types of microalgae and five types of heterotrophic microorganisms were used in combinations for wastewater treatment, and the optimal combination for wastewater treatment was determined. Subsequently, wastewater containing higher concentrations of carbon, nitrogen, and phosphorus was treated with the optimal combination of microorganisms, and the removal efficiency was evaluated. Furthermore, to elucidate the cause of the changes in wastewater treatment efficiency with the optimal combination, changes in gene expression were assessed through transcriptome analysis.

## 2. Results

### 2.1. Optimal combination of microalgae and heterotrophic microorganisms for wastewater treatment

Microalgae and heterotrophic microorganisms were combined for 1x artificial wastewater treatment to determine optimal combinations. Fig. 1 shows the removal rates of TOC (Total organic carbon), PO_4_^3-^, and NH_4_^+^ in wastewater treatment using microalgae and heterotrophic microorganisms alone and in combination.

**Fig. 1.**
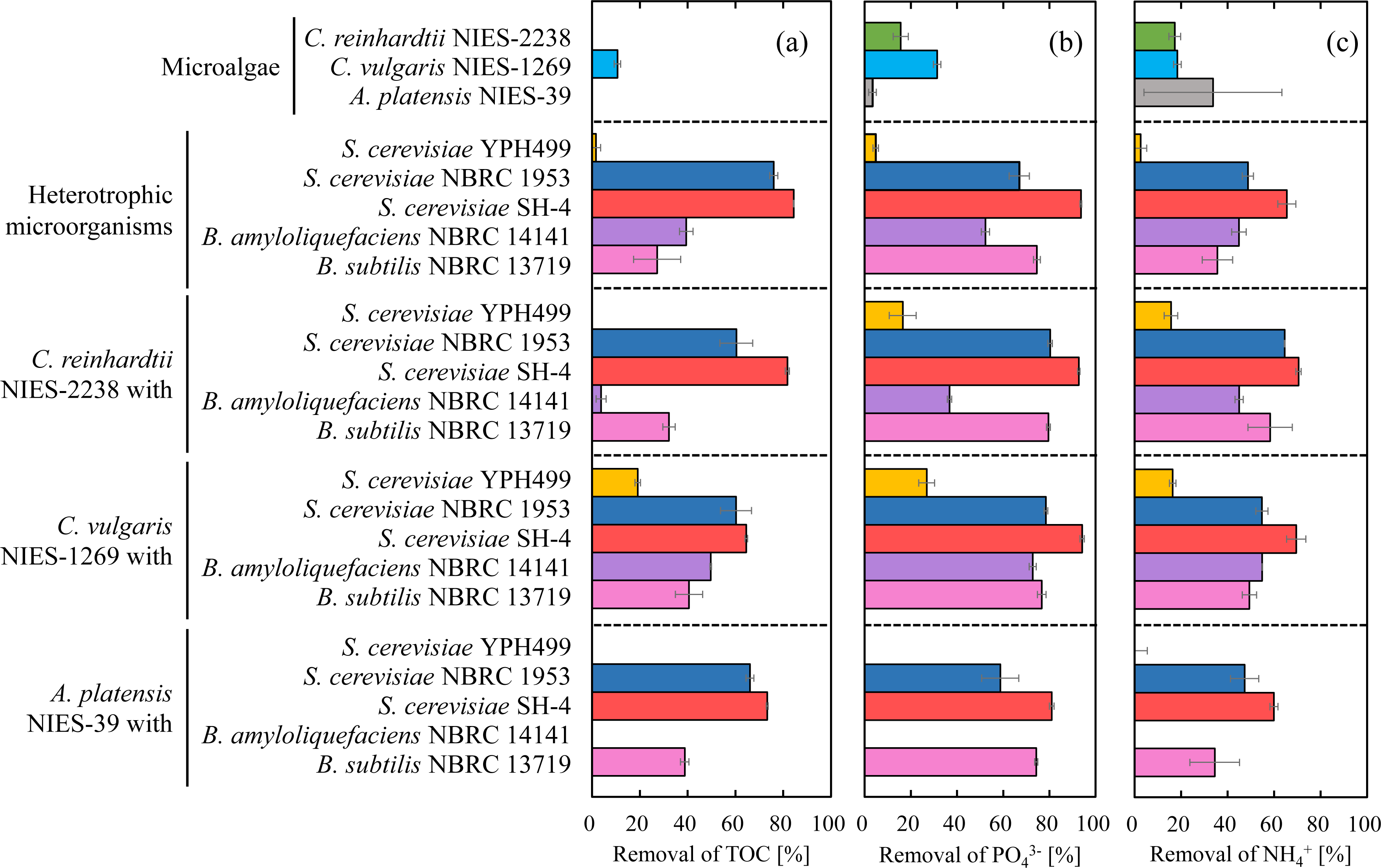
Removal rates of (a) TOC, (b) PO_4_^3-^, and (c) NH_4_^+^ from 1x artificial wastewater using microalgae and heterotrophic microorganisms alone and in combination. Data represent the averages of two replicates, and error bars indicate standard deviation.

The TOC removal rate was 11% for *C. vulgaris* NIES-1269, which was the highest among the three microalgae when treated alone (Fig. 1a). When only heterotrophic microorganisms were used, *S. cerevisiae* SH-4 exhibited the highest removal rate (84 %) among the five heterotrophic microorganisms. When the three types of microalgae and *S. cerevisiae* YPH499 were used in combination for wastewater treatment, the TOC removal rates remained low, whereas when the three types of microalgae and *S. cerevisiae* SH-4 were used in combination, the TOC removal rates were relatively high. Among the 15 microbial combinations, the combination of *C. reinhardtii* NIES-2238 and *S. cerevisiae* SH-4 resulted in the highest TOC removal rate (82 %).

*C. vulgaris* NIES-1269 exhibited the highest PO_4_ ^3-^ removal rate (31%) among the three microalgae when the wastewater was treated with microalgae alone (Fig. 1b). When heterotrophic microorganisms were used alone, *S. cerevisiae* SH-4 showed the highest removal rate (94 %) among the five microorganisms. Similar to the TOC removal rate, when the three types of microalgae and *S. cerevisiae* YPH499 were used in combination for wastewater treatment, the PO_4_^3-^ removal rates remained low, whereas when the three types of microalgae and *S. cerevisiae* SH-4 were used in combination, PO ^3-^ removal rates were relatively high. Among the 15 microbial combinations, the PO_4_^3-^ removal rate was the highest when *C. reinhardtii* NIES-2238 and *C. vulgaris* NIES-1269 were combined with *S. cerevisiae* SH-4, with 93% and 94%, respectively.

*A. platensis* NIES-39 exhibited the highest NH_4_^+^ removal rate (34%) among the three microalgae when wastewater was treated with microalgae alone (Fig. 1c). When only heterotrophic microorganisms were used, *S. cerevisiae* SH-4 exhibited the highest removal rate (66 %) among the five heterotrophic microorganisms. Similar to the TOC removal rate, when the three types of microalgae and *S. cerevisiae* YPH499 were used in combination for wastewater treatment, the NH_4_^+^ removal rates remained low. When the three types of microalgae and *S. cerevisiae* SH-4 were used in combination, the NH_4_^+^ removal rates were relatively high. Among the 15 microbial combinations, the highest NH_4_^+^ removal rate was achieved when *C. reinhardtii* NIES-2238 and *C. vulgaris* NIES-1269 were used in combination with *S. cerevisiae* SH-4 (71% and 70 %, respectively).

### 2.2. Wastewater treatment by optimal combination

Among 15 microbial combinations, the combination of *C. reinhardtii* NIES-2238 and *S. cerevisiae* SH-4 was optimal for wastewater treatment (Fig. 1). With this combination, wastewater treatment was conducted for 18 h using 2x artificial wastewater with increased concentrations of TOC, PO_4_^3-^ and NH_4_^+^ to evaluate the wastewater treatment performance (Fig. 2).

**Fig. 2.**
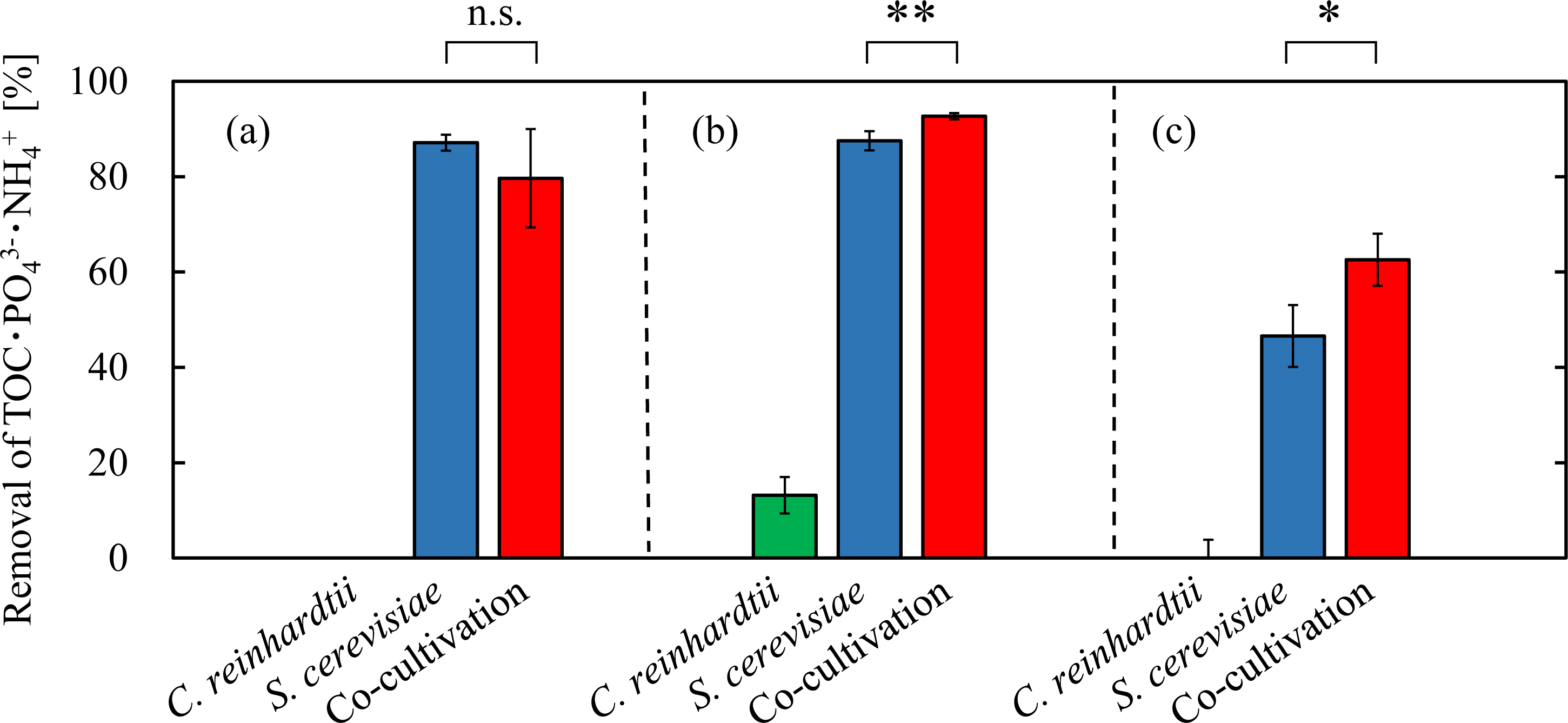
Removal rates of (a) TOC, (b) PO_4_^3-^, and (c) NH_4_^+^ from 2x artificial wastewater with *Chlamydomonas reinhardtii* NIES-2238 and *Saccharomyces cerevisiae* SH-4 alone and in combination. Data represent the average of three replicates, and error bars indicate standard deviation. Significant differences were calculated using Student’s t-test (*; 0.05 < p < 0.10, ** p < 0.05). n.s.; not significant.

A similarly high TOC removal rate of over 80% was achieved when *S. cerevisiae* SH-4 was used alone, and when *C. reinhardtii* NIES-2238 and *S. cerevisiae* SH-4 were used in combination (Fig. 2a). The combination of *C. reinhardtii* NIES-2238 and *S. cerevisiae* SH-4 achieved the highest PO_4_^3-^ removal rate of 93% (Fig. 2b). This was 7.0 times higher than that achieved with *C. reinhardtii* NIES-2238 alone (13%) and 1.1 times higher than that achieved with *S. cerevisiae* SH-4 alone (88%). The combination of *C. reinhardtii* NIES-2238 and *S. cerevisiae* SH-4 achieved the highest NH_4_^+^ removal rate of 63%, which was 1.3 times higher than that achieved with *S. cerevisiae* SH-4 alone (47%) (Fig. 2c). Therefore, the removal rate of PO_4_^3-^ and NH_4_^+^ from 2x artificial wastewater was significantly increased using *C. reinhardtii* NIES-2238 and *S. cerevisiae* SH-4 in combination for wastewater treatment.

### 2.3. Comparison of gene expression in wastewater treatment alone and in combination

After 18 h of wastewater treatment with *C. reinhardtii* NIES-2238 or *S. cerevisiae* SH-4 alone or in combination, total RNA was extracted from each cell and subjected to transcriptome analysis. Volcano plots showing p-values and fold-changes for each gene of *C. reinhardtii* NIES-2238 and *S. cerevisiae* SH-4 obtained by transcriptome analysis are shown in Fig. 3.

**Fig. 3.**
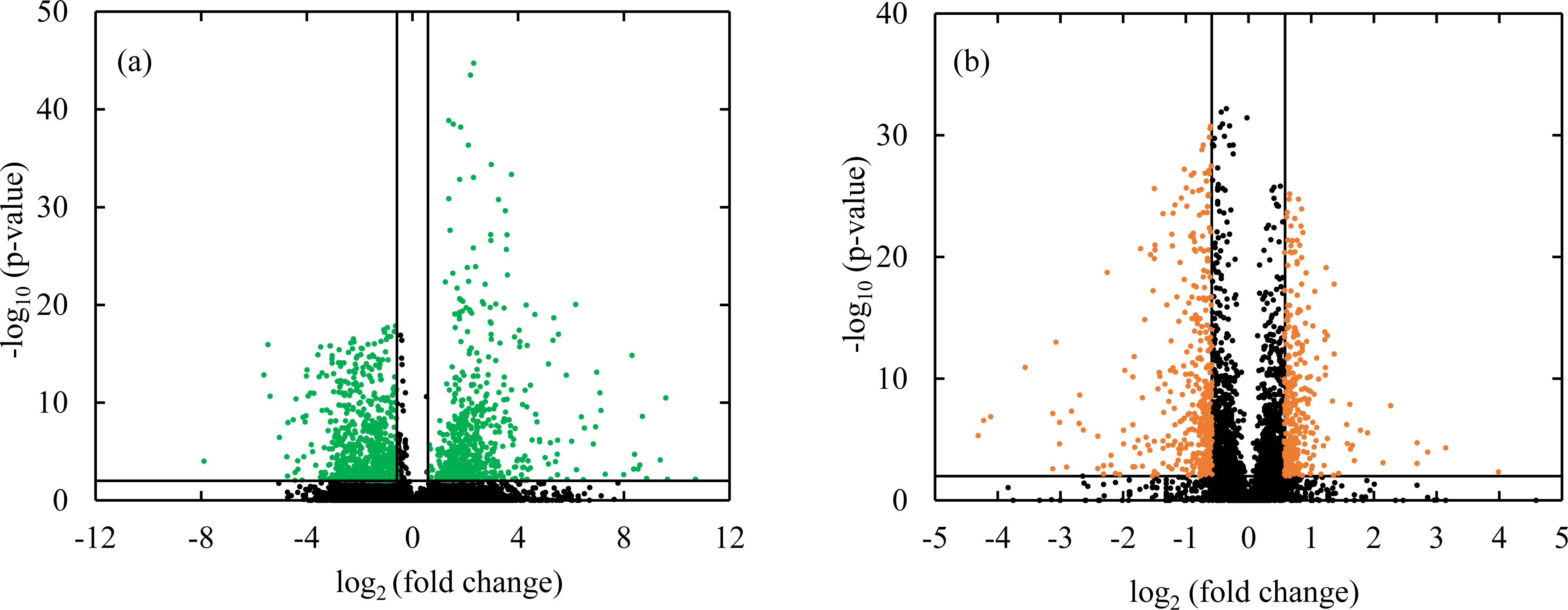
Volcano plot of genes for (a) *Chlamydomonas reinhardtii* NIES-2238 and (b) *Saccharomyces cerevisiae* SH-4.

In *C. reinhardtii* NIES-2238, the expression of 1371 genes, 636 with upregulated and 735 with downregulated expression, was altered by the combined wastewater treatment (Fig. 3a). In contrast, as shown in Fig. 3b, in *S. cerevisiae* SH-4, the expression of 692 genes, 305 with upregulated and 387 with downregulated expression, was altered by the combined wastewater treatment.

Gene ontology (GO) analysis was performed on the upregulated and downregulated genes in *C. reinhardtii* NIES-2238 and *S. cerevisiae* SH-4 to identify gene functions that were significantly more frequent among the differentially expressed genes (DEGs). The 20 GO terms with the highest log10 (p-value) values obtained by GO analysis are shown in Fig. 4 (for *C. reinhardtii* NIES-2238) and Fig. 5 (for *S. cerevisiae* SH-4).

**Fig. 4.**
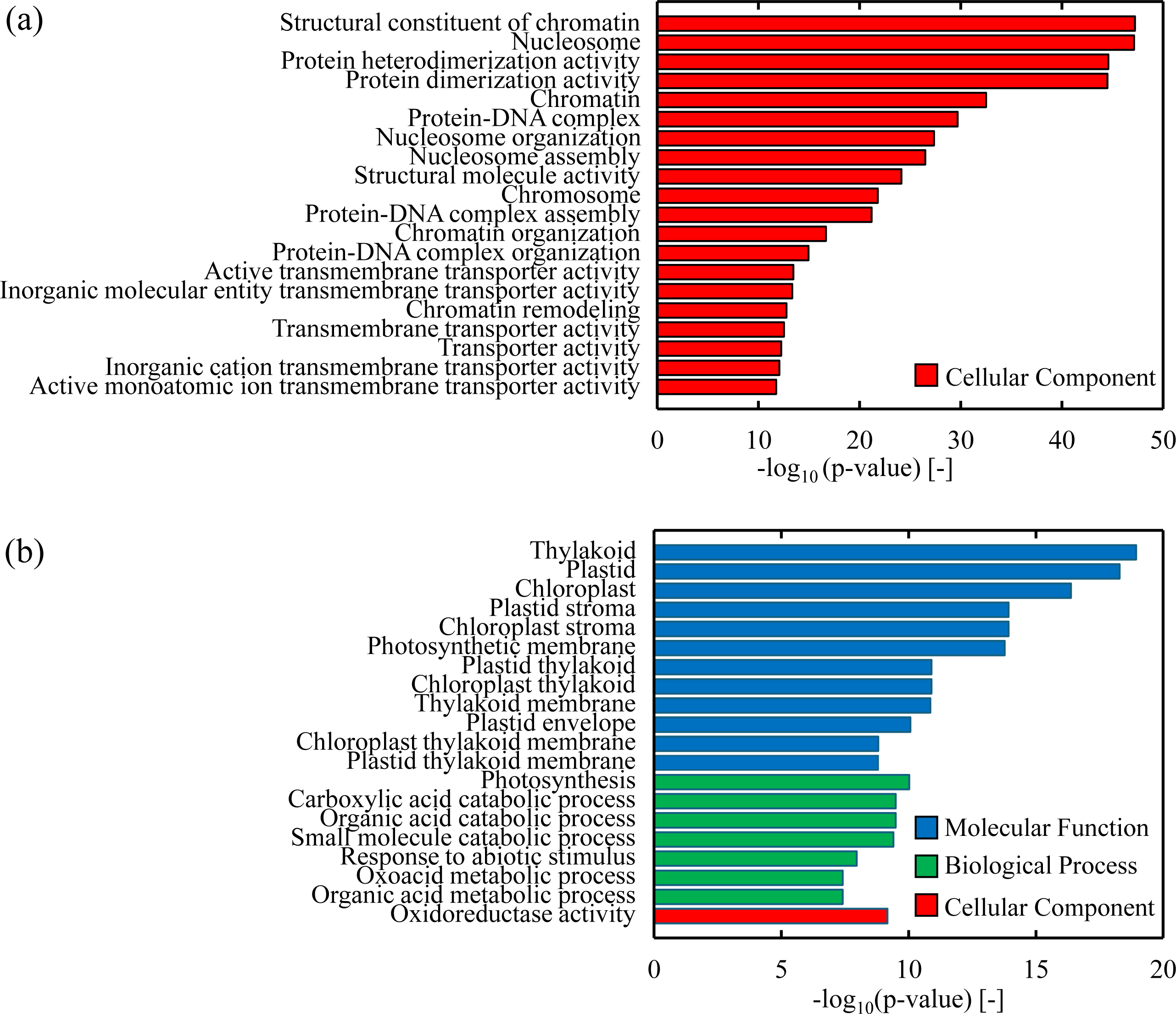
Top 20 GO terms associated with *Chlamydomonas reinhardtii* NIES-2238 genes whose expression was (a) increased and (b) decreased via co-cultivation.

**Fig. 5.**
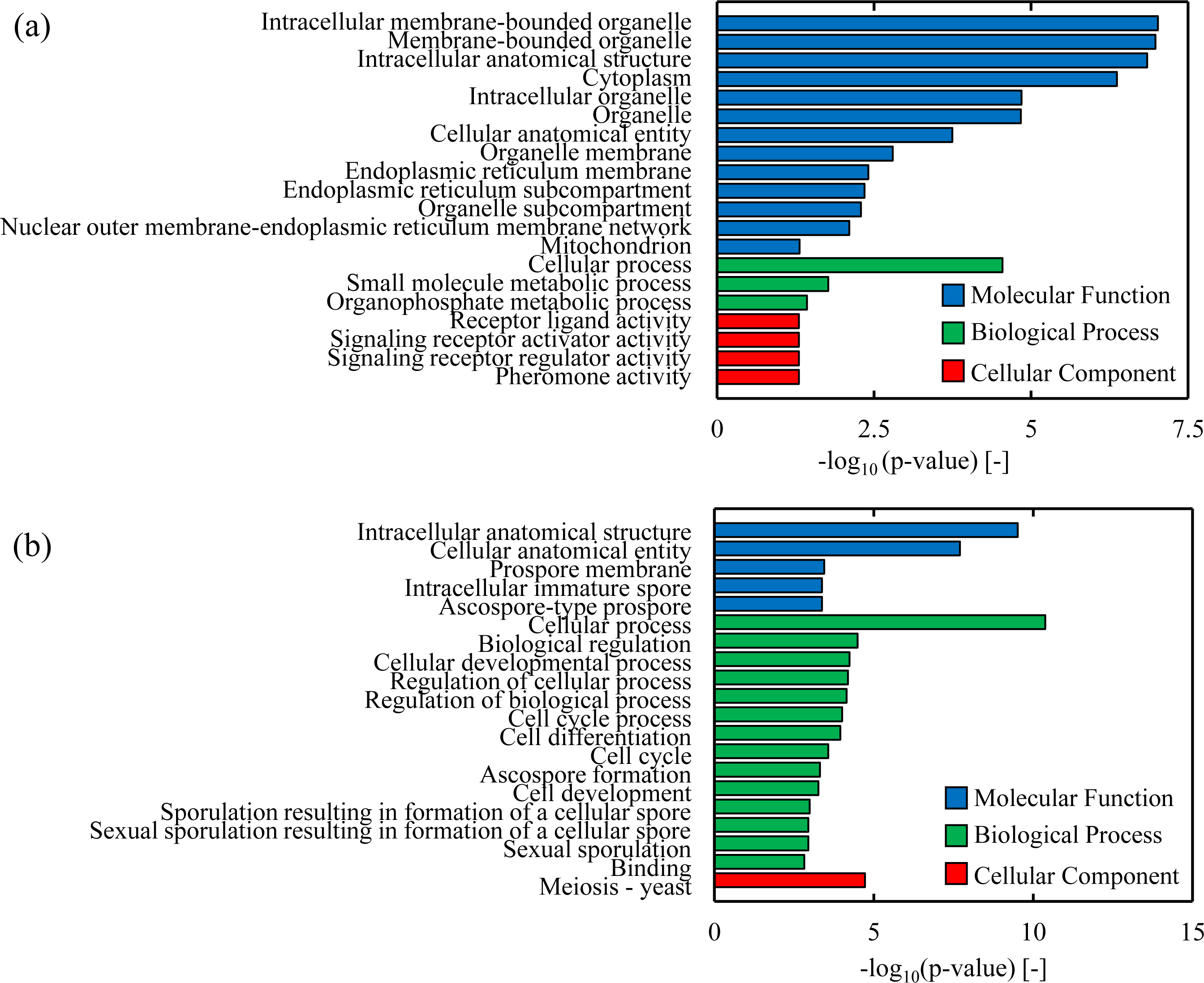
Top 20 GO terms of *Saccharomyces cerevisiae* SH-4 genes whose expression was (a) increased and (b) decreased via co-cultivation.

The GO terms for the upregulated genes in *C. reinhardtii* NIES-2238 were mainly associated with functions related to chromatin components and nucleosomes (Fig. 4a). They were also associated with functions related mainly to the transport of molecules and ions, such as transporter activity and inorganic molecular body transmembrane transporter activity. Some GO terms for the genes downregulated in *C. reinhardtii* NIES-2238 were related to functions of thylakoids, plastids, chloroplasts, and photosynthetic membranes (Fig. 4b).

Some GO terms for the genes upregulated in *S. cerevisiae* SH-4 were associated with functions related to membrane-bound organelles, cellular processes, and signal receptor activity (Fig. 5a). Some GO terms of the genes downregulated in *S. cerevisiae* SH-4 were associated with functions such as cellular processes, intracellular anatomical structure, and cellular anatomical entities (Fig. 5b).

## 3. Discussion

The objective of this study is to clarify the optimal combination of microalgae and heterotrophic microorganisms to improve the efficiency of wastewater treatment by combining microalgae and heterotrophic microorganisms. When *C. reinhardtii* NIES-2238 and *S. cerevisiae* SH-4 were combined for wastewater treatment, TOC decreased from 424 mg/L to 86 mg/L (equivalent to approximately 1275 mg/L and 264 mg/L of COD, respectively [Dubber et al., 2010]) with an 80% removal rate, while total phosphorus decreased from 12 mg/L to 0.90 mg/L with a 93% removal rate of PO_4_^3-^, and total nitrogen decreased from 39 mg/L to 14 mg/L with a 63% removal rate of NH_4_^+^ in 18 h (Fig. 2). Similar to the present study, a previous study treating wastewater containing similar concentrations of carbon, phosphorus, and nitrogen showed that combined wastewater treatment with the microalgae *C. vulgaris* and the fungal *Aspergillus* sp. in concentrated wastewater decreased COD from 1660 mg/L to 622 mg/L with a 63% removal rate, total phosphorus from 52.6 mg/L to 5.35 mg/L with a 90% removal rate, and total nitrogen from 97.2 mg/L to 40.0 mg/L with a 59% removal rate after 24 h. (Zhou et al., 2012). In another study, wastewater treatment using a combination of microalgae (*C. vulgaris*) and a fungus *(Ganoderma lucidum*) resulted in COD, total phosphorus, and total nitrogen removal rates of 80%, 85%, and 74%, respectively (Guo et al., 2017). Therefore, the wastewater treatment efficiency in this study using safer microorganisms was comparable to or higher than that of previous studies. Interestingly, in the present study, the removal rates of TOC, PO_4_^3-^, and NH_4_^+^ were similar whether 1x artificial wastewater or 2x artificial wastewater was used. Therefore, wastewater containing high concentrations of TOC, PO_4_^3-^, and NH_4_^+^ could be treated efficiently. However, the composition of actual urban wastewater varies greatly depending on factors, such as region and season. Future studies should examine the combination of *C. reinhardtii* NIES-2238 and *S. cerevisiae* SH-4 established in this study, including the treatment of wastewater with various TOC, PO_4_^3-^, and NH_4_^+^ concentrations and actual wastewater containing various compounds.

Among the genes with increased expression in *C. reinhardtii* NIES-2238, many were involved in transporting molecules and ions, such as transporter activity and inorganic molecule transmembrane transporter activity (Fig. 4a). These genes included those encoding proteins responsible for the uptake of phosphate (CHLRE_02g144750v5) and ammonium (CHLRE_03g159254v5) ions (Supplementary Table 1). Therefore, the combination of *C. reinhardtii* NIES-2238 and *S. cerevisiae* SH-4 may have increased the expression of these genes, contributing to increased phosphorus and nitrogen removal rates.

The GO terms of the downregulated genes in the microalga *C. reinhardtii* NIES-2238 were mainly related to thylakoids, chloroplasts, and photosynthesis (Fig. 4b). Previous studies have reported that when microalgae and heterotrophic microorganisms are co-cultured, light is shielded by heterotrophic microorganisms, and the photosynthetic activity of microalgae is suppressed (Karitani et al., 2024a; Yamada et al., 2023). In the present study, the photosynthetic activity of microalgae could have been suppressed by the co-utilization of microorganisms for wastewater treatment. However, in this study, the combination of *C. reinhardtii* NIES-2238 and *S. cerevisiae* SH-4 did not decrease the removal rate of nutrients but rather increased it. Therefore, it is likely that the light-shielding effect of the co-utilization of these microorganisms did not adversely affect the nutrient removal efficiency. However, the effect of light irradiation conditions on nutrient removal efficiency is unclear, and future optimization of light irradiation conditions for wastewater treatment combining *C. reinhardtii* NIES-2238 and *S. cerevisiae* SH-4 is required.

The GO terms associated with upregulated yeast genes in wastewater treatment in combination with the microalga *C. reinhardtii NIES-2238* were primarily linked to organelles (Fig. 5a). Among the genes in this GO term were those related to the mitochondria, a type of organelle. The mitochondria are responsible for ATP production (Dimmer et al., 2002; Frey et al., 2000). Previous studies have reported that ATP depletion within *S. cerevisiae* causes cell damage (Schimz et al., 1979). Therefore, when the expression of mitochondrial genes is elevated, the mitochondria are more active in ATP production, and the use of this ATP may protect yeast from cell damage. In line with this, *ATP1*, which is involved in ATP synthesis, was among the upregulated genes (Supplementary Table 3). Therefore, the increased expression of these genes may have contributed to protecting yeast from cell damage and improving the efficiency of wastewater treatment. Among the yeast genes with elevated expression (*TRX1* and *GRX3*), those that protect cells from oxidative stress and damage were identified (Supplementary Table 3). In general, toxicity to microorganisms caused by oxidative stress is an issue in wastewater treatment using microorganisms (Osundeko et al., 2013). Therefore, the microorganisms used must be resistant to oxidative stress. In this study, the combination of microalgae and yeast increased the expression of genes that protect cells from various types of damage and stress, which may have contributed to improved efficiency of wastewater treatment.

The GO terms of *S. cerevisiae* SH-4, genes that were downregulated in wastewater treatment in combination with *C. reinhardtii* NIES-2238 included cell process control, binding, and cell development (Fig. 5b). These functions are associated with yeast cell growth. Previous studies have reported that in co-culture, competition for microbial growth occurs and microbial growth is inhibited (Karitani et al., 2024a). Therefore, *S. cerevisiae* SH-4 growth potential may have been reduced in wastewater treatment combined with *C. reinhardtii* NIES-2238 in the present study. However, in this study, the nutrient removal rate was improved by the combined wastewater treatment, indicating that *S. cerevisiae* SH-4 growth potential did not decrease or that the reduction in *S. cerevisiae* SH-4 growth potential did not have a negative effect on nutrient removal.

## 4. Conclusion

Three microalgae and five heterotrophic microorganisms were combined to determine the optimal combination for wastewater treatment. The results showed that the combination of *C. reinhardtii* NIES-2238 and *S. cerevisiae* SH-4 for wastewater treatment improved nitrogen and phosphorus removal efficiency. To the best of our knowledge, this is the first study to show that a combination of green algae and yeast improves the efficiency of wastewater treatment. Transcriptome analysis revealed the reason for the improved wastewater treatment performance of the combination of green algae and yeast. As both the green alga *C. reinhardtii* and the yeast *S. cerevisiae* are highly safe microorganisms, the establishment of their effective combination for wastewater treatment is highly significant. Based on the findings of this study, it is expected that the combination of *C. reinhardtii* and *S. cerevisiae* will further improve the efficiency of wastewater treatment, thereby establishing this technology as a new alternative to the activated sludge method of wastewater treatment.

## 5. Material and methods

### 5.1. Media and artificial wastewater

The BG11Y medium (1 vol% BG11 broth for microbiology [Sigma-Aldrich Japan, Tokyo, Japan], 5 g/L yeast extract [Formedium, Norfolk, UK], 0.1 vol% trace metal mix A5 + Co [Sigma-Aldrich Japan]), SOT medium (16.8 g/L NaHCO_3_ [Nacalai Tesque, Kyoto, Japan], 500 mg/L K_2_HPO_4_ [Nacalai Tesque], 2500 mg/L NaNO_3_ [Nacalai Tesque], 1000 mg/L K_2_SO_4_ [Wako Pure Chemicals], 1000 mg/L NaCl [Nacalai Tesque], 200 mg/L MgSO_4_·7H_2_O [Nacalai Tesque], 40 mg/L CaCl_2_·2H_2_O [Nacalai Tesque], 10 mg/L FeSO_4_·7H_2_O [Wako Pure Chemicals], 80 mg/L Na_2_EDTA·2H_2_O [Nacalai Tesque], 0. 1 vol% trace metal mix A5+Co, pH 9.0), yeast/peptone/glucose (YPD) medium (10 g/L yeast extract, 20 g/L peptone [Formedium], 20 g/L glucose [Nacalai Tesque]), and NBRC 802 medium (10 g/L Casein peptone [Nacalai Tesque], 2 g/L Yeast extract, 1 g/L MgSO_4_·7H_2_O, pH 7.0) were used. For the solid medium, 20 g/L of agar powder [Nacalai Tesque] was added.

The 1x artificial wastewater (0.400 g/L D-(+)-Glucose, 0.200 g/L acetic acid [Nacalai Tesque], 0.078 g/L NH_4_Cl [Nacalai Tesque], 0.018 g/L KH_2_PO_4_ [Nacalai Tesque], 0.013 g/L MgSO_4_·7H_2_O, 0.043 g/L CaCl_2_·2H_2_O, 0.005 g/L FeSO_4_·7H_2_O, 0.200 g/L H_3_BO_3_ [Nacalai Tesque], 1 mL/L Trace metal mix A5+Co [Sigma-Aldrich Japan], pH 7.5), and 2x artificial wastewater (0.800 g/L D-(+)-Glucose, 0.400 g/L Acetic acid, 0.156 g/L NH_4_Cl, 0.036 g/L KH_2_PO_4_, 0 .013 g/L MgSO_4_·7H_2_O, 0.043 g/L CaCl_2_·2H_2_O, 0.005 g/L FeSO_4_·7H_2_O, 0.200 g/L H_3_BO_3_, 1 mL/L Trace metal mix A5+Co, pH 7.5) were prepared as described in a previous study (Feng et al., 2011) with some modifications. Both types of artificial wastewater were autoclaved at 121°C for 20 min before use.

### 5.2. Microorganisms and pre-culture conditions

The green algae *C. reinhardtii* NIES-2238 (Mohd et al., 2024), green algae *C. vulgaris* NIES-1269 (Tam et al., 1990), and the cyanobacterium *A. platensis* NIES-39 (Hena et al., 2018), autotrophic microorganisms used for wastewater treatment in previous studies, were used. Yeasts *S. cerevisiae* YPH499 (Ghasem et al., 2002), *S. cerevisiae* NBRC 1953 (Ghasem et al., 2002), and *S. cerevisiae* SH-4 (Ghasem et al., 2002), and bacteria *Bacillus subtilis* NBRC 13719 (Guo et al., 2021) and *Bacillus amyloliquefaciens* NBRC 14141 (Xie et al., 2013), heterotrophic microorganisms used for wastewater treatment in previous studies, were also used.

BG11Y medium was used for the pre-culture of *C. reinhardtii* NIES-2238 and *C. vulgaris* NIES-1269. The SOT medium was used for the pre-culture of *A. platensis* NIES-39. Microalgae were pre-cultivated at 30°C, 120 rpm, 60 μmol photons/(m^2^/s) for 5 days.

The YPD medium was used for the pre-culture of *S. cerevisiae* and NBRC 802 medium was used for the pre-culture of *B. subtilis* NBRC 13719 and *B. amyloliquefaciens* NBRC 14141. Heterotrophic microorganisms were pre-cultivated at 30°C and 120 rpm for 3 days.

### 5.3. Treatment of wastewater with microalgae and heterotrophic microorganisms alone and in combination

Pre-cultured microorganisms were inoculated into 50 mL flasks containing 15 mL of artificial wastewater after one wash with artificial wastewater to achieve an OD_750_ of 0.1. Wastewater treatment was performed at 30°C, 120 rpm, and 60 μmol photons/(m^2^/s) for 18 h.

### 5.4. Analysis of Wastewater

To measure TOC, wastewater was centrifuged at 10,000 × *g* for 1 min at 20°C. The supernatant was diluted with ultrapure water and measured using a total organic carbon meter (Shimadzu, Kyoto, Japan). C_8_H_5_KO_4_, NaHCO_3_, and Na_2_CO_3_ were used as standards.

For the determination of PO_4_^3-^ concentration, the effluent was centrifuged at 10,000 × *g* for 1 min at 20°C, and the supernatant was diluted appropriately with ultrapure water. Subsequently, 2 mL of the diluted supernatant and 400 μL of potassium peroxodisulfate solution (40 g/L potassium peroxodisulfate) were mixed and heated in a pressure vessel at 126°C for 30 min. After cooling the pressure vessel to room temperature, 900 μL of the reaction solution and 60 μL of molybdenum chromogenic solution (10 g/L hexammonium heptamolybdate tetrahydrate, 400 mg bis[(+)tartrate]diantimony(Ⅲ)acid dipotassium trihydrate, 133 mL/L concentrated sulfuric acid, 12 g/L ascorbic acid])were mixed. After standing at room temperature for 15 min, the PO_4_^3-^concentration was calculated by measuring Abs_880_ using a spectrophotometer (Shimadzu). Potassium hydrogen phosphate was used as a standard.

For the measurement of NH_4_^+^ concentration, the effluent was centrifuged at 10,000 × *g* for 1 min at 20°C, and the supernatant was diluted with ultrapure water. Subsequently, 2 mL of the diluted supernatant and 1000 μL of sodium hydroxide/potassium peroxodisulfate solution (3.0 g/L NaOH, 20 g/L potassium peroxodisulfate) were mixed and heated in a pressure vessel at 126°C for 30 min. After the pressure vessel was allowed to cool to room temperature, 900 μL of the reaction solution and 60 μL of sodium hydroxide-boron solution [61.8 g/L boric acid, 8.0 g/L NaOH] were mixed, and the NH_4_^+^ concentration was calculated by measuring Abs_220_ using a spectrophotometer. Potassium nitrate was used as the standard.

### 5.5. Transcriptome analysis

Total RNA was extracted from *C. reinhardtii* NIES-2238 and *S. cerevisiae* SH-4 cells using NucleoSpin RNA (Takara Bio, Otsu, Japan) and Quick-RNA MiniPrep Plus (Zymo Research The MGIEasy RNA Directional Library Prep Set (MGI Tech, Shenzhen, China) was used to prepare a complementary DNA library for NGS of the extracted RNA. RNA sequencing was performed using DNBSEQ-G400 (MGI Tech).

The genome sequences of C. reinhardtii CC-503 and S. cerevisiae S288c were used as reference sequences for read mapping using Geneious Prime version 2020.0.3 (Tomy Digital Biology, Tokyo, Japan). DEGs were defined as genes satisfying the following conditions: significance probability (p-value) of <0.01 and differential expression log_2_ ratio of <−1.5 or >1.5. RNA sequencing data were deposited in the DNA Data Bank of Japan nucleotide sequence database under the accession number DRR568808–DRR568810.

## Author Contributions

Miiku Takahashi: Investigation, Writing-Original Draft. Yukino Karitani: Investigation. Ryosuke Yamada: Conceptualization, Writing-Original Draft, Writing-Review & Editing, Supervision, Funding acquisition. Takuya Matsumoto: Supervision. Hiroyasu Ogino: Supervision

## Acknowledgements

The microalgae *C. reinhardtii* NIES-2238, *C. vulgaris* NIES-1269, and *A. platensis* NIES-39 were provided by NIES through the NBRP of MEXT (Japan). We thank Editage for the English language editing.

## Data Availability

The data supporting the findings of this study are available from the corresponding author upon reasonable request.

## Supplementary information

**Supplementary Table 1.** Microalgae genes whose expression levels increased due to co-cultivation of microalgae and yeast.

| locus tag<sup>a</sup> | -Log<sub>10</sub> (p-value) | Log<sub>2</sub> (fold change) |
| --- | --- | --- |
| CHLRE_10g433200v5 | 2.14 | 10.72 |
| CHLRE_16g691750v5 | 2.14 | 9.66 |
| CHLRE_03g159254v5 | 10.46 | 9.59 |
| CHLRE_10g433250v5 | 4.13 | 9.38 |
| CHLRE_01g009400v5 | 2.22 | 8.87 |
| CHLRE_17g708025v5 | 8.57 | 8.71 |
| CHLRE_10g433250v5 | 3.61 | 8.61 |
| CHLRE_02g089400v5 | 3.18 | 8.54 |
| CHLRE_07g346950v5 | 4.69 | 8.41 |
| CHLRE_03g206500v5 | 3.19 | 8.38 |
| CHLRE_02g096551v5 | 14.82 | 8.31 |
| CHLRE_10g433250v5 | 2.63 | 8 |
| CHLRE_03g204500v5 | 2.67 | 7.3 |
| CHLRE_05g241634v5 | 9.19 | 7.14 |
| CHLRE_07g325740v5 | 10.99 | 7.1 |
| CHLRE_09g390838v5 | 13.1 | 6.98 |
| CHLRE_12g551350v5 | 7.51 | 6.94 |
| CHLRE_06g263950v5 | 5.76 | 6.85 |
| CHLRE_10g425100v5 | 7.38 | 6.51 |
| CHLRE_10g433200v5 | 2.11 | 6.47 |
| CHLRE_01g011150v5 | 8.54 | 6.4 |
| CHLRE_10g433200v5 | 3.12 | 6.21 |
| CHLRE_16g679150v5 | 20.03 | 6.18 |
| CHLRE_07g346050v5 | 6.03 | 6.03 |
| CHLRE_10g442800v5 | 2.13 | 5.86 |
| CHLRE_01g009350v5 | 12.79 | 5.83 |
| CHLRE_17g702900v5 | 16.99 | 5.53 |
| CHLRE_09g410050v5 | 6.15 | 5.5 |
| CHLRE_10g442800v5 | 2.61 | 5.49 |
| CHLRE_05g234656v5 | 4.7 | 5.48 |
| CHLRE_09g410050v5 | 3.94 | 5.42 |
| CHLRE_06g252750v5 | 18.65 | 5.35 |
| CHLRE_01g008300v5 | 16.35 | 5.32 |
| CHLRE_09g400750v5 | 2.15 | 5.24 |
| CHLRE_07g350100v5 | 5.89 | 5.22 |
| CHLRE_08g362500v5 | 13.95 | 5.15 |
| CHLRE_01g012650v5 | 3.71 | 5.15 |
| CHLRE_16g673250v5 | 4.37 | 5.02 |
| CHLRE_03g200750v5 | 6.2 | 4.81 |
| CHLRE_12g546600v5 | 5.98 | 4.81 |
| CHLRE_11g467709v5 | 8.02 | 4.72 |
| CHLRE_07g315050v5 | 8.8 | 4.67 |
| CHLRE_01g012700v5 | 19 | 4.64 |
| CHLRE_11g467709v5 | 6.16 | 4.53 |
| CHLRE_09g393700v5 | 11.76 | 4.47 |
| CHLRE_03g149350v5 | 3.3 | 4.47 |
| CHLRE_03g199050v5 | 4.03 | 4.38 |
| CHLRE_10g433476v5 | 3.14 | 4.38 |
| CHLRE_17g735375v5 | 15.83 | 4.35 |
| CHLRE_04g216200v5 | 2.15 | 4.35 |
| CHLRE_10g448700v5 | 9.55 | 4.33 |
| CHLRE_16g656750v5 | 2.04 | 4.33 |
| CHLRE_02g093600v5 | 19.96 | 4.31 |
| CHLRE_06g271450v5 | 11.25 | 4.27 |
| CHLRE_07g321750v5 | 4.54 | 4.26 |
| CHLRE_01g007400v5 | 2.03 | 4.24 |
| CHLRE_11g467539v5 | 3.38 | 4.22 |
| CHLRE_03g212977v5 | 3.34 | 4.18 |
| CHLRE_03g165050v5 | 9.8 | 4.17 |
| CHLRE_10g433476v5 | 4.19 | 4.17 |
| CHLRE_10g433476v5 | 4.1 | 4.16 |
| CHLRE_04g214600v5 | 2.05 | 4.15 |
| CHLRE_11g467539v5 | 2.6 | 4.13 |
| CHLRE_01g034850v5 | 16.22 | 4.07 |
| CHLRE_07g347000v5 | 15.71 | 4.06 |
| CHLRE_12g547750v5 | 17.39 | 4.04 |
| CHLRE_02g145628v5 | 4.19 | 3.99 |
| CHLRE_02g087900v5 | 4.04 | 3.96 |
| CHLRE_12g525450v5 | 3.41 | 3.91 |
| CHLRE_09g393700v5 | 7.33 | 3.89 |
| CHLRE_04g222800v5 | 2.22 | 3.89 |
| CHLRE_07g346900v5 | 2.19 | 3.88 |
| CHLRE_09g414416v5 | 2.15 | 3.88 |
| CHLRE_14g619854v5 | 2.13 | 3.87 |
| CHLRE_16g657200v5 | 16.7 | 3.86 |
| CHLRE_10g445000v5 | 3.38 | 3.86 |
| CHLRE_06g257200v5 | 6.23 | 3.83 |
| CHLRE_06g309000v5 | 5.66 | 3.8 |
| CHLRE_03g203500v5 | 2.13 | 3.79 |
| CHLRE_02g144750v5 | 33.32 | 3.75 |
| CHLRE_16g674065v5 | 11.29 | 3.75 |
| CHLRE_01g012850v5 | 7.62 | 3.72 |
| CHLRE_17g742250v5 | 6.55 | 3.72 |
| CHLRE_07g343050v5 | 6.95 | 3.7 |
| CHLRE_10g445050v5 | 8.23 | 3.66 |
| CHLRE_06g300500v5 | 2.88 | 3.64 |
| CHLRE_01g066187v5 | 2.75 | 3.64 |
| CHLRE_02g099950v5 | 23.05 | 3.6 |
| CHLRE_12g552650v5 | 27.16 | 3.58 |
| CHLRE_07g332275v5 | 10.34 | 3.58 |
| CHLRE_03g202100v5 | 25.66 | 3.56 |
| CHLRE_17g700750v5 | 8.21 | 3.56 |
| CHLRE_10g445050v5 | 9.02 | 3.53 |
| CHLRE_16g681750v5 | 29.61 | 3.52 |
| CHLRE_17g708750v5 | 12.59 | 3.48 |
| CHLRE_06g271300v5 | 8.33 | 3.48 |
| CHLRE_17g733100v5 | 8.24 | 3.48 |
| CHLRE_01g008350v5 | 19.66 | 3.47 |
| CHLRE_02g095137v5 | 11.81 | 3.46 |
| CHLRE_13g587550v5 | 10.27 | 3.46 |
| CHLRE_02g108700v5 | 2.08 | 3.45 |
| CHLRE_07g325727v5 | 8.03 | 3.44 |
| CHLRE_06g294200v5 | 2.41 | 3.44 |
| CHLRE_12g524800v5 | 5.51 | 3.42 |
| CHLRE_02g107256v5 | 2.48 | 3.38 |
| CHLRE_17g711750v5 | 2.82 | 3.35 |
| CHLRE_13g572600v5 | 16.07 | 3.32 |
| CHLRE_17g730100v5 | 6.98 | 3.29 |
| CHLRE_10g452950v5 | 11.04 | 3.28 |
| CHLRE_10g445050v5 | 7.52 | 3.27 |
| CHLRE_02g144500v5 | 30.76 | 3.26 |
| CHLRE_07g342700v5 | 3.51 | 3.26 |
| CHLRE_10g451350v5 | 2.47 | 3.23 |
| CHLRE_10g442600v5 | 3.94 | 3.22 |
| CHLRE_11g467644v5 | 7.65 | 3.19 |
| CHLRE_06g307500v5 | 20.06 | 3.16 |
| CHLRE_01g049950v5 | 14.29 | 3.16 |
| CHLRE_02g112550v5 | 3.45 | 3.16 |
| CHLRE_10g458850v5 | 2.12 | 3.15 |
| CHLRE_02g112500v5 | 9.73 | 3.14 |
| CHLRE_17g713450v5 | 6.96 | 3.14 |
| CHLRE_14g616700v5 | 5.78 | 3.13 |
| CHLRE_10g458850v5 | 2.33 | 3.12 |
| CHLRE_16g663000v5 | 5.35 | 3.11 |
| CHLRE_09g397660v5 | 2.75 | 3.11 |
| CHLRE_16g675100v5 | 2.91 | 3.1 |
| CHLRE_08g359450v5 | 9.36 | 3.08 |
| CHLRE_07g339200v5 | 8.44 | 3.08 |
| CHLRE_06g264050v5 | 3.77 | 3.08 |
| CHLRE_03g144787v5 | 4.99 | 3.06 |
| CHLRE_10g458850v5 | 4.19 | 3.06 |
| CHLRE_12g549427v5 | 3.55 | 3.05 |
| CHLRE_04g220850v5 | 5.08 | 3.04 |
| CHLRE_17g714650v5 | 5.17 | 3.03 |
| CHLRE_03g168200v5 | 34.35 | 2.99 |
| CHLRE_11g468450v5 | 16.47 | 2.99 |
| CHLRE_11g479600v5 | 3.58 | 2.99 |
| CHLRE_10g442600v5 | 2.85 | 2.99 |
| CHLRE_02g145950v5 | 2.27 | 2.99 |
| CHLRE_02g145100v5 | 18.11 | 2.98 |
| CHLRE_04g229550v5 | 16.96 | 2.98 |
| CHLRE_17g710300v5 | 7.61 | 2.98 |
| CHLRE_06g266800v5 | 26.56 | 2.97 |
| CHLRE_16g690319v5 | 5.87 | 2.97 |
| CHLRE_12g550702v5 | 27.18 | 2.96 |
| CHLRE_06g264000v5 | 19.73 | 2.95 |
| CHLRE_02g095076v5 | 18.25 | 2.95 |
| CHLRE_09g398900v5 | 4.72 | 2.94 |
| CHLRE_01g000150v5 | 2.21 | 2.93 |
| CHLRE_03g168100v5 | 9.3 | 2.91 |
| CHLRE_15g635998v5 | 2.29 | 2.9 |
| CHLRE_17g702950v5 | 14.66 | 2.88 |
| CHLRE_06g276950v5 | 8.21 | 2.88 |
| CHLRE_17g735450v5 | 4.06 | 2.88 |
| CHLRE_11g467710v5 | 12.83 | 2.87 |
| CHLRE_07g332250v5 | 4.65 | 2.84 |
| CHLRE_17g710450v5 | 4.3 | 2.83 |
| CHLRE_12g531000v5 | 10.85 | 2.82 |
| CHLRE_12g507800v5 | 4.37 | 2.82 |
| CHLRE_11g478800v5 | 12.01 | 2.81 |
| CHLRE_12g541250v5 | 2.3 | 2.81 |
| CHLRE_09g389504v5 | 5.4 | 2.8 |
| CHLRE_06g274300v5 | 2.83 | 2.79 |
| CHLRE_13g563100v5 | 10.65 | 2.78 |
| CHLRE_10g417650v5 | 7.84 | 2.78 |
| CHLRE_17g713500v5 | 10.07 | 2.76 |
| CHLRE_17g696500v5 | 3.78 | 2.76 |
| CHLRE_06g269850v5 | 22.09 | 2.75 |
| CHLRE_17g714100v5 | 9.25 | 2.74 |
| CHLRE_09g398900v5 | 3.36 | 2.73 |
| CHLRE_11g467744v5 | 2.28 | 2.71 |
| CHLRE_09g409650v5 | 20.06 | 2.7 |
| CHLRE_17g739000v5 | 9.09 | 2.69 |
| CHLRE_12g504600v5 | 7.81 | 2.67 |
| CHLRE_11g467599v5 | 7.55 | 2.67 |
| CHLRE_09g409650v5 | 20.31 | 2.65 |
| CHLRE_09g389504v5 | 5.79 | 2.65 |
| CHLRE_10g454150v5 | 5.44 | 2.65 |
| CHLRE_09g397660v5 | 3.35 | 2.65 |
| CHLRE_09g389430v5 | 9.73 | 2.64 |
| CHLRE_12g551100v5 | 2.71 | 2.64 |
| CHLRE_01g035700v5 | 2.43 | 2.63 |
| CHLRE_02g095087v5 | 9.89 | 2.62 |
| CHLRE_17g709100v5 | 8.67 | 2.62 |
| CHLRE_05g238687v5 | 3.83 | 2.62 |
| CHLRE_13g586700v5 | 9.51 | 2.61 |
| CHLRE_11g467710v5 | 9.52 | 2.6 |
| CHLRE_13g586450v5 | 5.87 | 2.59 |
| CHLRE_03g174400v5 | 5.05 | 2.59 |
| CHLRE_16g658800v5 | 9.2 | 2.58 |
| CHLRE_01g053000v5 | 11.62 | 2.57 |
| CHLRE_04g222750v5 | 6.3 | 2.55 |
| CHLRE_09g396650v5 | 4.44 | 2.55 |
| CHLRE_06g265400v5 | 4.87 | 2.54 |
| CHLRE_02g097050v5 | 14.25 | 2.53 |
| CHLRE_07g331450v5 | 3.59 | 2.53 |
| CHLRE_12g506450v5 | 6.39 | 2.52 |
| CHLRE_06g274150v5 | 3.99 | 2.52 |
| CHLRE_01g012950v5 | 3.2 | 2.52 |
| CHLRE_11g479600v5 | 3.59 | 2.51 |
| CHLRE_08g374450v5 | 4.26 | 2.5 |
| CHLRE_16g677100v5 | 2.7 | 2.5 |
| CHLRE_14g619050v5 | 8.06 | 2.49 |
| CHLRE_06g292900v5 | 3.08 | 2.49 |
| CHLRE_06g283200v5 | 2.54 | 2.49 |
| CHLRE_17g707137v5 | 7.46 | 2.48 |
| CHLRE_17g714800v5 | 2.2 | 2.48 |
| CHLRE_17g700650v5 | 12.7 | 2.47 |
| CHLRE_06g278252v5 | 3.79 | 2.47 |
| CHLRE_08g374436v5 | 9.58 | 2.46 |
| CHLRE_10g442600v5 | 2.45 | 2.46 |
| CHLRE_03g177350v5 | 13.38 | 2.45 |
| CHLRE_12g554350v5 | 8.05 | 2.45 |
| CHLRE_11g467594v5 | 15.08 | 2.44 |
| CHLRE_11g467601v5 | 7.46 | 2.44 |
| CHLRE_12g550450v5 | 7.08 | 2.44 |
| CHLRE_12g489700v5 | 6.7 | 2.44 |
| CHLRE_10g440150v5 | 3.34 | 2.44 |
| CHLRE_08g378150v5 | 2.26 | 2.44 |
| CHLRE_12g488000v5 | 11.12 | 2.43 |
| CHLRE_07g349119v5 | 7.84 | 2.43 |
| CHLRE_02g097800v5 | 5.15 | 2.43 |
| CHLRE_07g332950v5 | 5.15 | 2.43 |
| CHLRE_04g216550v5 | 4.29 | 2.43 |
| CHLRE_17g714600v5 | 9.1 | 2.42 |
| CHLRE_17g714050v5 | 3.85 | 2.42 |
| CHLRE_02g142650v5 | 3.6 | 2.4 |
| CHLRE_13g567700v5 | 23.89 | 2.39 |
| CHLRE_02g089900v5 | 6.19 | 2.39 |
| CHLRE_12g523026v5 | 6.68 | 2.38 |
| CHLRE_12g523000v5 | 6.77 | 2.37 |
| CHLRE_06g267950v5 | 4.6 | 2.37 |
| CHLRE_05g243450v5 | 2.05 | 2.37 |
| CHLRE_09g389850v5 | 5.66 | 2.36 |
| CHLRE_09g389850v5 | 5.69 | 2.35 |
| CHLRE_06g275800v5 | 5.37 | 2.35 |
| CHLRE_03g155300v5 | 8.75 | 2.34 |
| CHLRE_09g396600v5 | 9.23 | 2.33 |
| CHLRE_17g714550v5 | 6.22 | 2.33 |
| CHLRE_12g549000v5 | 4.15 | 2.33 |
| CHLRE_14g629840v5 | 3.06 | 2.33 |
| CHLRE_10g454450v5 | 2.34 | 2.33 |
| CHLRE_12g519750v5 | 44.72 | 2.32 |
| CHLRE_03g156550v5 | 12.73 | 2.32 |
| CHLRE_06g296750v5 | 11.84 | 2.32 |
| CHLRE_03g200850v5 | 33.03 | 2.31 |
| CHLRE_17g735876v5 | 2.55 | 2.31 |
| CHLRE_17g723750v5 | 25.8 | 2.3 |
| CHLRE_03g182700v5 | 12.69 | 2.3 |
| CHLRE_09g396600v5 | 8.98 | 2.3 |
| CHLRE_06g271376v5 | 8.2 | 2.3 |
| CHLRE_09g402500v5 | 4.06 | 2.3 |
| CHLRE_12g538000v5 | 4.03 | 2.3 |
| CHLRE_09g406600v5 | 3.29 | 2.3 |
| CHLRE_10g454150v5 | 2.53 | 2.3 |
| CHLRE_09g405500v5 | 2.35 | 2.3 |
| CHLRE_02g109450v5 | 4.6 | 2.27 |
| CHLRE_16g693601v5 | 4.6 | 2.27 |
| CHLRE_03g181450v5 | 2.59 | 2.26 |
| CHLRE_04g217915v5 | 19.13 | 2.25 |
| CHLRE_12g498650v5 | 9.92 | 2.25 |
| CHLRE_12g517850v5 | 3.39 | 2.25 |
| CHLRE_17g705500v5 | 2.12 | 2.25 |
| CHLRE_16g663950v5 | 11.83 | 2.24 |
| CHLRE_12g492950v5 | 11.72 | 2.24 |
| CHLRE_17g713550v5 | 6.42 | 2.24 |
| CHLRE_17g708450v5 | 2.32 | 2.24 |
| CHLRE_01g008550v5 | 2.24 | 2.24 |
| CHLRE_14g620600v5 | 15.58 | 2.23 |
| CHLRE_02g115950v5 | 7.31 | 2.23 |
| CHLRE_04g225200v5 | 3.17 | 2.23 |
| CHLRE_17g720350v5 | 7.5 | 2.22 |
| CHLRE_09g394200v5 | 5.69 | 2.22 |
| CHLRE_08g374300v5 | 3.43 | 2.22 |
| CHLRE_08g370401v5 | 3.33 | 2.22 |
| CHLRE_08g374400v5 | 3.47 | 2.21 |
| CHLRE_17g714200v5 | 43.5 | 2.2 |
| CHLRE_06g306000v5 | 2.26 | 2.2 |
| CHLRE_13g605200v5 | 19.34 | 2.19 |
| CHLRE_02g101700v5 | 3.95 | 2.18 |
| CHLRE_09g413150v5 | 2.9 | 2.18 |
| CHLRE_11g467594v5 | 19.48 | 2.17 |
| CHLRE_12g506200v5 | 15.32 | 2.17 |
| CHLRE_03g149400v5 | 6.12 | 2.17 |
| CHLRE_16g681800v5 | 2.38 | 2.17 |
| CHLRE_12g548500v5 | 2.27 | 2.17 |
| CHLRE_14g625600v5 | 2.83 | 2.16 |
| CHLRE_13g589800v5 | 2.38 | 2.16 |
| CHLRE_04g224800v5 | 14.98 | 2.15 |
| CHLRE_12g504800v5 | 5.89 | 2.15 |
| CHLRE_13g584775v5 | 4.73 | 2.15 |
| CHLRE_12g505500v5 | 3.91 | 2.15 |
| CHLRE_06g265050v5 | 10.37 | 2.14 |
| CHLRE_11g468600v5 | 9.37 | 2.14 |
| CHLRE_09g416050v5 | 7.68 | 2.14 |
| CHLRE_17g709050v5 | 3.71 | 2.14 |
| CHLRE_17g732350v5 | 3.04 | 2.14 |
| CHLRE_11g481600v5 | 2.43 | 2.14 |
| CHLRE_16g650600v5 | 22.39 | 2.13 |
| CHLRE_13g582800v5 | 2.41 | 2.13 |
| CHLRE_07g355250v5 | 2.24 | 2.13 |
| CHLRE_17g746997v5 | 36.33 | 2.12 |
| CHLRE_07g348350v5 | 5.09 | 2.12 |
| CHLRE_11g481600v5 | 2.25 | 2.12 |
| CHLRE_10g452800v5 | 6.05 | 2.11 |
| CHLRE_16g690506v5 | 3.63 | 2.11 |
| CHLRE_06g260450v5 | 3.15 | 2.11 |
| CHLRE_09g413150v5 | 2.72 | 2.11 |
| CHLRE_12g505100v5 | 2.02 | 2.11 |
| CHLRE_17g717900v5 | 6.16 | 2.1 |
| CHLRE_09g387208v5 | 4.73 | 2.1 |
| CHLRE_01g034250v5 | 3.22 | 2.1 |
| CHLRE_17g711800v5 | 17.25 | 2.09 |
| CHLRE_06g271800v5 | 8.92 | 2.09 |
| CHLRE_06g290000v5 | 6.09 | 2.09 |
| CHLRE_09g387600v5 | 2.31 | 2.09 |
| CHLRE_07g336950v5 | 23.8 | 2.08 |
| CHLRE_03g207900v5 | 6.61 | 2.08 |
| CHLRE_06g265450v5 | 3.57 | 2.08 |
| CHLRE_04g219600v5 | 3.36 | 2.08 |
| CHLRE_09g403182v5 | 4.96 | 2.07 |
| CHLRE_09g387949v5 | 6.72 | 2.05 |
| CHLRE_02g076466v5 | 6.18 | 2.05 |
| CHLRE_13g570600v5 | 5.3 | 2.05 |
| CHLRE_09g394200v5 | 4.76 | 2.05 |
| CHLRE_14g631000v5 | 4.5 | 2.05 |
| CHLRE_02g113050v5 | 2.57 | 2.05 |
| CHLRE_03g172300v5 | 19.69 | 2.04 |
| CHLRE_10g439850v5 | 8.56 | 2.04 |
| CHLRE_16g672602v5 | 8.06 | 2.03 |
| CHLRE_16g667850v5 | 5.48 | 2.03 |
| CHLRE_17g714000v5 | 4.96 | 2.03 |
| CHLRE_06g275700v5 | 4.5 | 2.03 |
| CHLRE_10g432250v5 | 2.78 | 2.03 |
| CHLRE_12g537371v5 | 6.27 | 2.02 |
| CHLRE_14g625300v5 | 2.88 | 2.02 |
| CHLRE_10g435200v5 | 2.07 | 2.02 |
| CHLRE_16g669150v5 | 9.09 | 2.01 |
| CHLRE_12g506350v5 | 8.86 | 2 |
| CHLRE_09g392951v5 | 3.48 | 2 |
| CHLRE_12g525700v5 | 2.92 | 2 |
| CHLRE_06g265350v5 | 5.12 | 1.99 |
| CHLRE_15g637900v5 | 2.16 | 1.99 |
| CHLRE_07g344550v5 | 7.5 | 1.98 |
| CHLRE_14g613950v5 | 7.23 | 1.98 |
| CHLRE_06g295700v5 | 4.12 | 1.98 |
| CHLRE_09g387467v5 | 3.01 | 1.98 |
| CHLRE_06g282850v5 | 2.26 | 1.98 |
| CHLRE_06g288600v5 | 3.81 | 1.97 |
| CHLRE_13g584800v5 | 3.15 | 1.96 |
| CHLRE_06g293950v5 | 12.76 | 1.95 |
| CHLRE_03g159300v5 | 7.82 | 1.95 |
| CHLRE_02g141926v5 | 7.44 | 1.95 |
| CHLRE_06g276800v5 | 4.36 | 1.95 |
| CHLRE_16g695850v5 | 4 | 1.95 |
| CHLRE_06g264800v5 | 2.61 | 1.95 |
| CHLRE_06g283400v5 | 5.58 | 1.94 |
| CHLRE_08g370450v5 | 4.37 | 1.94 |
| CHLRE_11g467725v5 | 2.68 | 1.94 |
| CHLRE_07g346350v5 | 2.58 | 1.94 |
| CHLRE_03g200000v5 | 4.46 | 1.93 |
| CHLRE_06g274900v5 | 3.11 | 1.93 |
| CHLRE_02g074800v5 | 2.64 | 1.93 |
| CHLRE_06g274200v5 | 2.45 | 1.93 |
| CHLRE_09g401960v5 | 2.42 | 1.93 |
| CHLRE_11g468600v5 | 7.6 | 1.92 |
| CHLRE_03g178650v5 | 4.11 | 1.92 |
| CHLRE_09g396650v5 | 3.44 | 1.92 |
| CHLRE_04g217944v5 | 2.59 | 1.92 |
| CHLRE_15g639350v5 | 2.19 | 1.92 |
| CHLRE_03g199800v5 | 20.36 | 1.91 |
| CHLRE_09g410700v5 | 10.24 | 1.91 |
| CHLRE_02g095073v5 | 8.81 | 1.91 |
| CHLRE_06g264600v5 | 7.2 | 1.91 |
| CHLRE_08g372550v5 | 5.99 | 1.91 |
| CHLRE_06g267650v5 | 5.26 | 1.91 |
| CHLRE_06g276700v5 | 2.9 | 1.91 |
| CHLRE_06g266600v5 | 5.88 | 1.9 |
| CHLRE_17g709150v5 | 5.01 | 1.9 |
| CHLRE_12g504650v5 | 3.16 | 1.9 |
| CHLRE_12g552200v5 | 19.36 | 1.89 |
| CHLRE_17g715300v5 | 13.09 | 1.89 |
| CHLRE_01g009950v5 | 2.9 | 1.89 |
| CHLRE_09g409150v5 | 8.28 | 1.88 |
| CHLRE_02g094450v5 | 5.28 | 1.88 |
| CHLRE_17g711850v5 | 3.62 | 1.88 |
| CHLRE_17g728750v5 | 3.08 | 1.88 |
| CHLRE_12g503450v5 | 2.97 | 1.88 |
| CHLRE_15g636550v5 | 2.72 | 1.88 |
| CHLRE_07g347750v5 | 9.69 | 1.87 |
| CHLRE_08g368850v5 | 8.81 | 1.87 |
| CHLRE_10g458450v5 | 7.09 | 1.87 |
| CHLRE_09g401293v5 | 7.53 | 1.86 |
| CHLRE_17g699750v5 | 5.73 | 1.86 |
| CHLRE_17g710500v5 | 4.29 | 1.86 |
| CHLRE_09g396065v5 | 3.21 | 1.86 |
| CHLRE_16g659667v5 | 2.67 | 1.86 |
| CHLRE_06g273100v5 | 10.19 | 1.85 |
| CHLRE_12g522950v5 | 6.81 | 1.85 |
| CHLRE_13g588150v5 | 4.84 | 1.85 |
| CHLRE_02g098850v5 | 4.77 | 1.85 |
| CHLRE_06g301000v5 | 3.03 | 1.85 |
| CHLRE_02g113200v5 | 20.48 | 1.84 |
| CHLRE_17g714450v5 | 16.22 | 1.84 |
| CHLRE_14g618400v5 | 5.71 | 1.84 |
| CHLRE_10g452800v5 | 4.94 | 1.84 |
| CHLRE_14g620702v5 | 2.55 | 1.84 |
| CHLRE_01g015500v5 | 38.17 | 1.83 |
| CHLRE_06g260200v5 | 6.74 | 1.83 |
| CHLRE_09g399550v5 | 5.38 | 1.83 |
| CHLRE_10g458550v5 | 3.31 | 1.83 |
| CHLRE_07g327900v5 | 3.11 | 1.83 |
| CHLRE_03g164050v5 | 3.06 | 1.83 |
| CHLRE_06g311700v5 | 5.49 | 1.82 |
| CHLRE_12g504950v5 | 5.13 | 1.82 |
| CHLRE_09g387208v5 | 4.21 | 1.82 |
| CHLRE_07g353900v5 | 2.75 | 1.82 |
| CHLRE_02g076650v5 | 2.36 | 1.82 |
| CHLRE_06g276650v5 | 8.38 | 1.8 |
| CHLRE_16g673057v5 | 5.51 | 1.8 |
| CHLRE_10g458500v5 | 3.14 | 1.8 |
| CHLRE_16g653450v5 | 2.92 | 1.8 |
| CHLRE_13g592200v5 | 32.82 | 1.79 |
| CHLRE_14g620850v5 | 20.62 | 1.79 |
| CHLRE_13g588501v5 | 12.3 | 1.79 |
| CHLRE_02g083800v5 | 6.02 | 1.79 |
| CHLRE_05g232600v5 | 5.35 | 1.79 |
| CHLRE_10g431350v5 | 3.78 | 1.79 |
| CHLRE_09g394436v5 | 18.55 | 1.78 |
| CHLRE_06g258800v5 | 6.71 | 1.78 |
| CHLRE_11g467599v5 | 3.41 | 1.78 |
| CHLRE_12g526800v5 | 19.21 | 1.77 |
| CHLRE_14g611900v5 | 18.89 | 1.77 |
| CHLRE_09g410700v5 | 7.88 | 1.77 |
| CHLRE_02g095150v5 | 3.91 | 1.77 |
| CHLRE_03g175600v5 | 3.15 | 1.77 |
| CHLRE_01g050400v5 | 2.28 | 1.77 |
| CHLRE_17g725150v5 | 7.72 | 1.76 |
| CHLRE_09g401293v5 | 6.74 | 1.76 |
| CHLRE_02g110650v5 | 6.3 | 1.76 |
| CHLRE_17g712900v5 | 5.16 | 1.76 |
| CHLRE_01g008650v5 | 4.88 | 1.76 |
| CHLRE_14g630883v5 | 2.71 | 1.76 |
| CHLRE_11g467593v5 | 11.38 | 1.75 |
| CHLRE_10g432250v5 | 2.23 | 1.75 |
| CHLRE_08g360450v5 | 2.08 | 1.75 |
| CHLRE_09g391801v5 | 7.96 | 1.74 |
| CHLRE_06g290100v5 | 2.12 | 1.74 |
| CHLRE_02g115450v5 | 2.95 | 1.73 |
| CHLRE_10g432250v5 | 2.4 | 1.73 |
| CHLRE_02g111300v5 | 7.39 | 1.72 |
| CHLRE_06g265500v5 | 4.01 | 1.72 |
| CHLRE_07g336900v5 | 7.39 | 1.71 |
| CHLRE_09g396065v5 | 2.74 | 1.71 |
| CHLRE_12g508900v5 | 3.44 | 1.7 |
| CHLRE_06g285250v5 | 21.71 | 1.69 |
| CHLRE_01g034325v5 | 3.37 | 1.69 |
| CHLRE_12g545550v5 | 2.07 | 1.69 |
| CHLRE_14g622200v5 | 4.07 | 1.68 |
| CHLRE_17g718468v5 | 4 | 1.67 |
| CHLRE_01g008950v5 | 3.97 | 1.67 |
| CHLRE_06g297250v5 | 3.6 | 1.67 |
| CHLRE_09g386756v5 | 2.56 | 1.67 |
| CHLRE_06g273900v5 | 2.19 | 1.67 |
| CHLRE_13g570000v5 | 2.14 | 1.67 |
| CHLRE_02g141166v5 | 4.6 | 1.66 |
| CHLRE_17g735950v5 | 3.59 | 1.66 |
| CHLRE_06g265200v5 | 3.55 | 1.66 |
| CHLRE_17g713400v5 | 3.44 | 1.66 |
| CHLRE_09g395950v5 | 3.32 | 1.66 |
| CHLRE_01g022200v5 | 2.31 | 1.66 |
| CHLRE_09g392503v5 | 2.08 | 1.66 |
| CHLRE_09g388850v5 | 12.58 | 1.65 |
| CHLRE_11g467668v5 | 3.24 | 1.65 |
| CHLRE_10g431350v5 | 3.02 | 1.65 |
| CHLRE_17g734596v5 | 2.09 | 1.65 |
| CHLRE_16g653150v5 | 2.02 | 1.65 |
| CHLRE_03g148700v5 | 5.06 | 1.64 |
| CHLRE_07g350750v5 | 4.28 | 1.64 |
| CHLRE_01g005050v5 | 11.84 | 1.63 |
| CHLRE_08g362450v5 | 10.92 | 1.63 |
| CHLRE_06g268400v5 | 8.28 | 1.63 |
| CHLRE_17g735400v5 | 3.91 | 1.63 |
| CHLRE_03g145267v5 | 3.08 | 1.63 |
| CHLRE_17g722300v5 | 17.65 | 1.62 |
| CHLRE_09g416150v5 | 6.08 | 1.62 |
| CHLRE_01g012750v5 | 3.08 | 1.62 |
| CHLRE_03g189250v5 | 2.89 | 1.62 |
| CHLRE_09g388850v5 | 12.51 | 1.61 |
| CHLRE_11g467649v5 | 6.92 | 1.61 |
| CHLRE_12g485100v5 | 2.91 | 1.61 |
| CHLRE_17g735500v5 | 2.65 | 1.61 |
| CHLRE_04g213450v5 | 2.53 | 1.61 |
| CHLRE_12g554400v5 | 19.05 | 1.6 |
| CHLRE_03g194850v5 | 10.79 | 1.6 |
| CHLRE_17g725200v5 | 10.32 | 1.6 |
| CHLRE_07g340450v5 | 8.59 | 1.6 |
| CHLRE_13g582600v5 | 5.54 | 1.6 |
| CHLRE_10g433950v5 | 3.94 | 1.6 |
| CHLRE_01g013200v5 | 3.25 | 1.6 |
| CHLRE_03g195950v5 | 2.33 | 1.6 |
| CHLRE_06g268100v5 | 4.39 | 1.59 |
| CHLRE_13g563950v5 | 3.02 | 1.59 |
| CHLRE_16g685000v5 | 7.53 | 1.58 |
| CHLRE_03g150300v5 | 4.37 | 1.58 |
| CHLRE_09g399550v5 | 3.39 | 1.58 |
| CHLRE_03g146507v5 | 3.72 | 1.57 |
| CHLRE_12g491050v5 | 6.81 | 1.56 |
| CHLRE_01g004124v5 | 6.45 | 1.56 |
| CHLRE_03g180250v5 | 38.47 | 1.55 |
| CHLRE_16g685700v5 | 10.06 | 1.55 |
| CHLRE_09g416050v5 | 6.42 | 1.55 |
| CHLRE_10g444000v5 | 2.72 | 1.55 |
| CHLRE_06g264750v5 | 2.15 | 1.55 |
| CHLRE_13g574350v5 | 2.72 | 1.54 |
| CHLRE_12g486950v5 | 2.6 | 1.54 |
| CHLRE_03g207250v5 | 23.21 | 1.53 |
| CHLRE_07g347400v5 | 7.15 | 1.53 |
| CHLRE_11g467649v5 | 6.8 | 1.53 |
| CHLRE_12g505600v5 | 2.12 | 1.53 |
| CHLRE_02g077800v5 | 13.64 | 1.51 |
| CHLRE_06g282200v5 | 7.58 | 1.51 |
| CHLRE_17g713950v5 | 3.78 | 1.51 |
| CHLRE_07g326650v5 | 3.2 | 1.51 |
| CHLRE_03g177600v5 | 2.47 | 1.51 |
| CHLRE_10g434950v5 | 2.27 | 1.51 |
| CHLRE_12g512000v5 | 5.55 | 1.5 |
| CHLRE_09g391801v5 | 5.37 | 1.5 |
| CHLRE_12g504550v5 | 4.93 | 1.5 |
| CHLRE_17g709200v5 | 3 | 1.5 |
| CHLRE_02g079450v5 | 2.08 | 1.49 |
| CHLRE_06g268000v5 | 4.5 | 1.48 |
| CHLRE_01g011600v5 | 3.42 | 1.48 |
| CHLRE_09g392750v5 | 2.3 | 1.48 |
| CHLRE_06g311350v5 | 7.27 | 1.47 |
| CHLRE_11g467527v5 | 2.47 | 1.47 |
| CHLRE_01g023400v5 | 4.73 | 1.46 |
| CHLRE_12g515850v5 | 4.21 | 1.46 |
| CHLRE_13g579326v5 | 2.42 | 1.46 |
| CHLRE_06g287450v5 | 2.91 | 1.45 |
| CHLRE_02g105100v5 | 27.6 | 1.43 |
| CHLRE_12g505050v5 | 3.2 | 1.43 |
| CHLRE_07g347300v5 | 4.35 | 1.42 |
| CHLRE_14g627050v5 | 3.87 | 1.42 |
| CHLRE_02g102650v5 | 2.77 | 1.42 |
| CHLRE_01g001400v5 | 2.71 | 1.42 |
| CHLRE_09g395950v5 | 2.5 | 1.42 |
| CHLRE_09g387500v5 | 2 | 1.42 |
| CHLRE_03g145507v5 | 2.35 | 1.41 |
| CHLRE_06g308500v5 | 11.26 | 1.4 |
| CHLRE_08g358580v5 | 5.83 | 1.4 |
| CHLRE_10g461050v5 | 2.65 | 1.4 |
| CHLRE_16g680566v5 | 2.43 | 1.4 |
| CHLRE_02g097600v5 | 7.52 | 1.39 |
| CHLRE_09g386650v5 | 38.84 | 1.38 |
| CHLRE_07g355850v5 | 30.85 | 1.38 |
| CHLRE_03g200050v5 | 7.79 | 1.38 |
| CHLRE_16g660225v5 | 4.96 | 1.38 |
| CHLRE_02g142600v5 | 3.82 | 1.37 |
| CHLRE_08g367550v5 | 3.52 | 1.37 |
| CHLRE_10g459750v5 | 3.9 | 1.36 |
| CHLRE_02g095077v5 | 3.72 | 1.35 |
| CHLRE_06g268976v5 | 2.64 | 1.35 |
| CHLRE_03g176250v5 | 2.45 | 1.35 |
| CHLRE_01g004800v5 | 3.13 | 1.34 |
| CHLRE_11g467781v5 | 2.64 | 1.34 |
| CHLRE_12g504850v5 | 2.42 | 1.34 |
| CHLRE_09g411900v5 | 2.31 | 1.34 |
| CHLRE_03g176150v5 | 4.77 | 1.33 |
| CHLRE_10g419250v5 | 4.61 | 1.32 |
| CHLRE_11g468550v5 | 2.02 | 1.32 |
| CHLRE_11g467567v5 | 6.72 | 1.31 |
| CHLRE_06g278140v5 | 11.91 | 1.3 |
| CHLRE_06g257950v5 | 2.61 | 1.3 |
| CHLRE_02g079300v5 | 2.23 | 1.3 |
| CHLRE_16g675958v5 | 2.12 | 1.3 |
| CHLRE_03g177711v5 | 2.08 | 1.29 |
| CHLRE_06g266250v5 | 4.83 | 1.27 |
| CHLRE_13g567600v5 | 4.71 | 1.27 |
| CHLRE_02g077550v5 | 3.61 | 1.27 |
| CHLRE_02g077850v5 | 3.81 | 1.26 |
| CHLRE_06g278157v5 | 2.85 | 1.26 |
| CHLRE_03g190950v5 | 22.32 | 1.25 |
| CHLRE_03g180750v5 | 2.38 | 1.25 |
| CHLRE_08g358250v5 | 2.19 | 1.25 |
| CHLRE_10g465550v5 | 2.11 | 1.25 |
| CHLRE_01g007850v5 | 2.67 | 1.24 |
| CHLRE_17g741000v5 | 2.67 | 1.24 |
| CHLRE_07g319310v5 | 2.44 | 1.23 |
| CHLRE_10g420700v5 | 2.34 | 1.23 |
| CHLRE_16g683250v5 | 2.1 | 1.23 |
| CHLRE_06g262900v5 | 2.05 | 1.23 |
| CHLRE_09g396100v5 | 6.11 | 1.21 |
| CHLRE_16g685050v5 | 2.24 | 1.2 |
| CHLRE_16g650250v5 | 2.2 | 1.2 |
| CHLRE_10g458450v5 | 2.03 | 1.18 |
| CHLRE_08g379400v5 | 2.19 | 1.17 |
| CHLRE_10g424250v5 | 2.09 | 1.17 |
| CHLRE_08g372900v5 | 6.7 | 1.16 |
| CHLRE_05g238311v5 | 4.95 | 1.16 |
| CHLRE_12g499800v5 | 2.78 | 1.16 |
| CHLRE_03g154350v5 | 2.5 | 1.16 |
| CHLRE_02g074511v5 | 3.31 | 1.15 |
| CHLRE_02g141346v5 | 4.89 | 1.13 |
| CHLRE_03g198236v5 | 2.03 | 1.13 |
| CHLRE_16g667450v5 | 5.31 | 1.11 |
| CHLRE_02g081050v5 | 2.28 | 1.11 |
| CHLRE_07g330300v5 | 3.41 | 1.09 |
| CHLRE_14g620500v5 | 2.54 | 1.09 |
| CHLRE_02g086300v5 | 2.35 | 1.09 |
| CHLRE_01g038250v5 | 5.25 | 1.08 |
| CHLRE_02g114600v5 | 2.82 | 1.08 |
| CHLRE_06g278162v5 | 2.53 | 1.07 |
| CHLRE_02g142687v5 | 4.57 | 1.06 |
| CHLRE_01g027000v5 | 8.14 | 1.04 |
| CHLRE_12g521300v5 | 6.91 | 1.04 |
| CHLRE_07g321400v5 | 3.68 | 1.04 |
| CHLRE_01g010400v5 | 2.46 | 1.04 |
| CHLRE_01g045550v5 | 2.44 | 1.04 |
| CHLRE_17g718500v5 | 2.22 | 1.02 |
| CHLRE_04g216850v5 | 5.03 | 1.01 |
| CHLRE_07g331550v5 | 4.01 | 1.01 |
| CHLRE_06g251000v5 | 2.45 | 1 |
| CHLRE_01g042750v5 | 2.44 | 1 |
| CHLRE_06g257450v5 | 2.35 | 0.98 |
| CHLRE_14g619250v5 | 5.19 | 0.97 |
| CHLRE_02g095140v5 | 4.62 | 0.97 |
| CHLRE_03g159800v5 | 2.75 | 0.97 |
| CHLRE_05g235355v5 | 2.42 | 0.93 |
| CHLRE_02g116750v5 | 3.11 | 0.77 |
| CHLRE_12g528450v5 | 2.34 | 0.71 |
| CHLRE_09g397697v5 | 3.46 | 0.7 |
| CHLRE_12g549550v5 | 5.26 | 0.69 |
| CHLRE_12g516200v5 | 5.67 | 0.65 |
| CHLRE_17g698000v5 | 2.06 | 0.63 |
| CHLRE_12g551501v5 | 2.89 | 0.59 |

<sup>a</sup>Gramene (https://www.gramene.org/)

**Supplementary Table 2.**
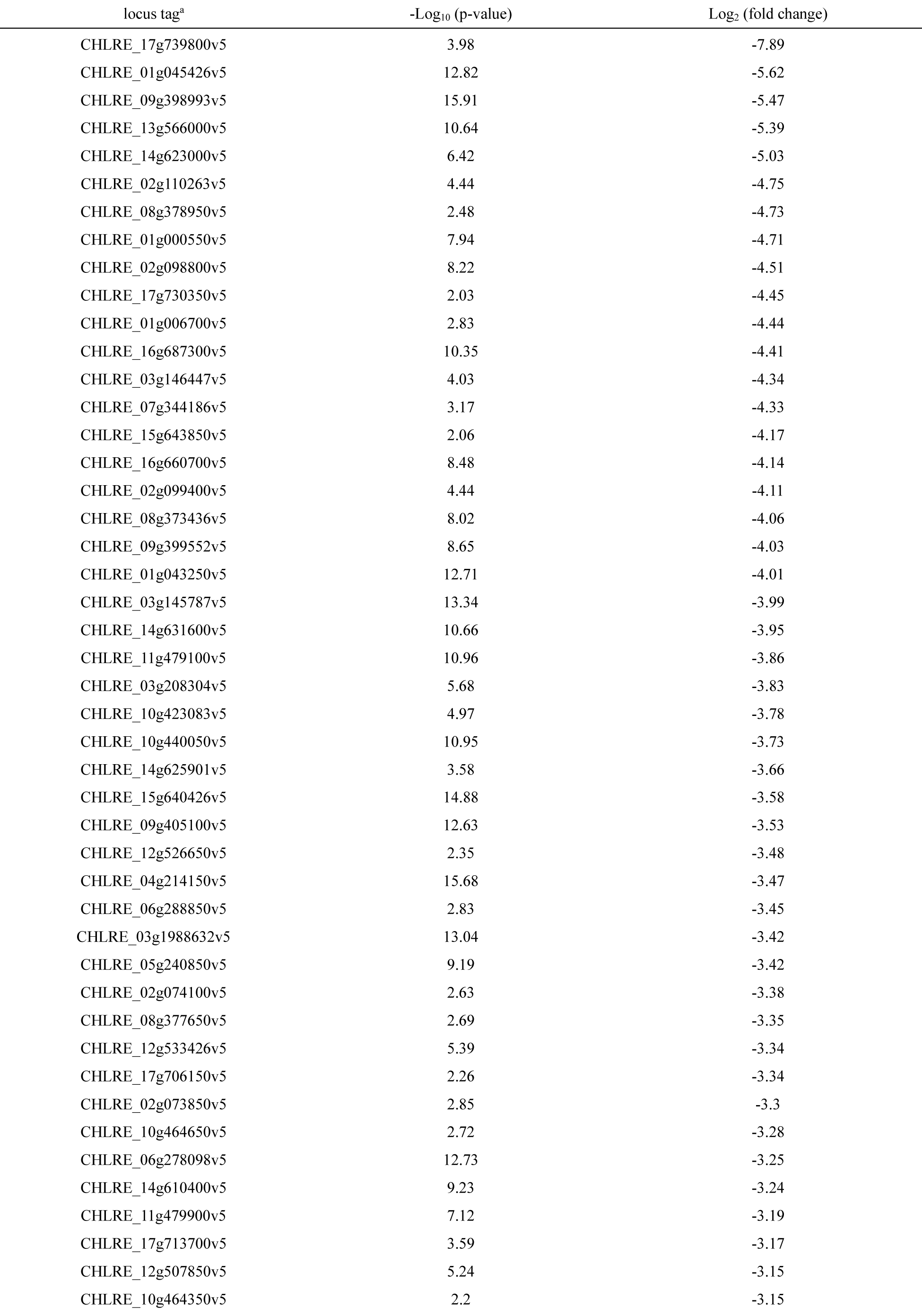
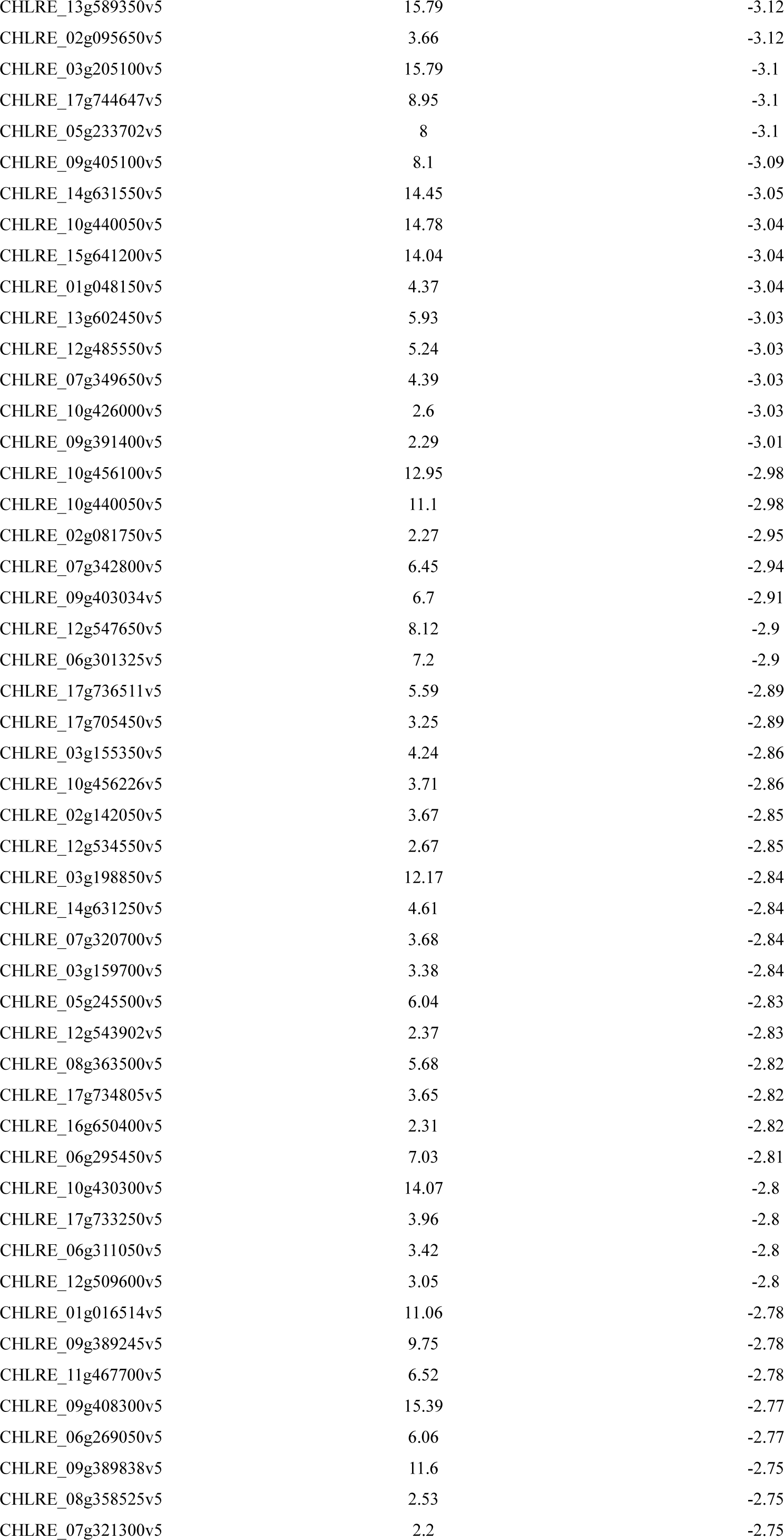
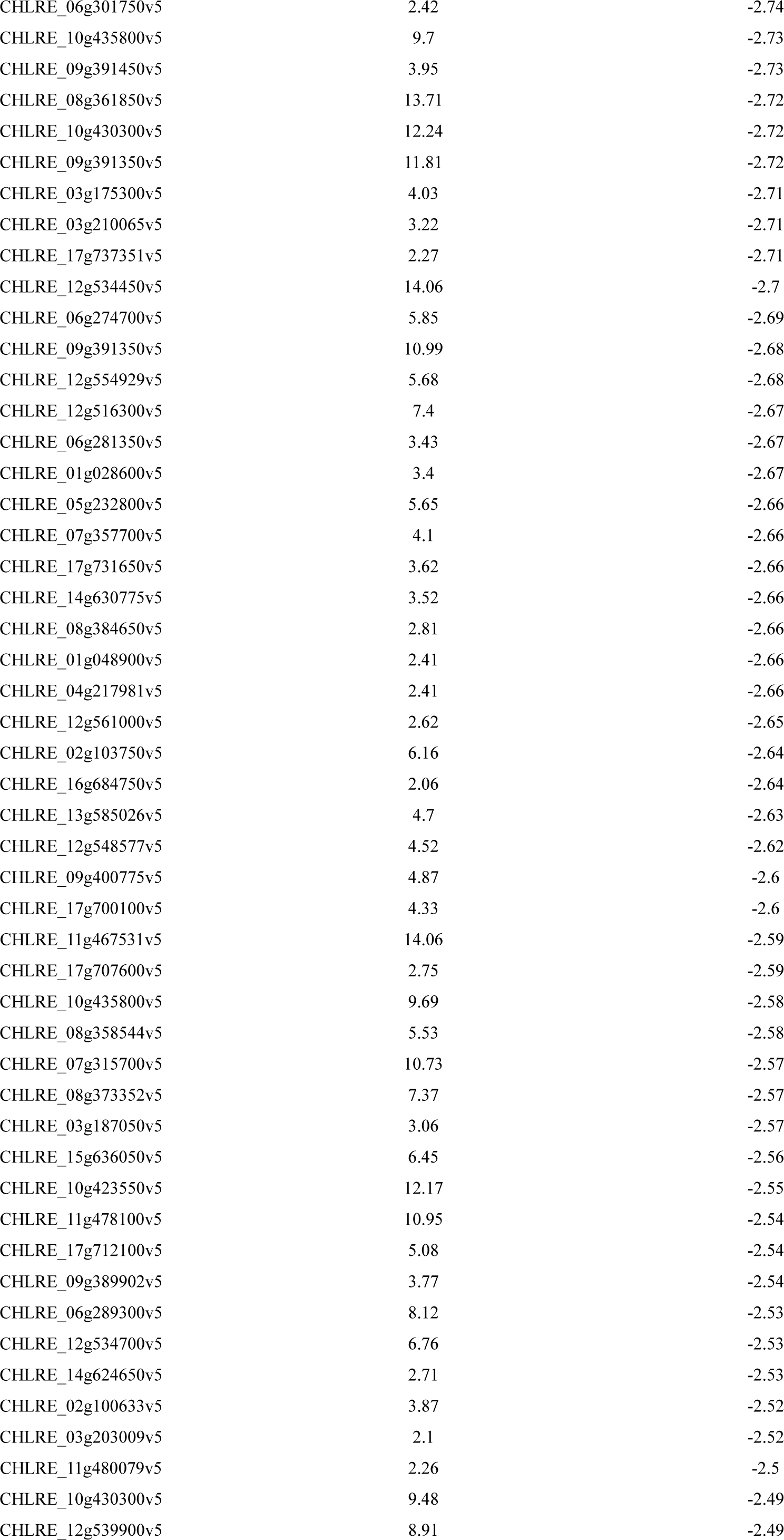
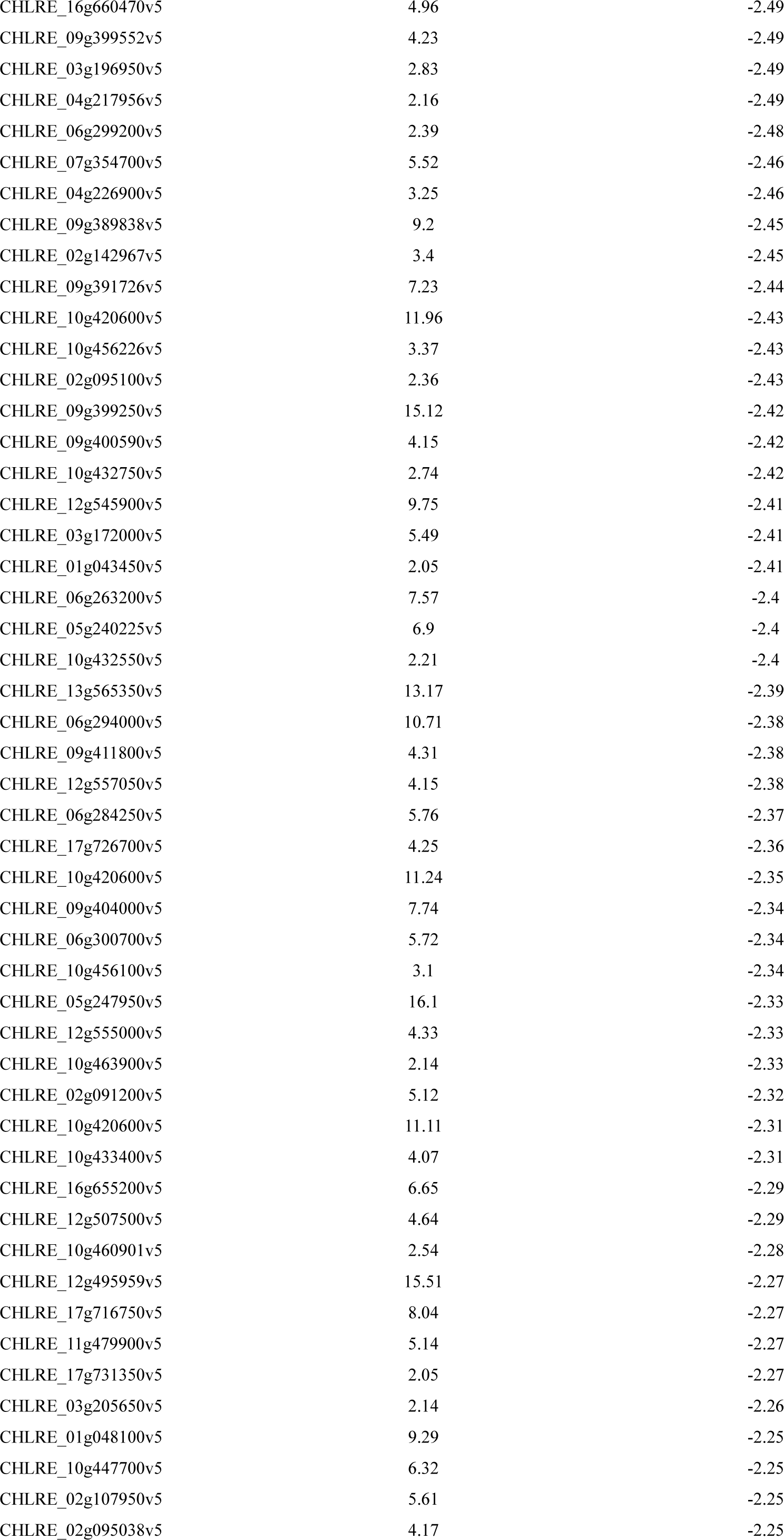
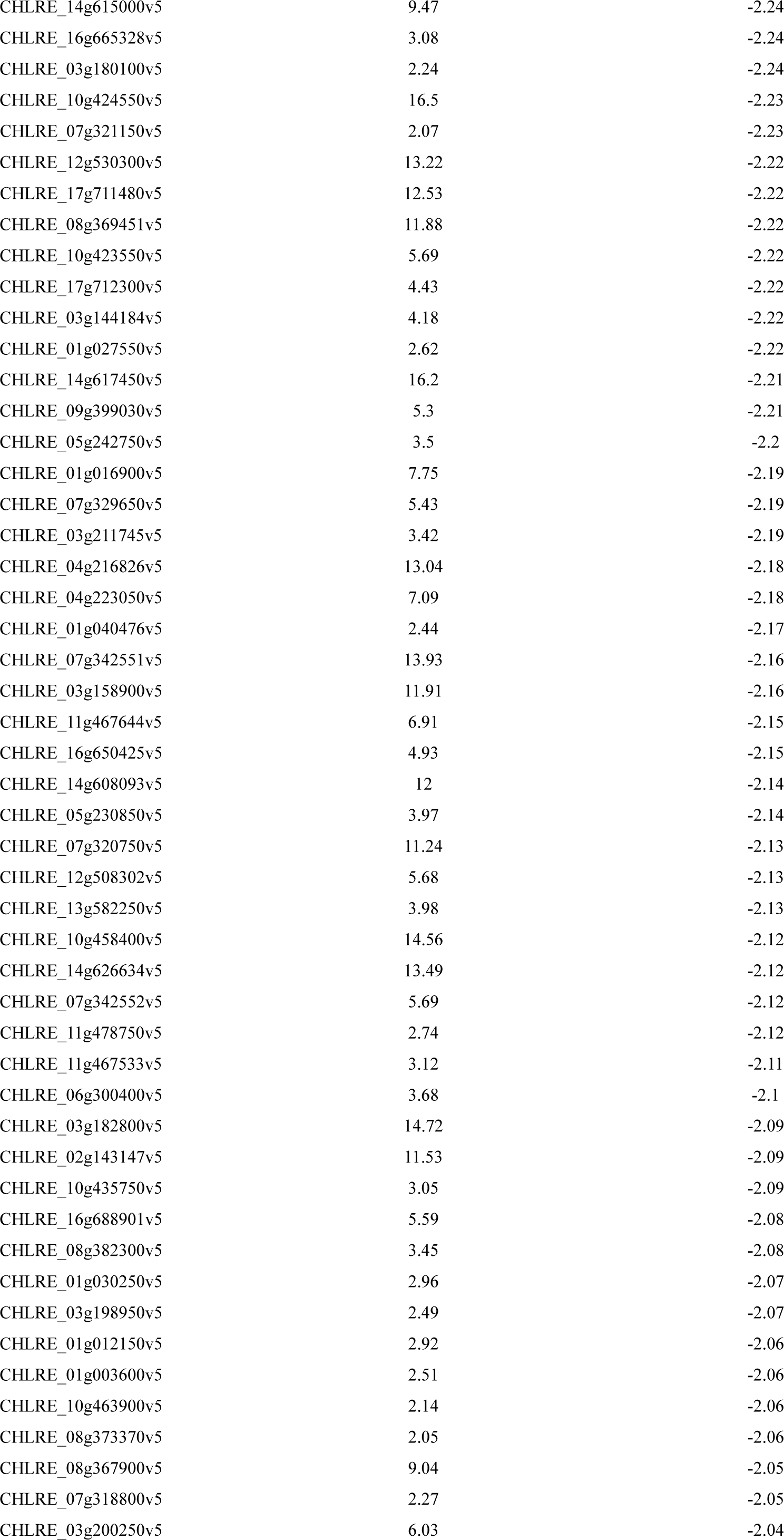
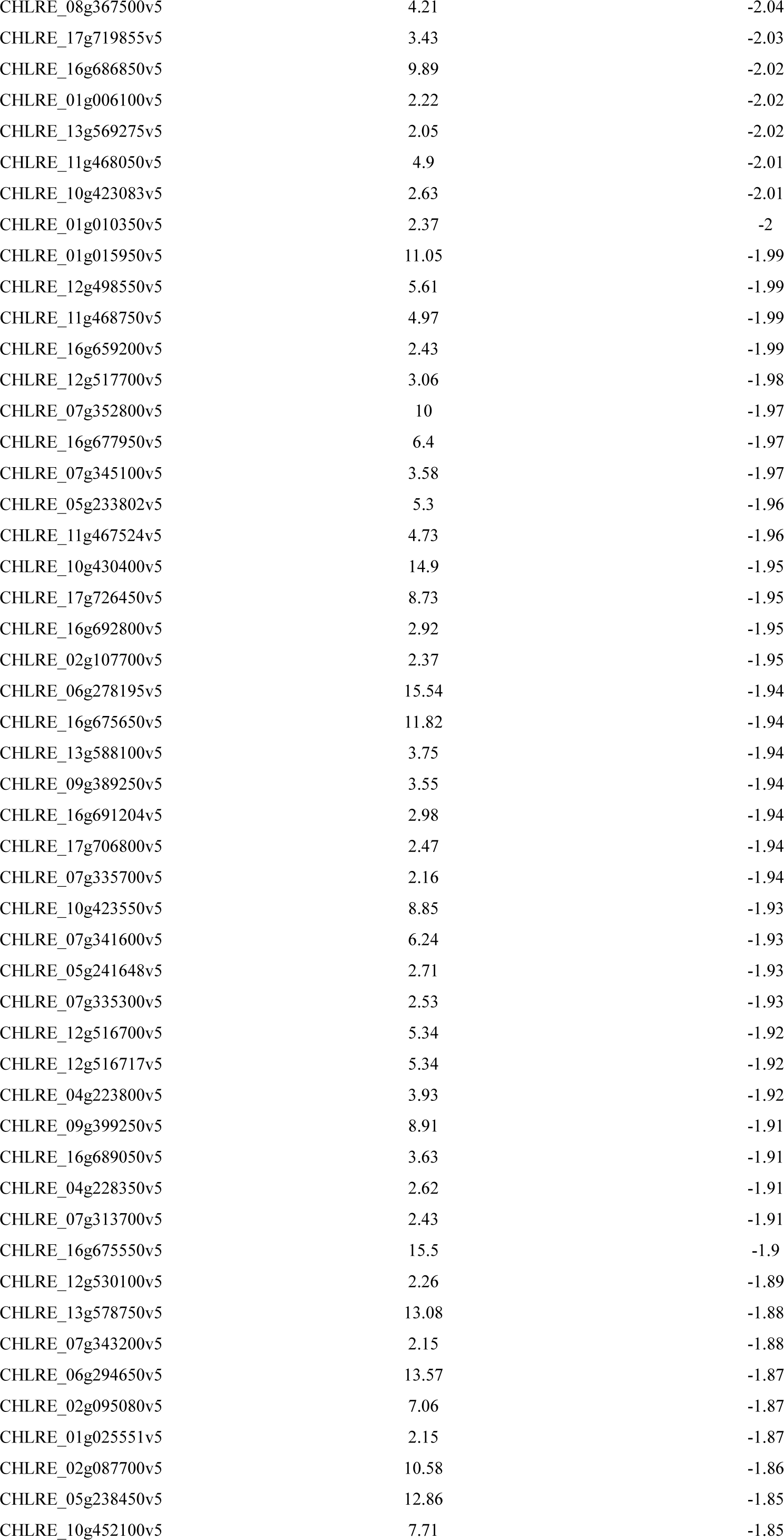
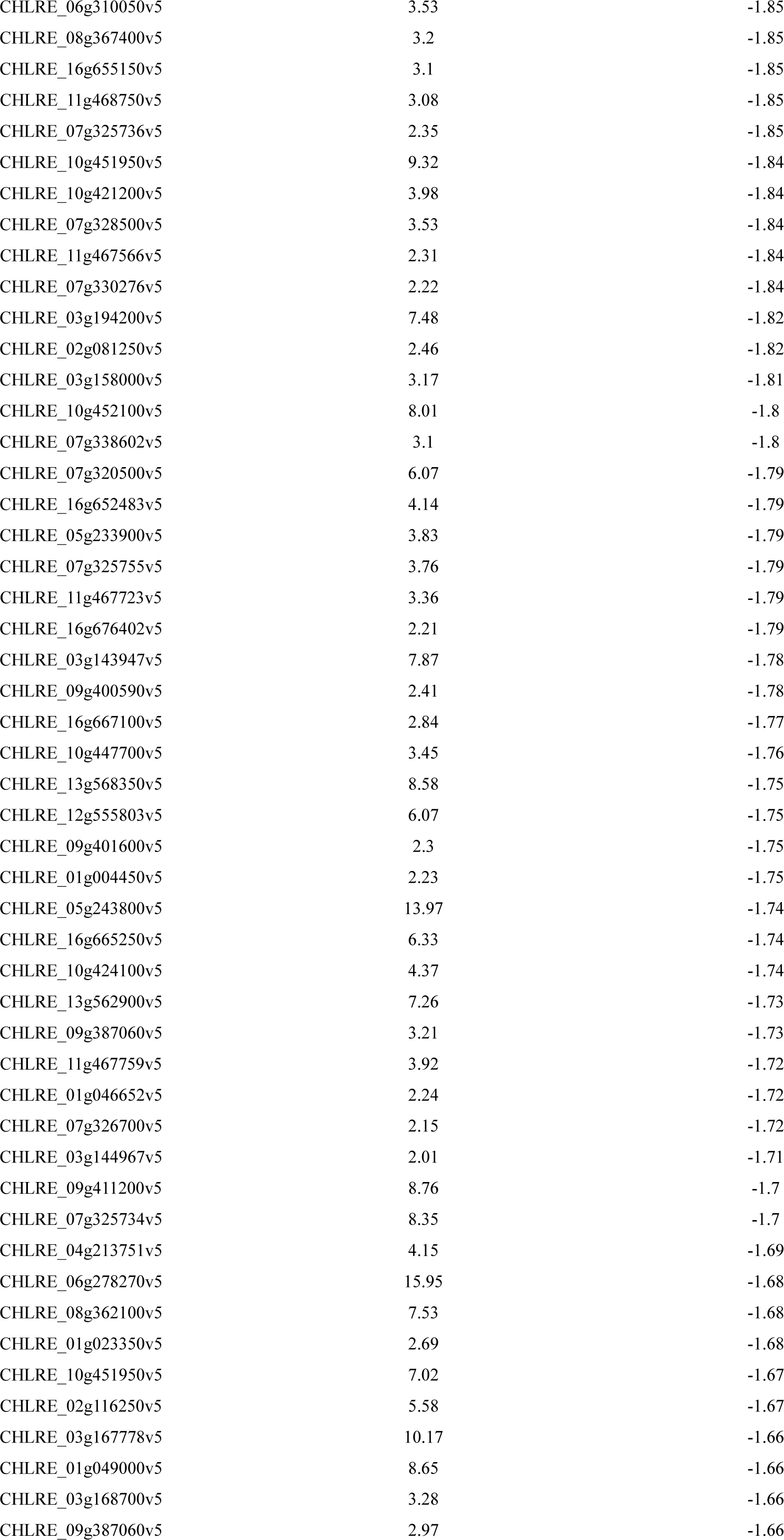
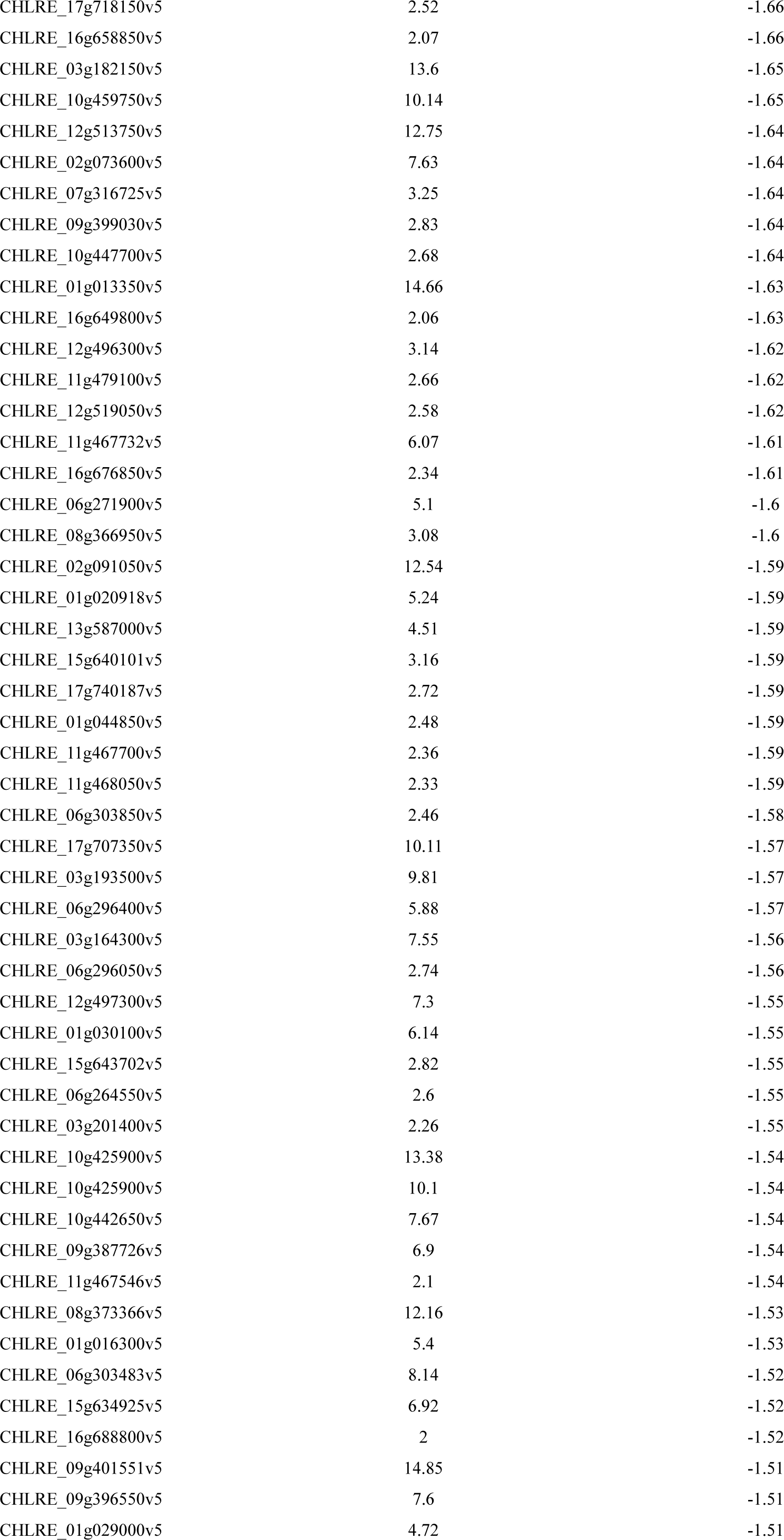
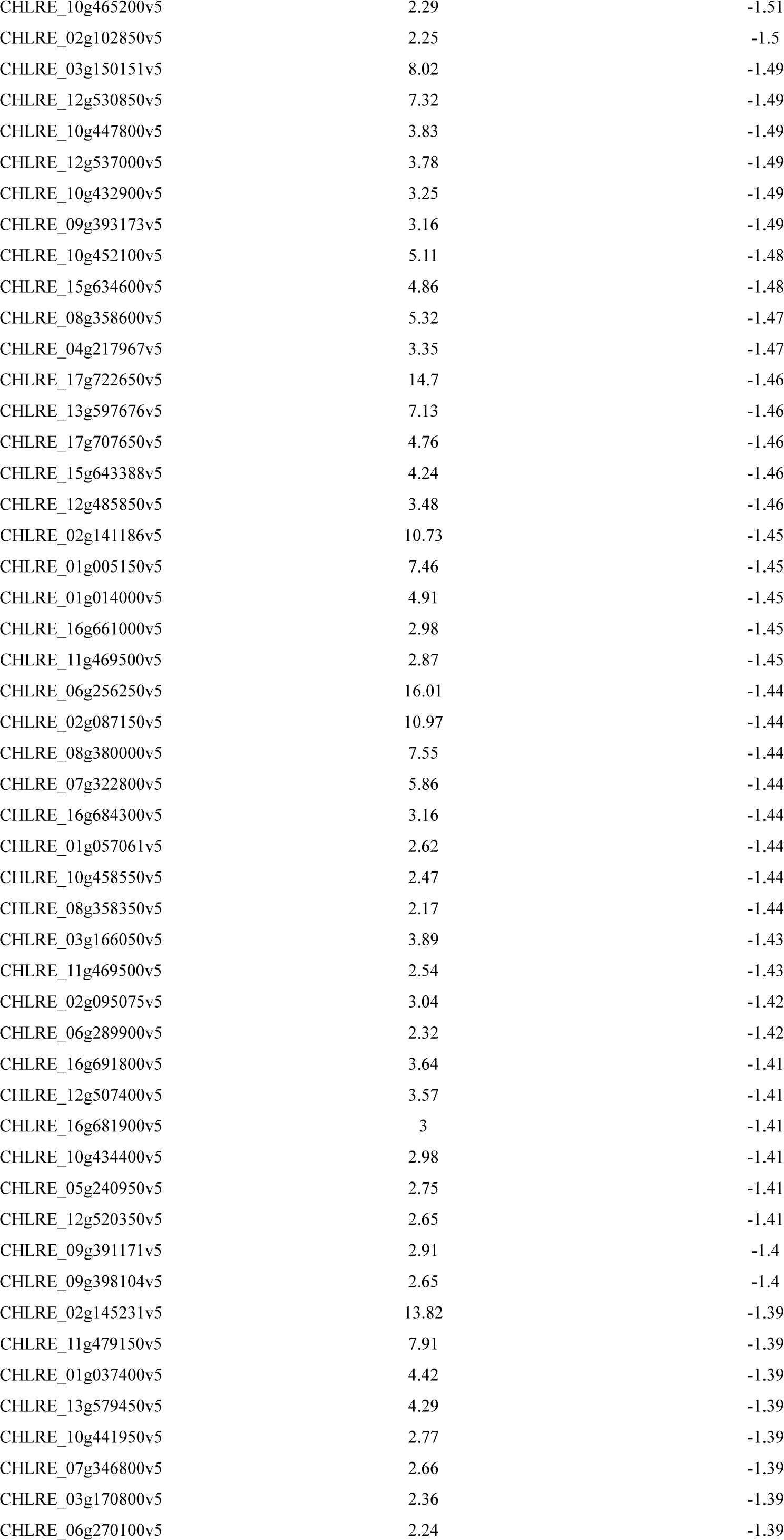
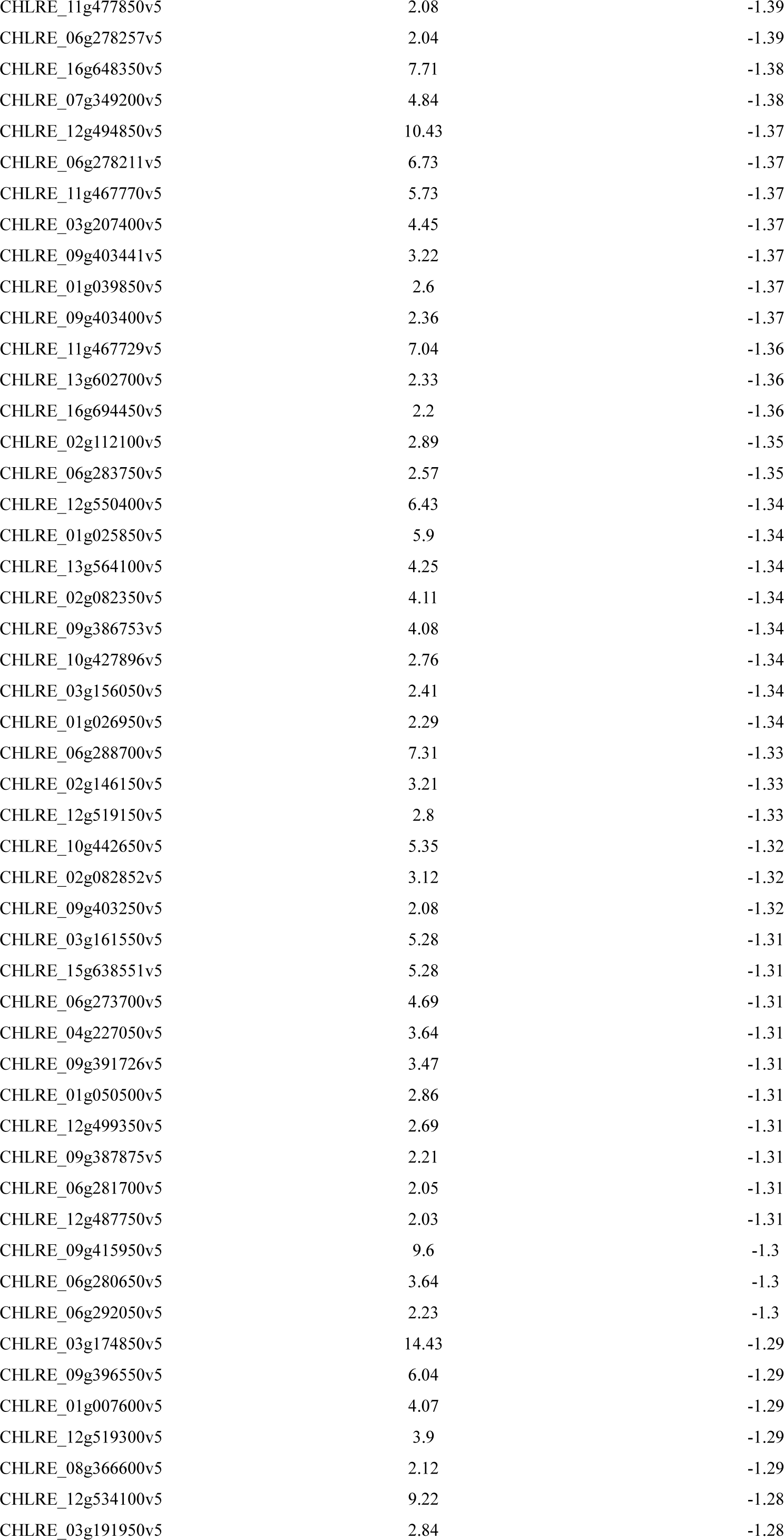
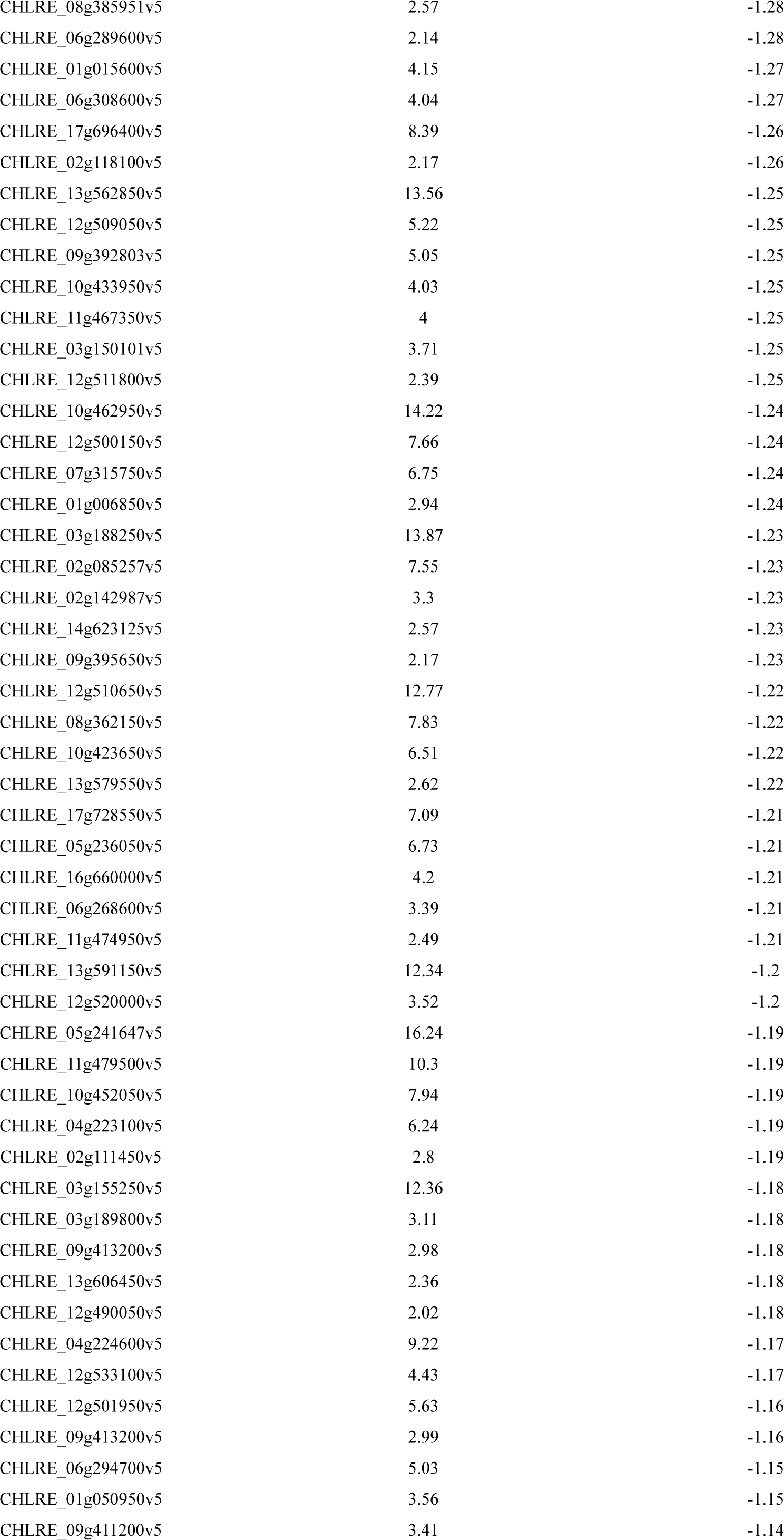
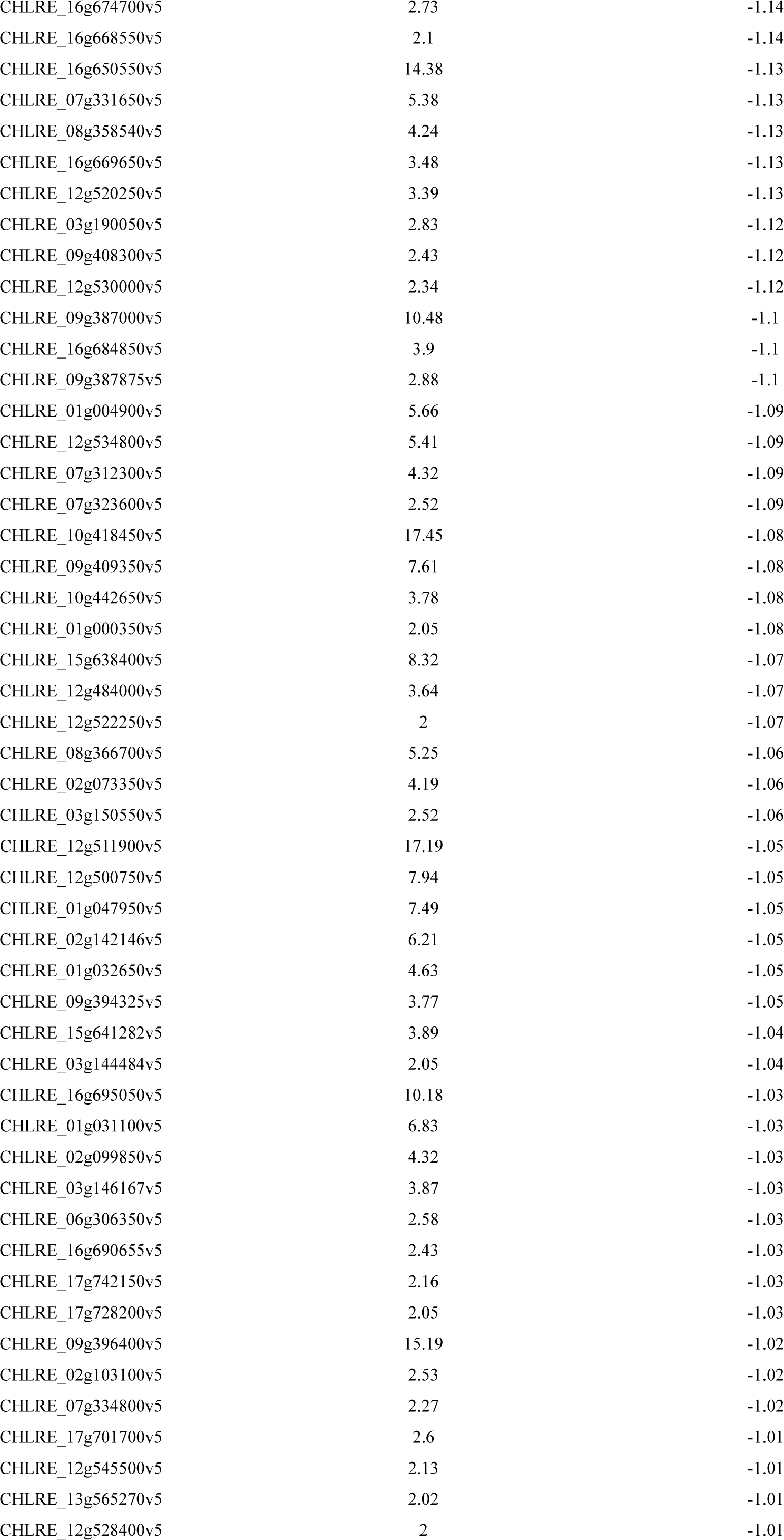
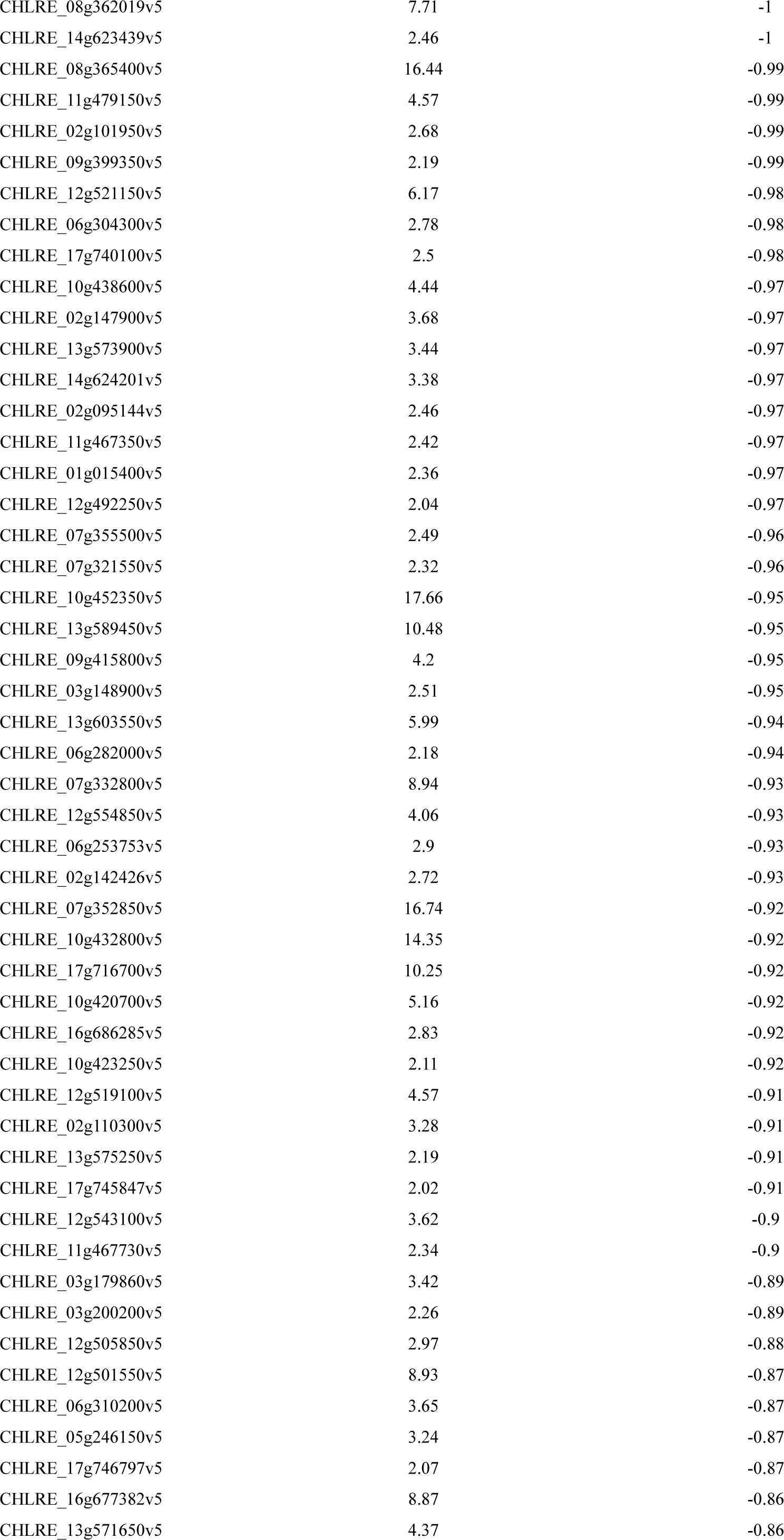
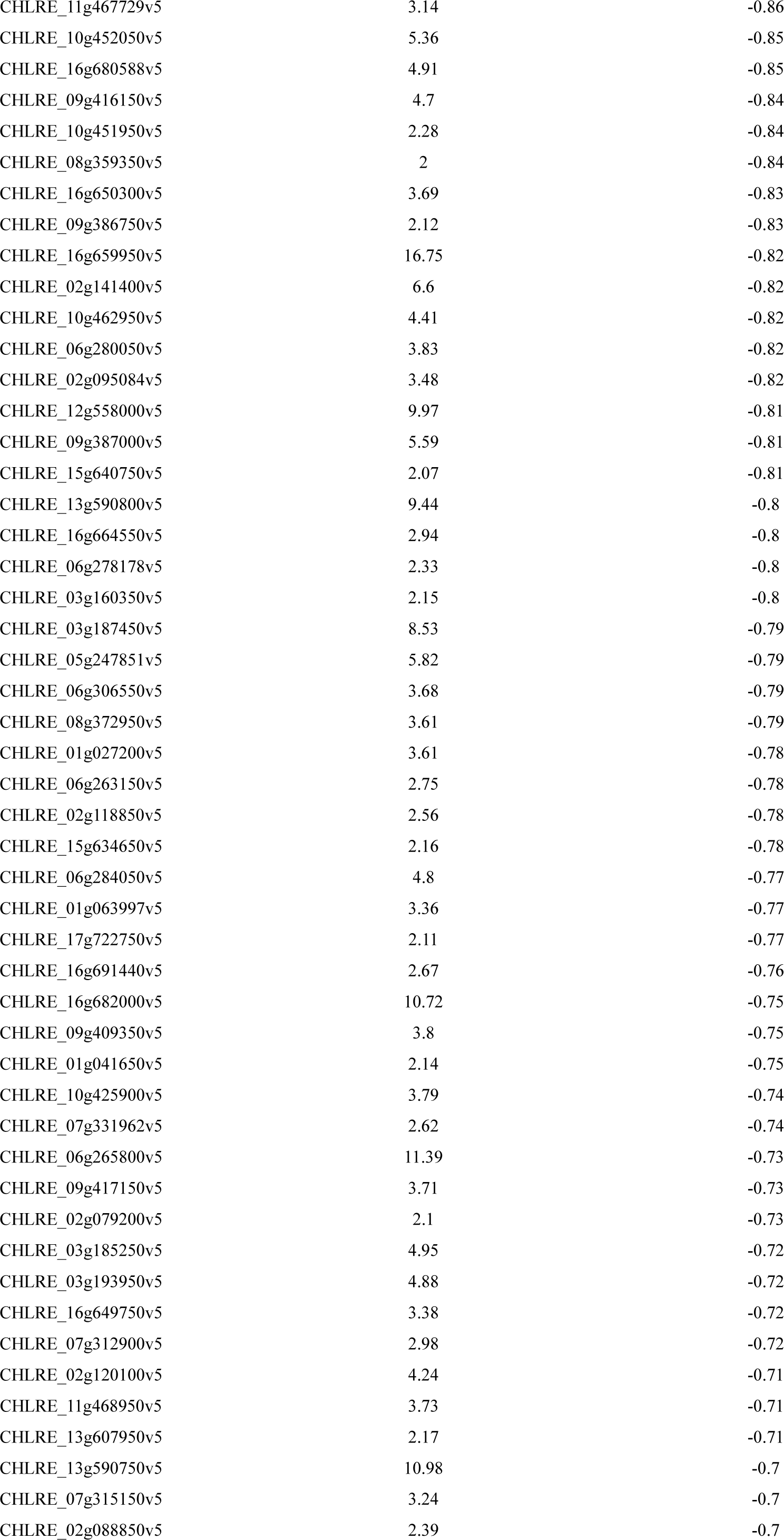
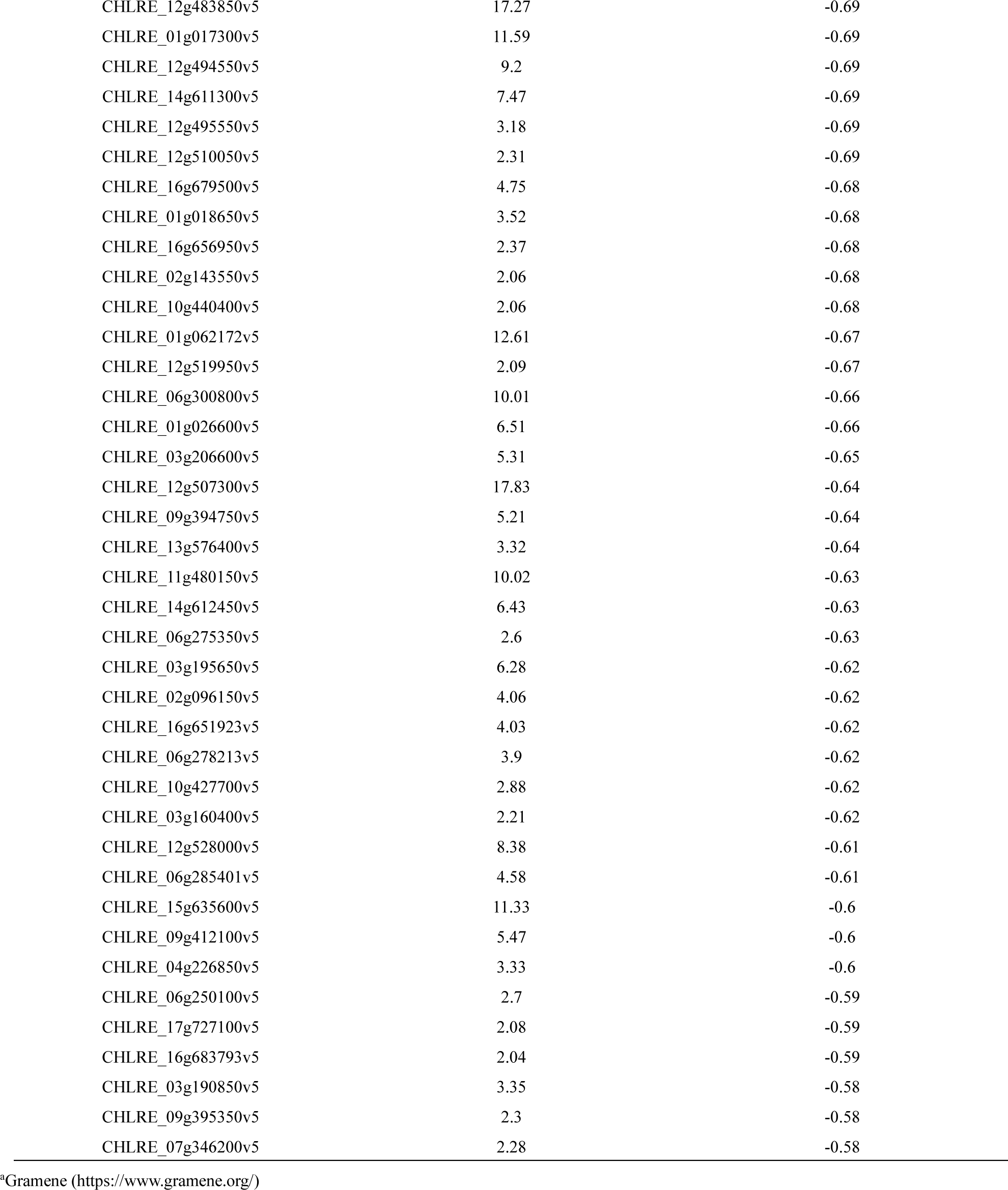
Microalgae genes whose expression levels decreased due to co-cultivation of microalgae and yeast.

| locus tag<sup>a</sup> | -Log<sub>10</sub> (p-value) | Log<sub>2</sub> (fold change) |
| --- | --- | --- |
| CHLRE_17g739800v5 | 3.98 | -7.89 |
| CHLRE_01g045426v5 | 12.82 | -5.62 |
| CHLRE_09g398993v5 | 15.91 | -5.47 |
| CHLRE_13g566000v5 | 10.64 | -5.39 |
| CHLRE_14g623000v5 | 6.42 | -5.03 |
| CHLRE_02g110263v5 | 4.44 | -4.75 |
| CHLRE_08g378950v5 | 2.48 | -4.73 |
| CHLRE_01g000550v5 | 7.94 | -4.71 |
| CHLRE_02g098800v5 | 8.22 | -4.51 |
| CHLRE_17g730350v5 | 2.03 | -4.45 |
| CHLRE_01g006700v5 | 2.83 | -4.44 |
| CHLRE_16g687300v5 | 10.35 | -4.41 |
| CHLRE_03g146447v5 | 4.03 | -4.34 |
| CHLRE_07g344186v5 | 3.17 | -4.33 |
| CHLRE_15g643850v5 | 2.06 | -4.17 |
| CHLRE_16g660700v5 | 8.48 | -4.14 |
| CHLRE_02g099400v5 | 4.44 | -4.11 |
| CHLRE_08g373436v5 | 8.02 | -4.06 |
| CHLRE_09g399552v5 | 8.65 | -4.03 |
| CHLRE_01g043250v5 | 12.71 | -4.01 |
| CHLRE_03g145787v5 | 13.34 | -3.99 |
| CHLRE_14g631600v5 | 10.66 | -3.95 |
| CHLRE_11g479100v5 | 10.96 | -3.86 |
| CHLRE_03g208304v5 | 5.68 | -3.83 |
| CHLRE_10g423083v5 | 4.97 | -3.78 |
| CHLRE_10g440050v5 | 10.95 | -3.73 |
| CHLRE_14g625901v5 | 3.58 | -3.66 |
| CHLRE_15g640426v5 | 14.88 | -3.58 |
| CHLRE_09g405100v5 | 12.63 | -3.53 |
| CHLRE_12g526650v5 | 2.35 | -3.48 |
| CHLRE_04g214150v5 | 15.68 | -3.47 |
| CHLRE_06g288850v5 | 2.83 | -3.45 |
| CHLRE_03g1988632v5 | 13.04 | -3.42 |
| CHLRE_05g240850v5 | 9.19 | -3.42 |
| CHLRE_02g074100v5 | 2.63 | -3.38 |
| CHLRE_08g377650v5 | 2.69 | -3.35 |
| CHLRE_12g533426v5 | 5.39 | -3.34 |
| CHLRE_17g706150v5 | 2.26 | -3.34 |
| CHLRE_02g073850v5 | 2.85 | -3.3 |
| CHLRE_10g464650v5 | 2.72 | -3.28 |
| CHLRE_06g278098v5 | 12.73 | -3.25 |
| CHLRE_14g610400v5 | 9.23 | -3.24 |
| CHLRE_11g479900v5 | 7.12 | -3.19 |
| CHLRE_17g713700v5 | 3.59 | -3.17 |
| CHLRE_12g507850v5 | 5.24 | -3.15 |
| CHLRE_10g464350v5 | 2.2 | -3.15 |
| CHLRE_13g589350v5 | 15.79 | -3.12 |
| CHLRE_02g095650v5 | 3.66 | -3.12 |
| CHLRE_03g205100v5 | 15.79 | -3.1 |
| CHLRE_17g744647v5 | 8.95 | -3.1 |
| CHLRE_05g233702v5 | 8 | -3.1 |
| CHLRE_09g405100v5 | 8.1 | -3.09 |
| CHLRE_14g631550v5 | 14.45 | -3.05 |
| CHLRE_10g440050v5 | 14.78 | -3.04 |
| CHLRE_15g641200v5 | 14.04 | -3.04 |
| CHLRE_01g048150v5 | 4.37 | -3.04 |
| CHLRE_13g602450v5 | 5.93 | -3.03 |
| CHLRE_12g485550v5 | 5.24 | -3.03 |
| CHLRE_07g349650v5 | 4.39 | -3.03 |
| CHLRE_10g426000v5 | 2.6 | -3.03 |
| CHLRE_09g391400v5 | 2.29 | -3.01 |
| CHLRE_10g456100v5 | 12.95 | -2.98 |
| CHLRE_10g440050v5 | 11.1 | -2.98 |
| CHLRE_02g081750v5 | 2.27 | -2.95 |
| CHLRE_07g342800v5 | 6.45 | -2.94 |
| CHLRE_09g403034v5 | 6.7 | -2.91 |
| CHLRE_12g547650v5 | 8.12 | -2.9 |
| CHLRE_06g301325v5 | 7.2 | -2.9 |
| CHLRE_17g736511v5 | 5.59 | -2.89 |
| CHLRE_17g705450v5 | 3.25 | -2.89 |
| CHLRE_03g155350v5 | 4.24 | -2.86 |
| CHLRE_10g456226v5 | 3.71 | -2.86 |
| CHLRE_02g142050v5 | 3.67 | -2.85 |
| CHLRE_12g534550v5 | 2.67 | -2.85 |
| CHLRE_03g198850v5 | 12.17 | -2.84 |
| CHLRE_14g631250v5 | 4.61 | -2.84 |
| CHLRE_07g320700v5 | 3.68 | -2.84 |
| CHLRE_03g159700v5 | 3.38 | -2.84 |
| CHLRE_05g245500v5 | 6.04 | -2.83 |
| CHLRE_12g543902v5 | 2.37 | -2.83 |
| CHLRE_08g363500v5 | 5.68 | -2.82 |
| CHLRE_17g734805v5 | 3.65 | -2.82 |
| CHLRE_16g650400v5 | 2.31 | -2.82 |
| CHLRE_06g295450v5 | 7.03 | -2.81 |
| CHLRE_10g430300v5 | 14.07 | -2.8 |
| CHLRE_17g733250v5 | 3.96 | -2.8 |
| CHLRE_06g311050v5 | 3.42 | -2.8 |
| CHLRE_12g509600v5 | 3.05 | -2.8 |
| CHLRE_01g016514v5 | 11.06 | -2.78 |
| CHLRE_09g389245v5 | 9.75 | -2.78 |
| CHLRE_11g467700v5 | 6.52 | -2.78 |
| CHLRE_09g408300v5 | 15.39 | -2.77 |
| CHLRE_06g269050v5 | 6.06 | -2.77 |
| CHLRE_09g389838v5 | 11.6 | -2.75 |
| CHLRE_08g358525v5 | 2.53 | -2.75 |
| CHLRE_07g321300v5 | 2.2 | -2.75 |
| CHLRE_06g301750v5 | 2.42 | -2.74 |
| CHLRE_10g435800v5 | 9.7 | -2.73 |
| CHLRE_09g391450v5 | 3.95 | -2.73 |
| CHLRE_08g361850v5 | 13.71 | -2.72 |
| CHLRE_10g430300v5 | 12.24 | -2.72 |
| CHLRE_09g391350v5 | 11.81 | -2.72 |
| CHLRE_03g175300v5 | 4.03 | -2.71 |
| CHLRE_03g210065v5 | 3.22 | -2.71 |
| CHLRE_17g737351v5 | 2.27 | -2.71 |
| CHLRE_12g534450v5 | 14.06 | -2.7 |
| CHLRE_06g274700v5 | 5.85 | -2.69 |
| CHLRE_09g391350v5 | 10.99 | -2.68 |
| CHLRE_12g554929v5 | 5.68 | -2.68 |
| CHLRE_12g516300v5 | 7.4 | -2.67 |
| CHLRE_06g281350v5 | 3.43 | -2.67 |
| CHLRE_01g028600v5 | 3.4 | -2.67 |
| CHLRE_05g232800v5 | 5.65 | -2.66 |
| CHLRE_07g357700v5 | 4.1 | -2.66 |
| CHLRE_17g731650v5 | 3.62 | -2.66 |
| CHLRE_14g630775v5 | 3.52 | -2.66 |
| CHLRE_08g384650v5 | 2.81 | -2.66 |
| CHLRE_01g048900v5 | 2.41 | -2.66 |
| CHLRE_04g217981v5 | 2.41 | -2.66 |
| CHLRE_12g561000v5 | 2.62 | -2.65 |
| CHLRE_02g103750v5 | 6.16 | -2.64 |
| CHLRE_16g684750v5 | 2.06 | -2.64 |
| CHLRE_13g585026v5 | 4.7 | -2.63 |
| CHLRE_12g548577v5 | 4.52 | -2.62 |
| CHLRE_09g400775v5 | 4.87 | -2.6 |
| CHLRE_17g700100v5 | 4.33 | -2.6 |
| CHLRE_11g467531v5 | 14.06 | -2.59 |
| CHLRE_17g707600v5 | 2.75 | -2.59 |
| CHLRE_10g435800v5 | 9.69 | -2.58 |
| CHLRE_08g358544v5 | 5.53 | -2.58 |
| CHLRE_07g315700v5 | 10.73 | -2.57 |
| CHLRE_08g373352v5 | 7.37 | -2.57 |
| CHLRE_03g187050v5 | 3.06 | -2.57 |
| CHLRE_15g636050v5 | 6.45 | -2.56 |
| CHLRE_10g423550v5 | 12.17 | -2.55 |
| CHLRE_11g478100v5 | 10.95 | -2.54 |
| CHLRE_17g712100v5 | 5.08 | -2.54 |
| CHLRE_09g389902v5 | 3.77 | -2.54 |
| CHLRE_06g289300v5 | 8.12 | -2.53 |
| CHLRE_12g534700v5 | 6.76 | -2.53 |
| CHLRE_14g624650v5 | 2.71 | -2.53 |
| CHLRE_02g100633v5 | 3.87 | -2.52 |
| CHLRE_03g203009v5 | 2.1 | -2.52 |
| CHLRE_11g480079v5 | 2.26 | -2.5 |
| CHLRE_10g430300v5 | 9.48 | -2.49 |
| CHLRE_12g539900v5 | 8.91 | -2.49 |
| CHLRE_16g660470v5 | 4.96 | -2.49 |
| CHLRE_09g399552v5 | 4.23 | -2.49 |
| CHLRE_03g196950v5 | 2.83 | -2.49 |
| CHLRE_04g217956v5 | 2.16 | -2.49 |
| CHLRE_06g299200v5 | 2.39 | -2.48 |
| CHLRE_07g354700v5 | 5.52 | -2.46 |
| CHLRE_04g226900v5 | 3.25 | -2.46 |
| CHLRE_09g389838v5 | 9.2 | -2.45 |
| CHLRE_02g142967v5 | 3.4 | -2.45 |
| CHLRE_09g391726v5 | 7.23 | -2.44 |
| CHLRE_10g420600v5 | 11.96 | -2.43 |
| CHLRE_10g456226v5 | 3.37 | -2.43 |
| CHLRE_02g095100v5 | 2.36 | -2.43 |
| CHLRE_09g399250v5 | 15.12 | -2.42 |
| CHLRE_09g400590v5 | 4.15 | -2.42 |
| CHLRE_10g432750v5 | 2.74 | -2.42 |
| CHLRE_12g545900v5 | 9.75 | -2.41 |
| CHLRE_03g172000v5 | 5.49 | -2.41 |
| CHLRE_01g043450v5 | 2.05 | -2.41 |
| CHLRE_06g263200v5 | 7.57 | -2.4 |
| CHLRE_05g240225v5 | 6.9 | -2.4 |
| CHLRE_10g432550v5 | 2.21 | -2.4 |
| CHLRE_13g565350v5 | 13.17 | -2.39 |
| CHLRE_06g294000v5 | 10.71 | -2.38 |
| CHLRE_09g411800v5 | 4.31 | -2.38 |
| CHLRE_12g557050v5 | 4.15 | -2.38 |
| CHLRE_06g284250v5 | 5.76 | -2.37 |
| CHLRE_17g726700v5 | 4.25 | -2.36 |
| CHLRE_10g420600v5 | 11.24 | -2.35 |
| CHLRE_09g404000v5 | 7.74 | -2.34 |
| CHLRE_06g300700v5 | 5.72 | -2.34 |
| CHLRE_10g456100v5 | 3.1 | -2.34 |
| CHLRE_05g247950v5 | 16.1 | -2.33 |
| CHLRE_12g555000v5 | 4.33 | -2.33 |
| CHLRE_10g463900v5 | 2.14 | -2.33 |
| CHLRE_02g091200v5 | 5.12 | -2.32 |
| CHLRE_10g420600v5 | 11.11 | -2.31 |
| CHLRE_10g433400v5 | 4.07 | -2.31 |
| CHLRE_16g655200v5 | 6.65 | -2.29 |
| CHLRE_12g507500v5 | 4.64 | -2.29 |
| CHLRE_10g460901v5 | 2.54 | -2.28 |
| CHLRE_12g495959v5 | 15.51 | -2.27 |
| CHLRE_17g716750v5 | 8.04 | -2.27 |
| CHLRE_11g479900v5 | 5.14 | -2.27 |
| CHLRE_17g731350v5 | 2.05 | -2.27 |
| CHLRE_03g205650v5 | 2.14 | -2.26 |
| CHLRE_01g048100v5 | 9.29 | -2.25 |
| CHLRE_10g447700v5 | 6.32 | -2.25 |
| CHLRE_02g107950v5 | 5.61 | -2.25 |
| CHLRE_02g095038v5 | 4.17 | -2.25 |
| CHLRE_14g615000v5 | 9.47 | -2.24 |
| CHLRE_16g665328v5 | 3.08 | -2.24 |
| CHLRE_03g180100v5 | 2.24 | -2.24 |
| CHLRE_10g424550v5 | 16.5 | -2.23 |
| CHLRE_07g321150v5 | 2.07 | -2.23 |
| CHLRE_12g530300v5 | 13.22 | -2.22 |
| CHLRE_17g711480v5 | 12.53 | -2.22 |
| CHLRE_08g369451v5 | 11.88 | -2.22 |
| CHLRE_10g423550v5 | 5.69 | -2.22 |
| CHLRE_17g712300v5 | 4.43 | -2.22 |
| CHLRE_03g144184v5 | 4.18 | -2.22 |
| CHLRE_01g027550v5 | 2.62 | -2.22 |
| CHLRE_14g617450v5 | 16.2 | -2.21 |
| CHLRE_09g399030v5 | 5.3 | -2.21 |
| CHLRE_05g242750v5 | 3.5 | -2.2 |
| CHLRE_01g016900v5 | 7.75 | -2.19 |
| CHLRE_07g329650v5 | 5.43 | -2.19 |
| CHLRE_03g211745v5 | 3.42 | -2.19 |
| CHLRE_04g216826v5 | 13.04 | -2.18 |
| CHLRE_04g223050v5 | 7.09 | -2.18 |
| CHLRE_01g040476v5 | 2.44 | -2.17 |
| CHLRE_07g342551v5 | 13.93 | -2.16 |
| CHLRE_03g158900v5 | 11.91 | -2.16 |
| CHLRE_11g467644v5 | 6.91 | -2.15 |
| CHLRE_16g650425v5 | 4.93 | -2.15 |
| CHLRE_14g608093v5 | 12 | -2.14 |
| CHLRE_05g230850v5 | 3.97 | -2.14 |
| CHLRE_07g320750v5 | 11.24 | -2.13 |
| CHLRE_12g508302v5 | 5.68 | -2.13 |
| CHLRE_13g582250v5 | 3.98 | -2.13 |
| CHLRE_10g458400v5 | 14.56 | -2.12 |
| CHLRE_14g626634v5 | 13.49 | -2.12 |
| CHLRE_07g342552v5 | 5.69 | -2.12 |
| CHLRE_11g478750v5 | 2.74 | -2.12 |
| CHLRE_11g467533v5 | 3.12 | -2.11 |
| CHLRE_06g300400v5 | 3.68 | -2.1 |
| CHLRE_03g182800v5 | 14.72 | -2.09 |
| CHLRE_02g143147v5 | 11.53 | -2.09 |
| CHLRE_10g435750v5 | 3.05 | -2.09 |
| CHLRE_16g688901v5 | 5.59 | -2.08 |
| CHLRE_08g382300v5 | 3.45 | -2.08 |
| CHLRE_01g030250v5 | 2.96 | -2.07 |
| CHLRE_03g198950v5 | 2.49 | -2.07 |
| CHLRE_01g012150v5 | 2.92 | -2.06 |
| CHLRE_01g003600v5 | 2.51 | -2.06 |
| CHLRE_10g463900v5 | 2.14 | -2.06 |
| CHLRE_08g373370v5 | 2.05 | -2.06 |
| CHLRE_08g367900v5 | 9.04 | -2.05 |
| CHLRE_07g318800v5 | 2.27 | -2.05 |
| CHLRE_03g200250v5 | 6.03 | -2.04 |
| CHLRE_08g367500v5 | 4.21 | -2.04 |
| CHLRE_17g719855v5 | 3.43 | -2.03 |
| CHLRE_16g686850v5 | 9.89 | -2.02 |
| CHLRE_01g006100v5 | 2.22 | -2.02 |
| CHLRE_13g569275v5 | 2.05 | -2.02 |
| CHLRE_11g468050v5 | 4.9 | -2.01 |
| CHLRE_10g423083v5 | 2.63 | -2.01 |
| CHLRE_01g010350v5 | 2.37 | -2 |
| CHLRE_01g015950v5 | 11.05 | -1.99 |
| CHLRE_12g498550v5 | 5.61 | -1.99 |
| CHLRE_11g468750v5 | 4.97 | -1.99 |
| CHLRE_16g659200v5 | 2.43 | -1.99 |
| CHLRE_12g517700v5 | 3.06 | -1.98 |
| CHLRE_07g352800v5 | 10 | -1.97 |
| CHLRE_16g677950v5 | 6.4 | -1.97 |
| CHLRE_07g345100v5 | 3.58 | -1.97 |
| CHLRE_05g233802v5 | 5.3 | -1.96 |
| CHLRE_11g467524v5 | 4.73 | -1.96 |
| CHLRE_10g430400v5 | 14.9 | -1.95 |
| CHLRE_17g726450v5 | 8.73 | -1.95 |
| CHLRE_16g692800v5 | 2.92 | -1.95 |
| CHLRE_02g107700v5 | 2.37 | -1.95 |
| CHLRE_06g278195v5 | 15.54 | -1.94 |
| CHLRE_16g675650v5 | 11.82 | -1.94 |
| CHLRE_13g588100v5 | 3.75 | -1.94 |
| CHLRE_09g389250v5 | 3.55 | -1.94 |
| CHLRE_16g691204v5 | 2.98 | -1.94 |
| CHLRE_17g706800v5 | 2.47 | -1.94 |
| CHLRE_07g335700v5 | 2.16 | -1.94 |
| CHLRE_10g423550v5 | 8.85 | -1.93 |
| CHLRE_07g341600v5 | 6.24 | -1.93 |
| CHLRE_05g241648v5 | 2.71 | -1.93 |
| CHLRE_07g335300v5 | 2.53 | -1.93 |
| CHLRE_12g516700v5 | 5.34 | -1.92 |
| CHLRE_12g516717v5 | 5.34 | -1.92 |
| CHLRE_04g223800v5 | 3.93 | -1.92 |
| CHLRE_09g399250v5 | 8.91 | -1.91 |
| CHLRE_16g689050v5 | 3.63 | -1.91 |
| CHLRE_04g228350v5 | 2.62 | -1.91 |
| CHLRE_07g313700v5 | 2.43 | -1.91 |
| CHLRE_16g675550v5 | 15.5 | -1.9 |
| CHLRE_12g530100v5 | 2.26 | -1.89 |
| CHLRE_13g578750v5 | 13.08 | -1.88 |
| CHLRE_07g343200v5 | 2.15 | -1.88 |
| CHLRE_06g294650v5 | 13.57 | -1.87 |
| CHLRE_02g095080v5 | 7.06 | -1.87 |
| CHLRE_01g025551v5 | 2.15 | -1.87 |
| CHLRE_02g087700v5 | 10.58 | -1.86 |
| CHLRE_05g238450v5 | 12.86 | -1.85 |
| CHLRE_10g452100v5 | 7.71 | -1.85 |
| CHLRE_06g310050v5 | 3.53 | -1.85 |
| CHLRE_08g367400v5 | 3.2 | -1.85 |
| CHLRE_16g655150v5 | 3.1 | -1.85 |
| CHLRE_11g468750v5 | 3.08 | -1.85 |
| CHLRE_07g325736v5 | 2.35 | -1.85 |
| CHLRE_10g451950v5 | 9.32 | -1.84 |
| CHLRE_10g421200v5 | 3.98 | -1.84 |
| CHLRE_07g328500v5 | 3.53 | -1.84 |
| CHLRE_11g467566v5 | 2.31 | -1.84 |
| CHLRE_07g330276v5 | 2.22 | -1.84 |
| CHLRE_03g194200v5 | 7.48 | -1.82 |
| CHLRE_02g081250v5 | 2.46 | -1.82 |
| CHLRE_03g158000v5 | 3.17 | -1.81 |
| CHLRE_10g452100v5 | 8.01 | -1.8 |
| CHLRE_07g338602v5 | 3.1 | -1.8 |
| CHLRE_07g320500v5 | 6.07 | -1.79 |
| CHLRE_16g652483v5 | 4.14 | -1.79 |
| CHLRE_05g233900v5 | 3.83 | -1.79 |
| CHLRE_07g325755v5 | 3.76 | -1.79 |
| CHLRE_11g467723v5 | 3.36 | -1.79 |
| CHLRE_16g676402v5 | 2.21 | -1.79 |
| CHLRE_03g143947v5 | 7.87 | -1.78 |
| CHLRE_09g400590v5 | 2.41 | -1.78 |
| CHLRE_16g667100v5 | 2.84 | -1.77 |
| CHLRE_10g447700v5 | 3.45 | -1.76 |
| CHLRE_13g568350v5 | 8.58 | -1.75 |
| CHLRE_12g555803v5 | 6.07 | -1.75 |
| CHLRE_09g401600v5 | 2.3 | -1.75 |
| CHLRE_01g004450v5 | 2.23 | -1.75 |
| CHLRE_05g243800v5 | 13.97 | -1.74 |
| CHLRE_16g665250v5 | 6.33 | -1.74 |
| CHLRE_10g424100v5 | 4.37 | -1.74 |
| CHLRE_13g562900v5 | 7.26 | -1.73 |
| CHLRE_09g387060v5 | 3.21 | -1.73 |
| CHLRE_11g467759v5 | 3.92 | -1.72 |
| CHLRE_01g046652v5 | 2.24 | -1.72 |
| CHLRE_07g326700v5 | 2.15 | -1.72 |
| CHLRE_03g144967v5 | 2.01 | -1.71 |
| CHLRE_09g411200v5 | 8.76 | -1.7 |
| CHLRE_07g325734v5 | 8.35 | -1.7 |
| CHLRE_04g213751v5 | 4.15 | -1.69 |
| CHLRE_06g278270v5 | 15.95 | -1.68 |
| CHLRE_08g362100v5 | 7.53 | -1.68 |
| CHLRE_01g023350v5 | 2.69 | -1.68 |
| CHLRE_10g451950v5 | 7.02 | -1.67 |
| CHLRE_02g116250v5 | 5.58 | -1.67 |
| CHLRE_03g167778v5 | 10.17 | -1.66 |
| CHLRE_01g049000v5 | 8.65 | -1.66 |
| CHLRE_03g168700v5 | 3.28 | -1.66 |
| CHLRE_09g387060v5 | 2.97 | -1.66 |
| CHLRE_17g718150v5 | 2.52 | -1.66 |
| CHLRE_16g658850v5 | 2.07 | -1.66 |
| CHLRE_03g182150v5 | 13.6 | -1.65 |
| CHLRE_10g459750v5 | 10.14 | -1.65 |
| CHLRE_12g513750v5 | 12.75 | -1.64 |
| CHLRE_02g073600v5 | 7.63 | -1.64 |
| CHLRE_07g316725v5 | 3.25 | -1.64 |
| CHLRE_09g399030v5 | 2.83 | -1.64 |
| CHLRE_10g447700v5 | 2.68 | -1.64 |
| CHLRE_01g013350v5 | 14.66 | -1.63 |
| CHLRE_16g649800v5 | 2.06 | -1.63 |
| CHLRE_12g496300v5 | 3.14 | -1.62 |
| CHLRE_11g479100v5 | 2.66 | -1.62 |
| CHLRE_12g519050v5 | 2.58 | -1.62 |
| CHLRE_11g467732v5 | 6.07 | -1.61 |
| CHLRE_16g676850v5 | 2.34 | -1.61 |
| CHLRE_06g271900v5 | 5.1 | -1.6 |
| CHLRE_08g366950v5 | 3.08 | -1.6 |
| CHLRE_02g091050v5 | 12.54 | -1.59 |
| CHLRE_01g020918v5 | 5.24 | -1.59 |
| CHLRE_13g587000v5 | 4.51 | -1.59 |
| CHLRE_15g640101v5 | 3.16 | -1.59 |
| CHLRE_17g740187v5 | 2.72 | -1.59 |
| CHLRE_01g044850v5 | 2.48 | -1.59 |
| CHLRE_11g467700v5 | 2.36 | -1.59 |
| CHLRE_11g468050v5 | 2.33 | -1.59 |
| CHLRE_06g303850v5 | 2.46 | -1.58 |
| CHLRE_17g707350v5 | 10.11 | -1.57 |
| CHLRE_03g193500v5 | 9.81 | -1.57 |
| CHLRE_06g296400v5 | 5.88 | -1.57 |
| CHLRE_03g164300v5 | 7.55 | -1.56 |
| CHLRE_06g296050v5 | 2.74 | -1.56 |
| CHLRE_12g497300v5 | 7.3 | -1.55 |
| CHLRE_01g030100v5 | 6.14 | -1.55 |
| CHLRE_15g643702v5 | 2.82 | -1.55 |
| CHLRE_06g264550v5 | 2.6 | -1.55 |
| CHLRE_03g201400v5 | 2.26 | -1.55 |
| CHLRE_10g425900v5 | 13.38 | -1.54 |
| CHLRE_10g425900v5 | 10.1 | -1.54 |
| CHLRE_10g442650v5 | 7.67 | -1.54 |
| CHLRE_09g387726v5 | 6.9 | -1.54 |
| CHLRE_11g467546v5 | 2.1 | -1.54 |
| CHLRE_08g373366v5 | 12.16 | -1.53 |
| CHLRE_01g016300v5 | 5.4 | -1.53 |
| CHLRE_06g303483v5 | 8.14 | -1.52 |
| CHLRE_15g634925v5 | 6.92 | -1.52 |
| CHLRE_16g688800v5 | 2 | -1.52 |
| CHLRE_09g401551v5 | 14.85 | -1.51 |
| CHLRE_09g396550v5 | 7.6 | -1.51 |
| CHLRE_01g029000v5 | 4.72 | -1.51 |
| CHLRE_10g465200v5 | 2.29 | -1.51 |
| CHLRE_02g102850v5 | 2.25 | -1.5 |
| CHLRE_03g150151v5 | 8.02 | -1.49 |
| CHLRE_12g530850v5 | 7.32 | -1.49 |
| CHLRE_10g447800v5 | 3.83 | -1.49 |
| CHLRE_12g537000v5 | 3.78 | -1.49 |
| CHLRE_10g432900v5 | 3.25 | -1.49 |
| CHLRE_09g393173v5 | 3.16 | -1.49 |
| CHLRE_10g452100v5 | 5.11 | -1.48 |
| CHLRE_15g634600v5 | 4.86 | -1.48 |
| CHLRE_08g358600v5 | 5.32 | -1.47 |
| CHLRE_04g217967v5 | 3.35 | -1.47 |
| CHLRE_17g722650v5 | 14.7 | -1.46 |
| CHLRE_13g597676v5 | 7.13 | -1.46 |
| CHLRE_17g707650v5 | 4.76 | -1.46 |
| CHLRE_15g643388v5 | 4.24 | -1.46 |
| CHLRE_12g485850v5 | 3.48 | -1.46 |
| CHLRE_02g141186v5 | 10.73 | -1.45 |
| CHLRE_01g005150v5 | 7.46 | -1.45 |
| CHLRE_01g014000v5 | 4.91 | -1.45 |
| CHLRE_16g661000v5 | 2.98 | -1.45 |
| CHLRE_11g469500v5 | 2.87 | -1.45 |
| CHLRE_06g256250v5 | 16.01 | -1.44 |
| CHLRE_02g087150v5 | 10.97 | -1.44 |
| CHLRE_08g380000v5 | 7.55 | -1.44 |
| CHLRE_07g322800v5 | 5.86 | -1.44 |
| CHLRE_16g684300v5 | 3.16 | -1.44 |
| CHLRE_01g057061v5 | 2.62 | -1.44 |
| CHLRE_10g458550v5 | 2.47 | -1.44 |
| CHLRE_08g358350v5 | 2.17 | -1.44 |
| CHLRE_03g166050v5 | 3.89 | -1.43 |
| CHLRE_11g469500v5 | 2.54 | -1.43 |
| CHLRE_02g095075v5 | 3.04 | -1.42 |
| CHLRE_06g289900v5 | 2.32 | -1.42 |
| CHLRE_16g691800v5 | 3.64 | -1.41 |
| CHLRE_12g507400v5 | 3.57 | -1.41 |
| CHLRE_16g681900v5 | 3 | -1.41 |
| CHLRE_10g434400v5 | 2.98 | -1.41 |
| CHLRE_05g240950v5 | 2.75 | -1.41 |
| CHLRE_12g520350v5 | 2.65 | -1.41 |
| CHLRE_09g391171v5 | 2.91 | -1.4 |
| CHLRE_09g398104v5 | 2.65 | -1.4 |
| CHLRE_02g145231v5 | 13.82 | -1.39 |
| CHLRE_11g479150v5 | 7.91 | -1.39 |
| CHLRE_01g037400v5 | 4.42 | -1.39 |
| CHLRE_13g579450v5 | 4.29 | -1.39 |
| CHLRE_10g441950v5 | 2.77 | -1.39 |
| CHLRE_07g346800v5 | 2.66 | -1.39 |
| CHLRE_03g170800v5 | 2.36 | -1.39 |
| CHLRE_06g270100v5 | 2.24 | -1.39 |
| CHLRE_11g477850v5 | 2.08 | -1.39 |
| CHLRE_06g278257v5 | 2.04 | -1.39 |
| CHLRE_16g648350v5 | 7.71 | -1.38 |
| CHLRE_07g349200v5 | 4.84 | -1.38 |
| CHLRE_12g494850v5 | 10.43 | -1.37 |
| CHLRE_06g278211v5 | 6.73 | -1.37 |
| CHLRE_11g467770v5 | 5.73 | -1.37 |
| CHLRE_03g207400v5 | 4.45 | -1.37 |
| CHLRE_09g403441v5 | 3.22 | -1.37 |
| CHLRE_01g039850v5 | 2.6 | -1.37 |
| CHLRE_09g403400v5 | 2.36 | -1.37 |
| CHLRE_11g467729v5 | 7.04 | -1.36 |
| CHLRE_13g602700v5 | 2.33 | -1.36 |
| CHLRE_16g694450v5 | 2.2 | -1.36 |
| CHLRE_02g112100v5 | 2.89 | -1.35 |
| CHLRE_06g283750v5 | 2.57 | -1.35 |
| CHLRE_12g550400v5 | 6.43 | -1.34 |
| CHLRE_01g025850v5 | 5.9 | -1.34 |
| CHLRE_13g564100v5 | 4.25 | -1.34 |
| CHLRE_02g082350v5 | 4.11 | -1.34 |
| CHLRE_09g386753v5 | 4.08 | -1.34 |
| CHLRE_10g427896v5 | 2.76 | -1.34 |
| CHLRE_03g156050v5 | 2.41 | -1.34 |
| CHLRE_01g026950v5 | 2.29 | -1.34 |
| CHLRE_06g288700v5 | 7.31 | -1.33 |
| CHLRE_02g146150v5 | 3.21 | -1.33 |
| CHLRE_12g519150v5 | 2.8 | -1.33 |
| CHLRE_10g442650v5 | 5.35 | -1.32 |
| CHLRE_02g082852v5 | 3.12 | -1.32 |
| CHLRE_09g403250v5 | 2.08 | -1.32 |
| CHLRE_03g161550v5 | 5.28 | -1.31 |
| CHLRE_15g638551v5 | 5.28 | -1.31 |
| CHLRE_06g273700v5 | 4.69 | -1.31 |
| CHLRE_04g227050v5 | 3.64 | -1.31 |
| CHLRE_09g391726v5 | 3.47 | -1.31 |
| CHLRE_01g050500v5 | 2.86 | -1.31 |
| CHLRE_12g499350v5 | 2.69 | -1.31 |
| CHLRE_09g387875v5 | 2.21 | -1.31 |
| CHLRE_06g281700v5 | 2.05 | -1.31 |
| CHLRE_12g487750v5 | 2.03 | -1.31 |
| CHLRE_09g415950v5 | 9.6 | -1.3 |
| CHLRE_06g280650v5 | 3.64 | -1.3 |
| CHLRE_06g292050v5 | 2.23 | -1.3 |
| CHLRE_03g174850v5 | 14.43 | -1.29 |
| CHLRE_09g396550v5 | 6.04 | -1.29 |
| CHLRE_01g007600v5 | 4.07 | -1.29 |
| CHLRE_12g519300v5 | 3.9 | -1.29 |
| CHLRE_08g366600v5 | 2.12 | -1.29 |
| CHLRE_12g534100v5 | 9.22 | -1.28 |
| CHLRE_03g191950v5 | 2.84 | -1.28 |
| CHLRE_08g385951v5 | 2.57 | -1.28 |
| CHLRE_06g289600v5 | 2.14 | -1.28 |
| CHLRE_01g015600v5 | 4.15 | -1.27 |
| CHLRE_06g308600v5 | 4.04 | -1.27 |
| CHLRE_17g696400v5 | 8.39 | -1.26 |
| CHLRE_02g118100v5 | 2.17 | -1.26 |
| CHLRE_13g562850v5 | 13.56 | -1.25 |
| CHLRE_12g509050v5 | 5.22 | -1.25 |
| CHLRE_09g392803v5 | 5.05 | -1.25 |
| CHLRE_10g433950v5 | 4.03 | -1.25 |
| CHLRE_11g467350v5 | 4 | -1.25 |
| CHLRE_03g150101v5 | 3.71 | -1.25 |
| CHLRE_12g511800v5 | 2.39 | -1.25 |
| CHLRE_10g462950v5 | 14.22 | -1.24 |
| CHLRE_12g500150v5 | 7.66 | -1.24 |
| CHLRE_07g315750v5 | 6.75 | -1.24 |
| CHLRE_01g006850v5 | 2.94 | -1.24 |
| CHLRE_03g188250v5 | 13.87 | -1.23 |
| CHLRE_02g085257v5 | 7.55 | -1.23 |
| CHLRE_02g142987v5 | 3.3 | -1.23 |
| CHLRE_14g623125v5 | 2.57 | -1.23 |
| CHLRE_09g395650v5 | 2.17 | -1.23 |
| CHLRE_12g510650v5 | 12.77 | -1.22 |
| CHLRE_08g362150v5 | 7.83 | -1.22 |
| CHLRE_10g423650v5 | 6.51 | -1.22 |
| CHLRE_13g579550v5 | 2.62 | -1.22 |
| CHLRE_17g728550v5 | 7.09 | -1.21 |
| CHLRE_05g236050v5 | 6.73 | -1.21 |
| CHLRE_16g660000v5 | 4.2 | -1.21 |
| CHLRE_06g268600v5 | 3.39 | -1.21 |
| CHLRE_11g474950v5 | 2.49 | -1.21 |
| CHLRE_13g591150v5 | 12.34 | -1.2 |
| CHLRE_12g520000v5 | 3.52 | -1.2 |
| CHLRE_05g241647v5 | 16.24 | -1.19 |
| CHLRE_11g479500v5 | 10.3 | -1.19 |
| CHLRE_10g452050v5 | 7.94 | -1.19 |
| CHLRE_04g223100v5 | 6.24 | -1.19 |
| CHLRE_02g111450v5 | 2.8 | -1.19 |
| CHLRE_03g155250v5 | 12.36 | -1.18 |
| CHLRE_03g189800v5 | 3.11 | -1.18 |
| CHLRE_09g413200v5 | 2.98 | -1.18 |
| CHLRE_13g606450v5 | 2.36 | -1.18 |
| CHLRE_12g490050v5 | 2.02 | -1.18 |
| CHLRE_04g224600v5 | 9.22 | -1.17 |
| CHLRE_12g533100v5 | 4.43 | -1.17 |
| CHLRE_12g501950v5 | 5.63 | -1.16 |
| CHLRE_09g413200v5 | 2.99 | -1.16 |
| CHLRE_06g294700v5 | 5.03 | -1.15 |
| CHLRE_01g050950v5 | 3.56 | -1.15 |
| CHLRE_09g411200v5 | 3.41 | -1.14 |
| CHLRE_16g674700v5 | 2.73 | -1.14 |
| CHLRE_16g668550v5 | 2.1 | -1.14 |
| CHLRE_16g650550v5 | 14.38 | -1.13 |
| CHLRE_07g331650v5 | 5.38 | -1.13 |
| CHLRE_08g358540v5 | 4.24 | -1.13 |
| CHLRE_16g669650v5 | 3.48 | -1.13 |
| CHLRE_12g520250v5 | 3.39 | -1.13 |
| CHLRE_03g190050v5 | 2.83 | -1.12 |
| CHLRE_09g408300v5 | 2.43 | -1.12 |
| CHLRE_12g530000v5 | 2.34 | -1.12 |
| CHLRE_09g387000v5 | 10.48 | -1.1 |
| CHLRE_16g684850v5 | 3.9 | -1.1 |
| CHLRE_09g387875v5 | 2.88 | -1.1 |
| CHLRE_01g004900v5 | 5.66 | -1.09 |
| CHLRE_12g534800v5 | 5.41 | -1.09 |
| CHLRE_07g312300v5 | 4.32 | -1.09 |
| CHLRE_07g323600v5 | 2.52 | -1.09 |
| CHLRE_10g418450v5 | 17.45 | -1.08 |
| CHLRE_09g409350v5 | 7.61 | -1.08 |
| CHLRE_10g442650v5 | 3.78 | -1.08 |
| CHLRE_01g000350v5 | 2.05 | -1.08 |
| CHLRE_15g638400v5 | 8.32 | -1.07 |
| CHLRE_12g484000v5 | 3.64 | -1.07 |
| CHLRE_12g522250v5 | 2 | -1.07 |
| CHLRE_08g366700v5 | 5.25 | -1.06 |
| CHLRE_02g073350v5 | 4.19 | -1.06 |
| CHLRE_03g150550v5 | 2.52 | -1.06 |
| CHLRE_12g511900v5 | 17.19 | -1.05 |
| CHLRE_12g500750v5 | 7.94 | -1.05 |
| CHLRE_01g047950v5 | 7.49 | -1.05 |
| CHLRE_02g142146v5 | 6.21 | -1.05 |
| CHLRE_01g032650v5 | 4.63 | -1.05 |
| CHLRE_09g394325v5 | 3.77 | -1.05 |
| CHLRE_15g641282v5 | 3.89 | -1.04 |
| CHLRE_03g144484v5 | 2.05 | -1.04 |
| CHLRE_16g695050v5 | 10.18 | -1.03 |
| CHLRE_01g031100v5 | 6.83 | -1.03 |
| CHLRE_02g099850v5 | 4.32 | -1.03 |
| CHLRE_03g146167v5 | 3.87 | -1.03 |
| CHLRE_06g306350v5 | 2.58 | -1.03 |
| CHLRE_16g690655v5 | 2.43 | -1.03 |
| CHLRE_17g742150v5 | 2.16 | -1.03 |
| CHLRE_17g728200v5 | 2.05 | -1.03 |
| CHLRE_09g396400v5 | 15.19 | -1.02 |
| CHLRE_02g103100v5 | 2.53 | -1.02 |
| CHLRE_07g334800v5 | 2.27 | -1.02 |
| CHLRE_17g701700v5 | 2.6 | -1.01 |
| CHLRE_12g545500v5 | 2.13 | -1.01 |
| CHLRE_13g565270v5 | 2.02 | -1.01 |
| CHLRE_12g528400v5 | 2 | -1.01 |
| CHLRE_08g362019v5 | 7.71 | -1 |
| CHLRE_14g623439v5 | 2.46 | -1 |
| CHLRE_08g365400v5 | 16.44 | -0.99 |
| CHLRE_11g479150v5 | 4.57 | -0.99 |
| CHLRE_02g101950v5 | 2.68 | -0.99 |
| CHLRE_09g399350v5 | 2.19 | -0.99 |
| CHLRE_12g521150v5 | 6.17 | -0.98 |
| CHLRE_06g304300v5 | 2.78 | -0.98 |
| CHLRE_17g740100v5 | 2.5 | -0.98 |
| CHLRE_10g438600v5 | 4.44 | -0.97 |
| CHLRE_02g147900v5 | 3.68 | -0.97 |
| CHLRE_13g573900v5 | 3.44 | -0.97 |
| CHLRE_14g624201v5 | 3.38 | -0.97 |
| CHLRE_02g095144v5 | 2.46 | -0.97 |
| CHLRE_11g467350v5 | 2.42 | -0.97 |
| CHLRE_01g015400v5 | 2.36 | -0.97 |
| CHLRE_12g492250v5 | 2.04 | -0.97 |
| CHLRE_07g355500v5 | 2.49 | -0.96 |
| CHLRE_07g321550v5 | 2.32 | -0.96 |
| CHLRE_10g452350v5 | 17.66 | -0.95 |
| CHLRE_13g589450v5 | 10.48 | -0.95 |
| CHLRE_09g415800v5 | 4.2 | -0.95 |
| CHLRE_03g148900v5 | 2.51 | -0.95 |
| CHLRE_13g603550v5 | 5.99 | -0.94 |
| CHLRE_06g282000v5 | 2.18 | -0.94 |
| CHLRE_07g332800v5 | 8.94 | -0.93 |
| CHLRE_12g554850v5 | 4.06 | -0.93 |
| CHLRE_06g253753v5 | 2.9 | -0.93 |
| CHLRE_02g142426v5 | 2.72 | -0.93 |
| CHLRE_07g352850v5 | 16.74 | -0.92 |
| CHLRE_10g432800v5 | 14.35 | -0.92 |
| CHLRE_17g716700v5 | 10.25 | -0.92 |
| CHLRE_10g420700v5 | 5.16 | -0.92 |
| CHLRE_16g686285v5 | 2.83 | -0.92 |
| CHLRE_10g423250v5 | 2.11 | -0.92 |
| CHLRE_12g519100v5 | 4.57 | -0.91 |
| CHLRE_02g110300v5 | 3.28 | -0.91 |
| CHLRE_13g575250v5 | 2.19 | -0.91 |
| CHLRE_17g745847v5 | 2.02 | -0.91 |
| CHLRE_12g543100v5 | 3.62 | -0.9 |
| CHLRE_11g467730v5 | 2.34 | -0.9 |
| CHLRE_03g179860v5 | 3.42 | -0.89 |
| CHLRE_03g200200v5 | 2.26 | -0.89 |
| CHLRE_12g505850v5 | 2.97 | -0.88 |
| CHLRE_12g501550v5 | 8.93 | -0.87 |
| CHLRE_06g310200v5 | 3.65 | -0.87 |
| CHLRE_05g246150v5 | 3.24 | -0.87 |
| CHLRE_17g746797v5 | 2.07 | -0.87 |
| CHLRE_16g677382v5 | 8.87 | -0.86 |
| CHLRE_13g571650v5 | 4.37 | -0.86 |
| CHLRE_11g467729v5 | 3.14 | -0.86 |
| CHLRE_10g452050v5 | 5.36 | -0.85 |
| CHLRE_16g680588v5 | 4.91 | -0.85 |
| CHLRE_09g416150v5 | 4.7 | -0.84 |
| CHLRE_10g451950v5 | 2.28 | -0.84 |
| CHLRE_08g359350v5 | 2 | -0.84 |
| CHLRE_16g650300v5 | 3.69 | -0.83 |
| CHLRE_09g386750v5 | 2.12 | -0.83 |
| CHLRE_16g659950v5 | 16.75 | -0.82 |
| CHLRE_02g141400v5 | 6.6 | -0.82 |
| CHLRE_10g462950v5 | 4.41 | -0.82 |
| CHLRE_06g280050v5 | 3.83 | -0.82 |
| CHLRE_02g095084v5 | 3.48 | -0.82 |
| CHLRE_12g558000v5 | 9.97 | -0.81 |
| CHLRE_09g387000v5 | 5.59 | -0.81 |
| CHLRE_15g640750v5 | 2.07 | -0.81 |
| CHLRE_13g590800v5 | 9.44 | -0.8 |
| CHLRE_16g664550v5 | 2.94 | -0.8 |
| CHLRE_06g278178v5 | 2.33 | -0.8 |
| CHLRE_03g160350v5 | 2.15 | -0.8 |
| CHLRE_03g187450v5 | 8.53 | -0.79 |
| CHLRE_05g247851v5 | 5.82 | -0.79 |
| CHLRE_06g306550v5 | 3.68 | -0.79 |
| CHLRE_08g372950v5 | 3.61 | -0.79 |
| CHLRE_01g027200v5 | 3.61 | -0.78 |
| CHLRE_06g263150v5 | 2.75 | -0.78 |
| CHLRE_02g118850v5 | 2.56 | -0.78 |
| CHLRE_15g634650v5 | 2.16 | -0.78 |
| CHLRE_06g284050v5 | 4.8 | -0.77 |
| CHLRE_01g063997v5 | 3.36 | -0.77 |
| CHLRE_17g722750v5 | 2.11 | -0.77 |
| CHLRE_16g691440v5 | 2.67 | -0.76 |
| CHLRE_16g682000v5 | 10.72 | -0.75 |
| CHLRE_09g409350v5 | 3.8 | -0.75 |
| CHLRE_01g041650v5 | 2.14 | -0.75 |
| CHLRE_10g425900v5 | 3.79 | -0.74 |
| CHLRE_07g331962v5 | 2.62 | -0.74 |
| CHLRE_06g265800v5 | 11.39 | -0.73 |
| CHLRE_09g417150v5 | 3.71 | -0.73 |
| CHLRE_02g079200v5 | 2.1 | -0.73 |
| CHLRE_03g185250v5 | 4.95 | -0.72 |
| CHLRE_03g193950v5 | 4.88 | -0.72 |
| CHLRE_16g649750v5 | 3.38 | -0.72 |
| CHLRE_07g312900v5 | 2.98 | -0.72 |
| CHLRE_02g120100v5 | 4.24 | -0.71 |
| CHLRE_11g468950v5 | 3.73 | -0.71 |
| CHLRE_13g607950v5 | 2.17 | -0.71 |
| CHLRE_13g590750v5 | 10.98 | -0.7 |
| CHLRE_07g315150v5 | 3.24 | -0.7 |
| CHLRE_02g088850v5 | 2.39 | -0.7 |
| CHLRE_12g483850v5 | 17.27 | -0.69 |
| CHLRE_01g017300v5 | 11.59 | -0.69 |
| CHLRE_12g494550v5 | 9.2 | -0.69 |
| CHLRE_14g611300v5 | 7.47 | -0.69 |
| CHLRE_12g495550v5 | 3.18 | -0.69 |
| CHLRE_12g510050v5 | 2.31 | -0.69 |
| CHLRE_16g679500v5 | 4.75 | -0.68 |
| CHLRE_01g018650v5 | 3.52 | -0.68 |
| CHLRE_16g656950v5 | 2.37 | -0.68 |
| CHLRE_02g143550v5 | 2.06 | -0.68 |
| CHLRE_10g440400v5 | 2.06 | -0.68 |
| CHLRE_01g062172v5 | 12.61 | -0.67 |
| CHLRE_12g519950v5 | 2.09 | -0.67 |
| CHLRE_06g300800v5 | 10.01 | -0.66 |
| CHLRE_01g026600v5 | 6.51 | -0.66 |
| CHLRE_03g206600v5 | 5.31 | -0.65 |
| CHLRE_12g507300v5 | 17.83 | -0.64 |
| CHLRE_09g394750v5 | 5.21 | -0.64 |
| CHLRE_13g576400v5 | 3.32 | -0.64 |
| CHLRE_11g480150v5 | 10.02 | -0.63 |
| CHLRE_14g612450v5 | 6.43 | -0.63 |
| CHLRE_06g275350v5 | 2.6 | -0.63 |
| CHLRE_03g195650v5 | 6.28 | -0.62 |
| CHLRE_02g096150v5 | 4.06 | -0.62 |
| CHLRE_16g651923v5 | 4.03 | -0.62 |
| CHLRE_06g278213v5 | 3.9 | -0.62 |
| CHLRE_10g427700v5 | 2.88 | -0.62 |
| CHLRE_03g160400v5 | 2.21 | -0.62 |
| CHLRE_12g528000v5 | 8.38 | -0.61 |
| CHLRE_06g285401v5 | 4.58 | -0.61 |
| CHLRE_15g635600v5 | 11.33 | -0.6 |
| CHLRE_09g412100v5 | 5.47 | -0.6 |
| CHLRE_04g226850v5 | 3.33 | -0.6 |
| CHLRE_06g250100v5 | 2.7 | -0.59 |
| CHLRE_17g727100v5 | 2.08 | -0.59 |
| CHLRE_16g683793v5 | 2.04 | -0.59 |
| CHLRE_03g190850v5 | 3.35 | -0.58 |
| CHLRE_09g395350v5 | 2.3 | -0.58 |
| CHLRE_07g346200v5 | 2.28 | -0.58 |

<sup>a</sup>Gramene (https://www.gramene.org/)

**Supplementary Table 3.** Yeast genes whose expression levels increased due to co-cultivation of microalgae and yeast.

| Gene<sup>a</sup> or locus tag<sup>b</sup> | -Log<sub>10</sub> (p-value) | Log<sub>2</sub> (fold change) |
| --- | --- | --- |
| YAR010C | 5.5 | 6.94 |
| YPR204W | 2.33 | 3.99 |
| YNCD0008W | 4.31 | 3.15 |
| YNCL0037W | 3.96 | 2.86 |
| YNCN0010C | 3.03 | 2.69 |
| PAU22 | 4.73 | 2.69 |
| FIT3 | 7.78 | 2.27 |
| YNCM0020W | 3.09 | 2.15 |
| MPH3 | 5.54 | 1.9 |
| DCG1 | 5.75 | 1.79 |
| YNCM0019W | 3.26 | 1.69 |
| YOR381W-A | 4.52 | 1.66 |
| YNCB0006W | 4.2 | 1.63 |
| YOL107W | 7.89 | 1.62 |
| YNCE0021C | 4.66 | 1.58 |
| HOT13 | 6.23 | 1.57 |
| YNCJ0014C | 2.21 | 1.44 |
| MIT1 | 12.03 | 1.37 |
| DUR3 | 17.77 | 1.37 |
| YKR104W | 2.05 | 1.36 |
| ELP6 | 8.15 | 1.34 |
| BSC5 | 5.08 | 1.26 |
| KEI1 | 13.54 | 1.26 |
| ARC18 | 19.12 | 1.24 |
| YGR035W-A | 10.3 | 1.23 |
| YNCK0016W | 10.9 | 1.23 |
| ATG41 | 13.19 | 1.22 |
| PBA1 | 13.84 | 1.22 |
| YNCK0005C | 2.86 | 1.21 |
| PCL1 | 4.48 | 1.2 |
| YMR247W-A | 2.14 | 1.16 |
| COQ9 | 9.97 | 1.12 |
| ATP8 | 5.98 | 1.09 |
| TKL2 | 11.69 | 1.09 |
| GAT1 | 4.91 | 1.08 |
| ARC19 | 17.17 | 1.06 |
| MF(ALPHA)2 | 4.38 | 1.05 |
| SDS22 | 8.73 | 1.05 |
| AGA2 | 3.72 | 1.04 |
| DIA4 | 5.52 | 1.03 |
| ACP1 | 6.4 | 1.03 |
| SPL2 | 14.32 | 1.03 |
| YNR068C | 2.8 | 1.02 |
| COS4 | 2.58 | 1.01 |
| YEL068C | 5.55 | 1 |
| AIM34 | 2.2 | 0.99 |
| MUD1 | 8.97 | 0.99 |
| YLR126C | 10.18 | 0.99 |
| SST2 | 3.43 | 0.98 |
| IME1 | 8.29 | 0.97 |
| SWI1 | 15.84 | 0.96 |
| COS6 | 6.83 | 0.95 |
| APA1 | 7.93 | 0.95 |
| YPR063C | 9.08 | 0.95 |
| LUG1 | 3.92 | 0.94 |
| ATG33 | 10.07 | 0.94 |
| YHM2 | 8.22 | 0.93 |
| PUP1 | 13.06 | 0.93 |
| OPI10 | 2.94 | 0.92 |
| PHS1 | 6.01 | 0.92 |
| DAL7 | 18.69 | 0.92 |
| HAL1 | 5.65 | 0.91 |
| DPM1 | 6.76 | 0.91 |
| EMC10 | 14.55 | 0.91 |
| YNCL0009C | 2.85 | 0.9 |
| GPI2 | 4.62 | 0.9 |
| PIB2 | 8.38 | 0.9 |
| TAF5 | 9.22 | 0.9 |
| ECM3 | 11.29 | 0.9 |
| VPS28 | 5.2 | 0.89 |
| MOB2 | 4.02 | 0.88 |
| MPT5 | 7.11 | 0.88 |
| LEU4 | 22.01 | 0.87 |
| SNX41 | 2.06 | 0.86 |
| YLR342W-A | 12.28 | 0.86 |
| ILM1 | 12.6 | 0.86 |
| ERG28 | 12.61 | 0.86 |
| ATF1 | 12.75 | 0.86 |
| ZNF1 | 6.03 | 0.85 |
| YPL229W | 7.15 | 0.85 |
| TVP15 | 9.1 | 0.85 |
| EAF3 | 17.76 | 0.85 |
| YPR145C-A | 19.95 | 0.85 |
| APT2 | 23.94 | 0.85 |
| BUD9 | 2.88 | 0.84 |
| PRO3 | 8.25 | 0.84 |
| ATP1 | 22.52 | 0.84 |
| CIT3 | 11.55 | 0.83 |
| YIP1 | 15.58 | 0.83 |
| PKP2 | 5.5 | 0.82 |
| YPR053C | 5.53 | 0.82 |
| VPS34 | 6.55 | 0.82 |
| RNH203 | 6.65 | 0.82 |
| SWC5 | 10.18 | 0.82 |
| YFH1 | 12.09 | 0.82 |
| ATG40 | 20.96 | 0.82 |
| SUT390 | 2.14 | 0.81 |
| YBR196C-A | 2.63 | 0.81 |
| PHB1 | 15.67 | 0.81 |
| BNA3 | 3.04 | 0.8 |
| YMC2 | 4.73 | 0.8 |
| AIM17 | 5.82 | 0.8 |
| UBC7 | 12.97 | 0.8 |
| GUT2 | 24.74 | 0.8 |
| SNR6 | 2.42 | 0.79 |
| CSM1 | 2.91 | 0.79 |
| NNR2 | 3.77 | 0.79 |
| ICP55 | 3.82 | 0.79 |
| SCM4 | 4.13 | 0.79 |
| TOS7 | 4.77 | 0.79 |
| YFL051C | 4.81 | 0.79 |
| BSC6 | 7.44 | 0.79 |
| GIS1 | 13.59 | 0.79 |
| SNC1 | 15.41 | 0.79 |
| PHO80 | 8.18 | 0.78 |
| NHP6A | 12.9 | 0.78 |
| CAT8 | 16.85 | 0.78 |
| PDB1 | 19.44 | 0.78 |
| UBC9 | 19.62 | 0.78 |
| GPD1 | 21.36 | 0.78 |
| ECM11 | 2.79 | 0.77 |
| MET17 | 3.48 | 0.77 |
| TOM20 | 11.18 | 0.77 |
| YDL012C | 3.99 | 0.76 |
| SEC22 | 4.74 | 0.76 |
| YLL056C | 5.11 | 0.76 |
| PMT6 | 6.16 | 0.76 |
| YMR105W-A | 7.02 | 0.76 |
| YNL134C | 9.94 | 0.76 |
| GID10 | 10.1 | 0.76 |
| ATP5 | 13.92 | 0.76 |
| GAB1 | 4.65 | 0.75 |
| YMC1 | 5.17 | 0.75 |
| RNH1 | 5.68 | 0.75 |
| ARG7 | 7.83 | 0.75 |
| POL30 | 8.71 | 0.75 |
| YNR064C | 11.98 | 0.75 |
| TOS2 | 4.2 | 0.74 |
| YMR181C | 5.73 | 0.74 |
| RFA3 | 9.53 | 0.74 |
| TRX1 | 14.08 | 0.74 |
| MRX18 | 14.19 | 0.74 |
| SNU13 | 20.41 | 0.74 |
| DMC1 | 23.16 | 0.74 |
| YVC1 | 2.17 | 0.73 |
| YHR173C | 2.19 | 0.73 |
| BIO2 | 2.26 | 0.73 |
| GCR1 | 2.42 | 0.73 |
| MVP1 | 4.98 | 0.73 |
| FRE3 | 8.22 | 0.73 |
| AUA1 | 13.79 | 0.73 |
| SLM2 | 2.27 | 0.72 |
| YER181C | 2.61 | 0.72 |
| PBP2 | 3.38 | 0.72 |
| MCP1 | 3.55 | 0.72 |
| CGI121 | 3.64 | 0.72 |
| YML054C-A | 3.65 | 0.72 |
| YFR012W-A | 4.28 | 0.72 |
| YSP3 | 4.87 | 0.72 |
| ULP1 | 7.94 | 0.72 |
| ORM1 | 13.41 | 0.72 |
| YBR071W | 13.44 | 0.72 |
| GGA1 | 2.9 | 0.71 |
| ODC2 | 5.1 | 0.71 |
| YNL034W | 5.63 | 0.71 |
| TCB1 | 6.42 | 0.71 |
| ERV15 | 7.8 | 0.71 |
| ISC10 | 9.52 | 0.71 |
| MFA1 | 9.97 | 0.71 |
| MCK1 | 10.45 | 0.71 |
| TDA10 | 3.56 | 0.7 |
| TPS1 | 3.82 | 0.7 |
| RMR1 | 5.71 | 0.7 |
| VHR1 | 6.3 | 0.7 |
| AVT1 | 6.68 | 0.7 |
| MSC2 | 8.5 | 0.7 |
| VPS36 | 9.35 | 0.7 |
| YNL190W | 21.34 | 0.7 |
| YJR079W | 2.58 | 0.69 |
| YIP5 | 2.62 | 0.69 |
| YNL018C | 3.67 | 0.69 |
| NRS1 | 3.97 | 0.69 |
| SLC1 | 4.03 | 0.69 |
| MET28 | 4.08 | 0.69 |
| YPT53 | 20.34 | 0.69 |
| PCF11 | 2.52 | 0.68 |
| ARD1 | 2.95 | 0.68 |
| BDH2 | 3.15 | 0.68 |
| TES1 | 4.63 | 0.68 |
| SUL1 | 7.96 | 0.68 |
| GPB2 | 8.85 | 0.68 |
| GID11 | 17.21 | 0.68 |
| MIN6 | 20.93 | 0.68 |
| GSF2 | 22.55 | 0.68 |
| CIN5 | 2.88 | 0.67 |
| YLR285C-A | 3.6 | 0.67 |
| AIM7 | 4.17 | 0.67 |
| MDS3 | 5.17 | 0.67 |
| ARO80 | 5.3 | 0.67 |
| GRX3 | 6.27 | 0.67 |
| YAK1 | 7.45 | 0.67 |
| COA6 | 9.25 | 0.67 |
| YPF1 | 12.62 | 0.67 |
| CMI7 | 15.22 | 0.67 |
| CAT2 | 20.39 | 0.67 |
| MFA2 | 2.51 | 0.66 |
| YKU80 | 3.6 | 0.66 |
| CKI1 | 3.83 | 0.66 |
| SNG1 | 3.94 | 0.66 |
| SGN1 | 4.81 | 0.66 |
| YIF1 | 7.24 | 0.66 |
| ATG2 | 7.95 | 0.66 |
| NCA3 | 10.91 | 0.66 |
| SNR80 | 16.01 | 0.66 |
| ITS2-1 | 25.18 | 0.66 |
| SYM1 | 2.05 | 0.65 |
| HXT8 | 2.66 | 0.65 |
| YJL070C | 2.72 | 0.65 |
| DSE1 | 3.43 | 0.65 |
| MCT1 | 4.48 | 0.65 |
| IMG1 | 6.48 | 0.65 |
| HMG1 | 9.69 | 0.65 |
| DFP2 | 10.5 | 0.65 |
| PCL5 | 11.8 | 0.65 |
| ADR1 | 14.26 | 0.65 |
| ITS2-2 | 24.75 | 0.65 |
| RRT13 | 2.11 | 0.64 |
| GAS3 | 2.7 | 0.64 |
| AIM18 | 3.28 | 0.64 |
| PMA2 | 3.61 | 0.64 |
| SEC11 | 7.4 | 0.64 |
| NGR1 | 9.26 | 0.64 |
| WWM1 | 10.36 | 0.64 |
| GVP36 | 11.91 | 0.64 |
| MSS18 | 3.45 | 0.63 |
| GPM2 | 4.34 | 0.63 |
| MRX19 | 4.54 | 0.63 |
| IRC24 | 7.17 | 0.63 |
| TED1 | 9.42 | 0.63 |
| PEX18 | 10.92 | 0.63 |
| ORA1 | 14.76 | 0.63 |
| LSC1 | 15.06 | 0.63 |
| BLI1 | 16.01 | 0.63 |
| YHR175W-A | 19.3 | 0.63 |
| YCP4 | 21.37 | 0.63 |
| YGR237C | 2.12 | 0.62 |
| YCR099C | 2.39 | 0.62 |
| YGR117C | 2.4 | 0.62 |
| HAP2 | 2.73 | 0.62 |
| FRE5 | 2.96 | 0.62 |
| CAP2 | 4.87 | 0.62 |
| YPR1 | 5.48 | 0.62 |
| PEX13 | 6.45 | 0.62 |
| SNN1 | 7.02 | 0.62 |
| PST2 | 7.58 | 0.62 |
| RNY1 | 8.16 | 0.62 |
| ATP16 | 8.5 | 0.62 |
| AIM6 | 11.66 | 0.62 |
| UGX2 | 14.14 | 0.62 |
| AMS1 | 23.66 | 0.62 |
| YLR410W-A | 2.34 | 0.61 |
| AIM11 | 3.04 | 0.61 |
| CYB2 | 3.24 | 0.61 |
| ANK1 | 3.31 | 0.61 |
| MET5 | 3.77 | 0.61 |
| ECM29 | 4.26 | 0.61 |
| SBH1 | 6.7 | 0.61 |
| HED1 | 7.54 | 0.61 |
| PAT1 | 8.15 | 0.61 |
| IZH1 | 8.26 | 0.61 |
| BOL1 | 9.68 | 0.61 |
| RVS167 | 12.83 | 0.61 |
| SMT3 | 14.82 | 0.61 |
| COI1 | 15.52 | 0.61 |
| ATG42 | 23.27 | 0.61 |
| YNCE0026W | 2.01 | 0.6 |
| PAU2 | 2.05 | 0.6 |
| GPH1 | 2.66 | 0.6 |
| RUF21 | 3 | 0.6 |
| MDV1 | 3.33 | 0.6 |
| MEF2 | 3.8 | 0.6 |
| MIM2 | 3.91 | 0.6 |
| RBL2 | 4.27 | 0.6 |
| HOP1 | 5.47 | 0.6 |
| YDL199C | 8.91 | 0.6 |
| AIM24 | 2.44 | 0.59 |
| MTC5 | 3.66 | 0.59 |
| YJR096W | 7.2 | 0.59 |
| YRA2 | 7.36 | 0.59 |
| TFS1 | 7.63 | 0.59 |

<sup>a</sup>UniProt (https://www.uniprot.org/)
<sup>b</sup>*Saccharomyces* Genome Database (https://www.yeastgenome.org/)

**Supplementary Table 4.** Yeast genes whose expression levels decreased due to co-cultivation of microalgae and yeast.

| Gene<sup>a</sup> or locus tag<sup>b</sup> | -Log<sub>10</sub> (p-value) | Log<sub>2</sub> (fold change) |
| --- | --- | --- |
| YHR214C-E | 4.85 | -5.25 |
| YNCJ0012W | 5.33 | -4.31 |
| SUP53 | 6.56 | -4.22 |
| SUF8 | 6.88 | -4.11 |
| EMT1 | 10.93 | -3.56 |
| YNCP0017C | 2.6 | -3.12 |
| TRR4 | 7.14 | -3.12 |
| YNCL0007W | 13 | -3.07 |
| YNCD0024C | 4.64 | -3.01 |
| YNCD0004W | 6.41 | -3.01 |
| YNCI0009W | 2.74 | -2.9 |
| YNCM0023C | 7.32 | -2.82 |
| YNCF0009C | 6.31 | -2.7 |
| YNCP0008C | 8.65 | -2.69 |
| YNCN0008C | 5.78 | -2.63 |
| YNCL0002C | 2.61 | -2.4 |
| SUP51 | 5.27 | -2.4 |
| TRN1 | 2.16 | -2.31 |
| YNCG0027W | 2.82 | -2.31 |
| YNCH0002C | 2.86 | -2.31 |
| YOR389W | 18.72 | -2.25 |
| YNCB0012W | 2.89 | -2.19 |
| YNCG0037W | 3.69 | -2.19 |
| YNCM0025C | 2.22 | -2.12 |
| YNCG0029C | 2.12 | -2.05 |
| YNCG0032W | 4.64 | -1.99 |
| YIL054W | 5.75 | -1.99 |
| YNCJ0008W | 10.67 | -1.97 |
| YNCP0023W | 2.75 | -1.84 |
| YNL067W-B | 6.21 | -1.84 |
| RGD2 | 10.13 | -1.84 |
| YPL277C | 11.82 | -1.82 |
| YNCG0042C | 2.75 | -1.81 |
| SOE1 | 4.36 | -1.81 |
| YNCO0011W | 4.93 | -1.76 |
| YNCG0011W | 3.86 | -1.73 |
| tK(UUU)Q | 20.67 | -1.72 |
| SUF23 | 2.81 | -1.69 |
| YNCP0001C | 8.43 | -1.69 |
| NAT4 | 14.85 | -1.65 |
| SUP54 | 5.18 | -1.63 |
| YNCG0033W | 4.53 | -1.6 |
| YLR365W | 4.12 | -1.57 |
| POG1 | 20.19 | -1.56 |
| YNCO0008W | 2.01 | -1.53 |
| GIS4 | 17.22 | -1.52 |
| SUP61 | 2.38 | -1.5 |
| YNCM0033C | 4.85 | -1.5 |
| ASF1 | 19.85 | -1.5 |
| SNR50 | 25.61 | -1.5 |
| EFM6 | 20.58 | -1.49 |
| AQY2 | 20.98 | -1.49 |
| CDC65 | 2.15 | -1.45 |
| ECM19 | 9.16 | -1.44 |
| PIG2 | 10.51 | -1.42 |
| tP(UGG)Q | 10.19 | -1.41 |
| YNCD0017W | 3.03 | -1.36 |
| YNCD0019C | 4.27 | -1.36 |
| VHR2 | 4.89 | -1.36 |
| YNCG0023C | 23.56 | -1.36 |
| HXT3 | 7.23 | -1.35 |
| ZRG8 | 9.55 | -1.33 |
| BUR2 | 16.06 | -1.3 |
| ZDS1 | 4.52 | -1.28 |
| tY(GUA)Q | 5.28 | -1.28 |
| YPR064W | 2.75 | -1.27 |
| YJL107C | 8.13 | -1.27 |
| ATF2 | 8.92 | -1.24 |
| YJL027C | 10.67 | -1.24 |
| ANS1 | 10.67 | -1.23 |
| SSK22 | 13.11 | -1.23 |
| SRV2 | 2.59 | -1.22 |
| YER158C | 21.87 | -1.22 |
| SRL3 | 20.9 | -1.21 |
| BNR1 | 4.55 | -1.2 |
| YNCC0005W | 10.88 | -1.2 |
| YOR008C-A | 23.59 | -1.2 |
| MNL1 | 7.56 | -1.18 |
| tN(GUU)Q | 4.85 | -1.17 |
| AQR1 | 6.21 | -1.17 |
| SNR71 | 24.27 | -1.17 |
| SNF1 | 10.54 | -1.16 |
| YAH1 | 2.92 | -1.15 |
| IRT1 | 16.71 | -1.14 |
| YMR082C | 3.3 | -1.12 |
| YNCK0021W | 4.23 | -1.11 |
| TAT2 | 14.71 | -1.11 |
| HTC1 | 3.83 | -1.1 |
| FLX1 | 6.48 | -1.09 |
| SNR66 | 19.54 | -1.09 |
| HLR1 | 4.19 | -1.08 |
| SRX1 | 4.57 | -1.08 |
| UPA1 | 24.83 | -1.07 |
| RDN5-1 | 6.56 | -1.06 |
| YNCG0028W | 2.57 | -1.05 |
| SNR53 | 8.69 | -1.05 |
| BUD7 | 11.05 | -1.03 |
| YKR041W | 4.97 | -1.02 |
| YPL245W | 6.76 | -1.02 |
| COM2 | 18.14 | -1.02 |
| ADE17 | 27.21 | -1.02 |
| PYC2 | 15.44 | -1.01 |
| YLR406C-A | 4.31 | -1 |
| AST1 | 5.51 | -1 |
| OSW1 | 10.34 | -1 |
| YNCE0028C | 2.51 | -0.99 |
| SIZ1 | 3.19 | -0.99 |
| MTD1 | 25.66 | -0.99 |
| FKH2 | 12.19 | -0.98 |
| SED1 | 24.16 | -0.98 |
| YPL068C | 4.61 | -0.97 |
| PRY1 | 2.1 | -0.96 |
| SPT10 | 7.06 | -0.96 |
| DIG1 | 9.39 | -0.96 |
| TUP1 | 2.07 | -0.95 |
| HXT13 | 5.78 | -0.95 |
| CTH1 | 9.8 | -0.95 |
| AIR1 | 16.21 | -0.95 |
| ACK1 | 7.14 | -0.94 |
| CAB3 | 10.38 | -0.93 |
| YML037C | 3.2 | -0.92 |
| YNCD0023C | 12.66 | -0.92 |
| ENT4 | 21.71 | -0.92 |
| PRM10 | 4.63 | -0.91 |
| KNH1 | 10.39 | -0.91 |
| MTR4 | 26.72 | -0.91 |
| RAX1 | 5.87 | -0.9 |
| YGL235W | 6.68 | -0.9 |
| SNR65 | 14.93 | -0.9 |
| YGL230C | 16.69 | -0.9 |
| HSU1 | 2.29 | -0.89 |
| CLN3 | 2.81 | -0.89 |
| UPS3 | 7.86 | -0.89 |
| EST1 | 11.3 | -0.89 |
| GCV1 | 25.36 | -0.89 |
| CHA1 | 3.51 | -0.88 |
| YGR121W-A | 3.76 | -0.88 |
| RBS1 | 3.77 | -0.88 |
| PDR12 | 8.97 | -0.88 |
| RAD7 | 21.6 | -0.88 |
| SPR3 | 21.95 | -0.88 |
| CMR1 | 2.68 | -0.87 |
| ASG7 | 2.81 | -0.87 |
| YOR293C-A | 3.84 | -0.87 |
| TIM50 | 5.8 | -0.87 |
| RGS2 | 17.64 | -0.87 |
| SPS1 | 26.89 | -0.87 |
| GAT3 | 7.14 | -0.86 |
| MAE1 | 13.91 | -0.86 |
| RTT10 | 20.61 | -0.86 |
| PCA1 | 21.26 | -0.86 |
| YLR255C | 6.44 | -0.85 |
| SAM4 | 3.48 | -0.84 |
| YNL095C | 6.88 | -0.84 |
| SGA1 | 11.09 | -0.84 |
| PRM7 | 15.23 | -0.84 |
| DUS3 | 20.46 | -0.84 |
| YLR225C | 9.51 | -0.83 |
| TPK3 | 10.17 | -0.83 |
| LCP5 | 14.83 | -0.83 |
| TCD2 | 17.54 | -0.83 |
| PRK1 | 3.1 | -0.82 |
| YRO2 | 3.8 | -0.82 |
| GRC3 | 6.81 | -0.82 |
| PMA1 | 14.52 | -0.81 |
| YEL023C | 15 | -0.81 |
| BZZ1 | 3.91 | -0.8 |
| YHL026C | 5.68 | -0.8 |
| PES4 | 11.66 | -0.8 |
| USV1 | 3.07 | -0.79 |
| DSN1 | 3.2 | -0.79 |
| SPC29 | 3.32 | -0.79 |
| DOA1 | 4.56 | -0.79 |
| YPR027C | 7.81 | -0.79 |
| SDS23 | 9.93 | -0.79 |
| UTP18 | 16.59 | -0.79 |
| SRP40 | 25.45 | -0.79 |
| SMA2 | 5.32 | -0.78 |
| YGR054W | 7.78 | -0.78 |
| POP3 | 9.18 | -0.78 |
| ARN1 | 23.44 | -0.78 |
| YNR014W | 2.25 | -0.77 |
| MSA1 | 2.73 | -0.77 |
| MYO4 | 5.62 | -0.77 |
| ART5 | 5.96 | -0.77 |
| RTC4 | 6.27 | -0.77 |
| PSK2 | 11.72 | -0.77 |
| CBF2 | 12.35 | -0.77 |
| RRT12 | 13.77 | -0.77 |
| SHQ1 | 14.93 | -0.77 |
| UBP7 | 2.59 | -0.76 |
| IFM1 | 3.18 | -0.76 |
| MRS3 | 6.46 | -0.76 |
| DIN7 | 8.06 | -0.76 |
| DEG1 | 13.12 | -0.76 |
| SPR28 | 13.12 | -0.76 |
| SPO19 | 20.66 | -0.76 |
| CLB4 | 3.85 | -0.75 |
| LAF1 | 6.55 | -0.75 |
| YPK2 | 11.58 | -0.75 |
| BAG7 | 25.54 | -0.75 |
| SCM3 | 2.82 | -0.74 |
| YLR179C | 6.54 | -0.74 |
| FRE8 | 6.76 | -0.74 |
| LSG1 | 7.6 | -0.74 |
| YMR315W-A | 7.9 | -0.74 |
| GAT4 | 8.02 | -0.74 |
| NDE1 | 14.28 | -0.74 |
| GFA1 | 28.81 | -0.74 |
| RBH1 | 2.58 | -0.73 |
| PGD1 | 3.16 | -0.73 |
| YPR096C | 3.28 | -0.73 |
| AIM4 | 4.93 | -0.73 |
| UBA3 | 7.9 | -0.73 |
| GAS2 | 8.2 | -0.73 |
| NAB2 | 21.7 | -0.73 |
| NEJ1 | 2.23 | -0.72 |
| YDL211C | 3.04 | -0.72 |
| YNCD0007C | 4.6 | -0.72 |
| ALY2 | 4.7 | -0.72 |
| ADD37 | 7.24 | -0.72 |
| tM(CAU)Q1 | 8.69 | -0.72 |
| OCA5 | 16.63 | -0.72 |
| SOK2 | 18.35 | -0.72 |
| BCK2 | 23.64 | -0.72 |
| RPL17B | 29.2 | -0.72 |
| SRS2 | 3.04 | -0.71 |
| SEC6 | 3.49 | -0.71 |
| AAR2 | 3.5 | -0.71 |
| TAX4 | 3.71 | -0.71 |
| CBP1 | 4.59 | -0.71 |
| SGO1 | 4.8 | -0.71 |
| PKH3 | 5.66 | -0.71 |
| MCM10 | 5.88 | -0.71 |
| SER2 | 6.18 | -0.71 |
| RRG1 | 6.81 | -0.71 |
| PBS2 | 7.44 | -0.71 |
| YKL106C-A | 8.07 | -0.71 |
| SNR55 | 16.5 | -0.71 |
| YOR342C | 18.9 | -0.71 |
| SSF1 | 20.49 | -0.71 |
| THR4 | 5.12 | -0.7 |
| YLR162W | 5.55 | -0.7 |
| NDT80 | 7.95 | -0.7 |
| CDA1 | 8.77 | -0.7 |
| SKT5 | 9.79 | -0.7 |
| SRG1 | 17.48 | -0.7 |
| UME6 | 3.15 | -0.69 |
| RRM3 | 3.71 | -0.69 |
| TRF5 | 8.35 | -0.69 |
| RPL6B | 26.88 | -0.69 |
| PAC1 | 2.11 | -0.68 |
| EDC1 | 5 | -0.68 |
| SIR3 | 6.06 | -0.68 |
| BUL2 | 6.79 | -0.68 |
| YGR273C | 8.44 | -0.68 |
| NUP60 | 9.26 | -0.68 |
| YJR107C-A | 9.78 | -0.68 |
| ATG15 | 13.82 | -0.68 |
| UBC11 | 15.62 | -0.68 |
| AQY1 | 16.43 | -0.68 |
| SAP155 | 16.54 | -0.68 |
| AFT1 | 2.81 | -0.67 |
| TAT1 | 2.92 | -0.67 |
| GAL7 | 3.15 | -0.67 |
| ADA2 | 3.92 | -0.67 |
| PET309 | 4.03 | -0.67 |
| SAE2 | 4.57 | -0.67 |
| PNS1 | 5.21 | -0.67 |
| UPS2 | 7.62 | -0.67 |
| HXT10 | 9.56 | -0.67 |
| GCN3 | 10.88 | -0.67 |
| PWP2 | 26.23 | -0.67 |
| MAM1 | 3.37 | -0.66 |
| GDE1 | 5.21 | -0.66 |
| SRD1 | 5.63 | -0.66 |
| PPR1 | 5.84 | -0.66 |
| DON1 | 5.95 | -0.66 |
| ALD5 | 6.05 | -0.66 |
| tS(UGA)Q2 | 6.16 | -0.66 |
| SOG2 | 7.09 | -0.66 |
| NUT2 | 9.35 | -0.66 |
| UTP30 | 13.29 | -0.66 |
| TRX3 | 14.39 | -0.66 |
| SPR6 | 23.65 | -0.66 |
| FLO8 | 2.46 | -0.65 |
| TBRT | 2.58 | -0.65 |
| ESP1 | 2.81 | -0.65 |
| MSL5 | 3.48 | -0.65 |
| YCL021W-A | 3.49 | -0.65 |
| YOL097W-A | 3.92 | -0.65 |
| BIL2 | 4.46 | -0.65 |
| REX2 | 5.95 | -0.65 |
| KCH1 | 6.47 | -0.65 |
| SWT21 | 6.78 | -0.65 |
| SPS22 | 9.05 | -0.65 |
| SXM1 | 18.78 | -0.65 |
| ANY1 | 19.67 | -0.65 |
| CHS3 | 19.87 | -0.65 |
| GPI18 | 24.13 | -0.65 |
| TPC1 | 2.7 | -0.64 |
| APC5 | 3.04 | -0.64 |
| SNF5 | 3.16 | -0.64 |
| YPL039W | 3.93 | -0.64 |
| YER085C | 4.58 | -0.64 |
| DUS4 | 6.37 | -0.64 |
| EMI2 | 11.08 | -0.64 |
| TMA46 | 12.29 | -0.64 |
| KRI1 | 16.59 | -0.64 |
| UTP9 | 20.57 | -0.64 |
| SPB1 | 25.03 | -0.64 |
| MED6 | 3.82 | -0.63 |
| ECM23 | 5.34 | -0.63 |
| APP1 | 5.86 | -0.63 |
| MIP6 | 6.06 | -0.63 |
| YIL067C | 6.89 | -0.63 |
| FRS2 | 7.41 | -0.63 |
| HXT14 | 7.73 | -0.63 |
| PRM5 | 25.13 | -0.63 |
| RPL7A | 27.1 | -0.63 |
| MAK10 | 2.37 | -0.62 |
| MRX11 | 2.6 | -0.62 |
| YBR182C-A | 2.71 | -0.62 |
| GEM1 | 2.94 | -0.62 |
| YDR541C | 3.16 | -0.62 |
| YML6 | 3.17 | -0.62 |
| YJL043W | 3.28 | -0.62 |
| PXL1 | 3.94 | -0.62 |
| YNL143C | 4.05 | -0.62 |
| UBP3 | 5.12 | -0.62 |
| NSE5 | 5.43 | -0.62 |
| DPB4 | 5.55 | -0.62 |
| ATP22 | 8.66 | -0.62 |
| PRP43 | 22.38 | -0.62 |
| GIP3 | 26.83 | -0.62 |
| RPS15 | 29.83 | -0.62 |
| PRP9 | 2.95 | -0.61 |
| PFS2 | 3.73 | -0.61 |
| SMK1 | 4.7 | -0.61 |
| MLC1 | 4.92 | -0.61 |
| MTO1 | 4.92 | -0.61 |
| ASK1 | 7.95 | -0.61 |
| RAP1 | 8.47 | -0.61 |
| SMP1 | 8.98 | -0.61 |
| NUP133 | 9.09 | -0.61 |
| PXR1 | 10.51 | -0.61 |
| PSR2 | 12.43 | -0.61 |
| RIX1 | 12.94 | -0.61 |
| UTP4 | 18.35 | -0.61 |
| PDE2 | 30.55 | -0.61 |
| GCG1 | 2.95 | -0.6 |
| YGL138C | 3.29 | -0.6 |
| SED5 | 3.29 | -0.6 |
| DUG3 | 4.29 | -0.6 |
| THR1 | 4.61 | -0.6 |
| MRL1 | 4.83 | -0.6 |
| SRL4 | 5.37 | -0.6 |
| SCS3 | 5.69 | -0.6 |
| YGL149W | 5.9 | -0.6 |
| YSP2 | 8.49 | -0.6 |
| SLD7 | 9 | -0.6 |
| PAM18 | 10.94 | -0.6 |
| CDC3 | 11.14 | -0.6 |
| FPK1 | 13.4 | -0.6 |
| RIA1 | 14.07 | -0.6 |
| DBP3 | 22.06 | -0.6 |
| URA7 | 30.74 | -0.6 |
| PTP1 | 2.86 | -0.59 |
| EPO1 | 6.11 | -0.59 |
| CTS2 | 6.85 | -0.59 |
| PMD1 | 8.73 | -0.59 |
| SNF6 | 9.54 | -0.59 |
| PDC6 | 16.06 | -0.59 |
| DED81 | 16.72 | -0.59 |
| RIM4 | 27.45 | -0.59 |

<sup>a</sup>UniProt (https://www.uniprot.org/)
<sup>b</sup>Saccharomyces Genome Database (https://www.yeastgenome.org/)

